# Elevated MPS1 converts the fibrous corona into a source of chromosomal instability in colon cancer

**DOI:** 10.64898/2026.08.07.743489

**Authors:** Ana Pinto-Teixeira, André Oliveira, Gonçalo Gouveia, Ana Sousa, Jessica Roelands, Marco Grillo, Sergi Rodriguez-Calado, Carlos Resende, Helena Xavier-Ferreira, Mariana Osswald, Hugo Girão, Patrícia Mesquita, Raquel Almeida, Marin Barisic, José Carlos Machado, Claudio Sunkel, Tom van Wezel, Daniele Fachinetti, Karoly Szuhai, Noel F.C.C. de Miranda, Carlos Conde

**Affiliations:** i3S, Instituto de Investigação e Inovação em Saúde, Universidade do Porto, Rua Alfredo Allen 208, 4200-135, Porto, Portugal; IBMC, Instituto de Biologia Molecular e Celular, Universidade do Porto, Rua Alfredo Allen 208, 4200-135, Porto, Portugal; Programa Doutoral em Biologia Molecular e Celular (MCbiology), Instituto de Ciências Biomédicas Abel Salazar (ICBAS), Universidade do Porto, Rua Jorge Viterbo Ferreira 228, 4050-313 Porto, Portugal; Department of Pathology, Leiden University Medical Center, Leiden, The Netherlands; Institut Curie, PSL Research University, CNRS, UMR 144 and 3664, F-75005 Paris, France; Cell Division and Cytoskeleton, Danish Cancer Institute, Strandboulevarden 49, DK-2100 Copenhagen, Denmark; IPATIMUP, Instituto de Patologia e Imunologia Molecular da Universidade do Porto, Rua Júlio Amaral de Carvalho 45, 4200-135 Porto; Department of Cellular and Molecular Medicine, University of Copenhagen, Blegdamsvej 3B, 2200 Copenhagen N, Denmark; Departamento de Biologia Molecular, Instituto de Ciências Biomédicas Abel Salazar (ICBAS), Universidade do Porto, Rua Jorge Viterbo Ferreira 228, 4050-313 Porto, Portugal; Department of Cell and Chemical Biology, Leiden University Medical Center, Leiden, The Netherlands

**Keywords:** CIN, Colon Cancer, MPS1, Fibrous corona, Merotely

## Abstract

Chromosomal instability (CIN), characterised by recurrent chromosome mis-segregation, fuels intratumour heterogeneity and tumour evolution. Yet, its proximal molecular causes remain ill-defined. Tumour-scale genomic and transcriptomic analyses associate MPS1 overexpression with CIN in colon carcinomas. Consistently, CIN^+^ patient-derived colon cancer cells (PCCCs) display elevated MPS1, frequent merotelic kinetochore-microtubule attachments and lagging chromosomes that are suppressed by partial MPS1 inhibition. Conversely, ectopic MPS1 overexpression in otherwise stable near-diploid cells phenocopies these defects, inducing micronuclei formation and karyotypic divergence. Mechanistically, MPS1 overactivity maintains ROD phosphorylation at Thr13/Ser15, continuously sustaining fibrous corona assembly despite mature end-on attachments. 3D-STED microscopy reveals split-interface configurations in which microtubules from opposing poles engage the canonical outer-kinetochore surface and a spatially distinct corona domain, explaining how merotely forms and persists into anaphase. Disrupting corona assembly restores chromosome segregation fidelity in MPS1-overexpressing RPE-1 cells and in CIN^+^ PCCCs. Together, our findings establish MPS1 overexpression as a driver of CIN and uncover how oncogenic transcriptional rewiring of mitotic signalling converts a transient microtubule-capture structure into a pathological source of merotely and chromosome mis-segregation in colon cancer.

## INTRODUCTION

Chromosomal instability (CIN), the persistent mis-segregation of chromosomes during mitosis, generates ongoing karyotypic diversity within tumours and is widely viewed as a key engine of tumour evolution (Vasan et al., 2019; Vasudevan et al., 2021; van Dijk et al., 2021; Chen et al., 2025; Mennie et al., 2026). By continually producing new aneuploid karyotypes, CIN fuels intratumor heterogeneity, accelerates clonal selection and can facilitate metastatic adaptation and therapeutic resistance (Carter et al., 2006; Lee et al., 2011; Andor et al., 2016; McGranahan et al., 2016; Bakhoum et al., 2018; Li et al., 2023; Al-Rawi et al., 2024). Yet, despite detailed knowledge of the cellular manifestations of CIN (lagging chromosomes, DNA bridges, micronuclei and aneuploid karyotypes), the proximal molecular events that initiate and sustain CIN in most human cancers remain poorly defined (Thompson et al., 2010; Bakhoum and Cantley, 2018; Hosea et al., 2024). Tackling this gap matters because distinct routes to CIN are likely to create different evolutionary trajectories, impose different stresses on tumour cells and offer targetable therapeutic vulnerabilities.

In human cells, CIN most commonly originates from errors in kinetochore-microtubule attachments (Cimini et al., 2001; Thompson and Compton, 2008; Bakhoum et al., 2009a; Bakhoum et al., 2009b; Thompson et al., 2010). Correct chromosome segregation requires sister kinetochores to be bioriented, establishing stable end-on attachments to microtubules from opposite poles, while avoiding erroneous attachment configurations such as syntelic attachment (both sisters attached to one pole) or merotelic attachment, in which a single kinetochore binds microtubules from both poles (Lampson et al., 2004; Cimini et al., 2006). Merotelic attachments are especially insidious because they can evade canonical spindle assembly checkpoint (SAC) surveillance (Cimini et al., 2001; Cimini et al., 2002; Salmon et al., 2005). A merotelic kinetochore can generate a lagging chromosome during anaphase as opposing pole-directed forces act on the same kinetochore (Cimini et al., 2002 Cimini et al., 2003; Gregan et al., 2011). Lagging chromosomes can subsequently form micronuclei, which are implicated in DNA damage, genome remodelling and catastrophic rearrangements, providing one path by which transient mitotic errors can be converted into durable genomic instability (Crasta et al., 2012; Hatch et al., 2013; Zhang et al., 2015; Mackenzie et al., 2017; Bakhoum et al., 2018; Umbreit et al., 2020; Krupina et al., 2021).

The fidelity of kinetochore-microtubule attachments is enforced by tightly coupled checkpoint, error-correction and attachment-maturation pathways. The SAC delays anaphase onset until kinetochores achieve appropriate attachment states, whereas error correction promotes turnover of improper attachments to enable re-attachment with correct geometry (Lara-Gonzalez et al., 2021; McAinsh and Kops, 2023). Central to these processes is the architecture of the outer kinetochore, dominated by the KMN network (KNL1:MIS12c:NDC80c), which provides the principal microtubule-binding interface, together with a dynamic set of accessory factors that tune attachment strength and signalling (Cheeseman et al., 2006; DeLuca et al., 2006; Wei et al., 2007; Ciferri et al., 2008). A particularly prominent transient feature of early mitotic kinetochores is the fibrous corona, a meshwork of proteins assembled on the outer kinetochore that expands the microtubule-interaction surface and supports microtubule capture, congression and SAC signalling. The corona is scaffolded by the RZZ complex (ROD:ZW10:Zwilch) and its interactor Spindly, and recruits additional components including Dynein:Dynactin modules, CENP-E, CENP-F and checkpoint proteins MAD1 and MAD2 (Yao et al., 1997; Starr et al., 1998; Scaërou et al., 2001; Gassmann et al., 2010; Musinipally et al., 2013; Mosalaganti et al., 2017; Pereira et al, 2018; Rodriguez-Rodriguez et al., 2018; Sacristan et al., 2018; d’Amico et al., 2022; Raisch et al., 2022; Cmentowski et al., 2023). During normal progression to metaphase, the kinetochore transitions from a capture state to a load-bearing end-on attachment state, and this transition is accompanied by corona compaction/stripping and reduced checkpoint signalling as biorientation is achieved (Griffis et al., 2007; Gassmann et al., 2010; Sacristan et al., 2018; Auckland et al., 2020; Barbosa et al., 2020; d’Amico et al., 2022; Eibes et al., 2023; Cmentowski et al., 2023; Ide et al., 2023). Multiple mitotic kinases coordinate these transitions. Aurora B and Aurora A kinases, enriched at centromeres and centrosomes respectively, phosphorylate outer-kinetochore substrates (including sites in the NDC80/HEC1 tail) to destabilize inappropriate attachments and promote error correction (Cheeseman et al., 2002; Tanaka et al., 2002; Cheeseman et al., 2006; DeLuca et al., 2006; Ciferri et al., 2008; Welburn et al., 2010; DeLuca et al., 2011; Chmátal et al., 2015; Ye et al., 2015; DeLuca et al., 2018). The kinase MPS1, also known as TTK in humans, is positioned at the nexus of these pathways. MPS1 orchestrates SAC signalling and has been implicated in tuning error-correction outputs through regulation of kinetochore substrates and crosstalk with Aurora pathways (Jelluma et al., 2008b; van der Waal et al., 2012;

Maciejowski et al., 2017; Sarangapani et al., 2021; Leça et al., 2025). MPS1 also plays a direct role in kinetochore remodelling by driving fibrous corona assembly. MPS1 phosphorylates the RZZ subunit ROD at Thr13 and Ser15, which promotes Spindly activation and triggers RZZ:Spindly oligomerization into the corona meshwork (Rodriguez-Rodriguez et al., 2018; Raisch et al., 2022). Thus, in principle, dysregulation of MPS1 could affect chromosome segregation through multiple routes: checkpoint signalling, attachment turnover, error-correction circuitry and kinetochore architecture.

Colon cancer presents a striking and clinically important context in which CIN is common but mechanistically unresolved. At a broad level, colon cancers are commonly stratified into microsatellite-instable (MSI) tumours, characterised by mismatch-repair deficiency and a hypermutator phenotype, and microsatellite-stable (MSS) tumours, in which CIN is the predominant form of genome instability. The canonical multistep genetic model of colon tumorigenesis, best exemplified by chromosomally unstable MSS tumours, involves Adenomatous Polyposis Coli (APC) inactivation, followed by KRAS activation and later impairment of P53 and TGF-β function. This model captures key signalling events that underpin malignant progression, but it does not provide a satisfying mechanistic explanation for chromosome mis-segregation that drives CIN (Vogelstein et al., 1988; Baker et al., 1989; Powell et al., 1992; Takagi et al., 1996; Thiagalingam et al.,1996; Pino and Chung, 2010). CIN is detectable early in benign adenomas and can occur in lesions with heterogeneous or incomplete driver trajectories (Shih et al., 2001; Hermsen et al., 2002; Giaretti et al., 2004a; Giaretti et al., 2004b; Pino and Chung, 2010). Cytogenetic and sequencing studies have described adenomas with intact *APC* that nonetheless exhibit chromosomal gains and losses (Giaretti et al., 2004a; Giaretti et al., 2004b). Moreover, CIN has been shown to correlate more strongly with adenoma size, dysplasia grade and proliferative activity rather than *APC* mutation status alone (Hermsen et al., 2002; Giaretti et al., 2004a; Giaretti et al., 2004b). These observations argue that the canonical *APC–KRAS–SMAD4–TP53* trajectory, while central to colon tumorigenesis, is insufficient by itself to explain the origin of recurrent segregation errors in many cases. Instead, oncogenic signalling may create a CIN permissive state in which altered expression or activity of mitotic regulators act as the proximal drivers of CIN. Consistent with this view, large-scale tumour datasets repeatedly point to a transcriptional rewiring of mitotic programmes in chromosomally unstable MSS tumours of the colon (Lengauer et al., 1997; Carter et al., 2006; Pino and Chung, 2010).

Among mitotic regulators, MPS1 stands out as a compelling candidate CIN driver. Across many tumour types, MPS1 is frequently upregulated as part of a broader proliferation/mitotic transcriptional programme and has repeatedly been associated with aggressive disease, aneuploidy signatures and poor prognosis (Yuan et al., 2006; Ling et al., 2014; Slee et al., 2014; Miao et al., 2016; Choi et al., 2017; Xie et al., 2017; King et al., 2018). MPS1 is also part of the CIN70 chromosomal instability expression signature, a gene set whose overexpression correlates with aneuploidy/CIN across tumour cohorts (Carter et al., 2006; Slee et al., 2014). In parallel, pharmacological and genetic studies have suggested that aneuploid tumour cells become dependent on MPS1 activity for proliferation and survival under mitotic stress, raising the possibility that MPS1 contributes to the maintenance of CIN permissive states (Daniel et al., 2011; Maire et al., 2013; Slee et al., 2014; Maia et al., 2015; Zhang et al., 2016; Anderhub et al., 2019; Chandler et al., 2020). A long-standing question, however, is whether elevated MPS1 is merely an adaptive response to mitotic stress in already aneuploid cells, bolstering checkpoint signalling to tolerate instability, or whether it can actively drive the segregation errors that sustain CIN and promote tumour evolution. Here we address this question by integrating tumour-scale genomics with mechanistic cell biology in disease-relevant models. Building on the observation that CIN^+^ colon carcinomas exhibit elevated MPS1 levels, we combine analyses of primary tumour cohorts with low-passage patient-derived colon cancer cell (PCCC) lines and engineered near-diploid epithelial cells to test whether MPS1 overexpression is sufficient to drive CIN, whether it causally contributes to chromosome mis-segregation, and to define the underlying mechanism. By connecting tumour-associated MPS1 overexpression to a concrete, targetable structural defect at the kinetochore, this work provides a conceptual bridge between oncogenic transcriptional programmes and the mitotic errors that generate karyotypic diversity in colon cancer.

## RESULTS

### MPS1 overexpression is prevalent in chromosomally unstable colon carcinoma and is associated with a worse clinical outcome

While the canonical *APC–KRAS–SMAD4–TP53* pathway accounts for colon tumorigenesis, it does not sufficiently explain the recurrent chromosome mis-segregation events that define CIN (Vogelstein et al., 1988; Baker et al., 1989; Powell et al., 1992; Takagi et al., 1996; Thiagalingam et al.,1996; Pino and Chung, 2010 ; Dow et al., 2015). Specifically, CIN is observed in *APC*-wild-type lesions and early adenomas, suggesting it may arise independently of these classic drivers (Hermsen et al., 2002; Giaretti et al., 2004a; Giaretti et al., 2004b). We therefore asked whether CIN in colon cancer is associated with alterations in the mitotic machinery itself, focusing first on whether core mitotic regulators are recurrently mutated or transcriptionally deregulated in chromosomally unstable tumours. To address this, we performed a mitosis-focused analysis of the AC-ICAM colon cancer cohort, which provides comprehensive genomic profiles of fresh-frozen primary tumours and matched clinical endpoints (Roelands et al., 2023). We examined somatic mutations in 44 mitotic genes across 281 tumours stratified by CIN, based on whole-exome-wide copy-number alteration (CNA) burden, and categorized by microsatellite instability status (MSI-H vs MSS) (**Fig. 1A**). Most mitotic genes were mutated in MSI-H tumours, albeit at varying frequencies. In contrast, these genes were rarely mutated in MSS tumours, which instead showed the expected enrichment of canonical CIN-associated mutations in *APC*, *KRAS* and *TP53* (**Fig. 1B**). This pattern was independently reproduced in 531 tumours of the TCGA colon cancer cohort (**Appendix Fig. S1A**), reinforcing that mitotic genes are rarely mutated in MSS colon cancer. To minimize confounding by the mismatch-repair–driven hypermutator phenotype and the distinct tumour-immune context of MSI tumours, we limited subsequent analyses to MSS cases and compared CIN^+^ versus CIN^−^ tumours to better isolate CIN-associated mechanisms. MSS tumours share the same broad instability context (no mismatch-repair hypermutation), so differences between CIN^+^ and CIN^−^ tumours are more likely to reflect CIN-related mechanisms. Expression analysis of the 44 mitotic genes in MSS tumours (**Fig. 1C**) revealed that *AURKA* (Aurora A), *SKA3*, and *MPS1/TTK* were the three most upregulated genes in CIN^+^ samples (**Fig. 1D**). Under normal physiological conditions, MPS1 phosphorylates both Aurora A and SKA3 to destabilize kinetochore-microtubule attachments and enable error correction (Maciejowski et al., 2017; Leça et al., 2025). Given this upstream regulatory role, we investigated whether elevated MPS1 expression could contribute to CIN in colon cancer. Consistent with CIN promoting tumour aggressiveness, CIN^+^ MSS tumours showed poorer clinical outcomes than CIN^−^ MSS tumours in the AC-ICAM cohort (**Fig. 1E,F**). In stage III and stage IV colon cancer, high MPS1 expression was associated with a trend toward reduced overall survival (**Fig. 1G,H**), and analyses of independent cohorts supported the association between high MPS1 expression and worse prognosis (**Appendix Fig. S1B–E**). Together, these data identify MPS1 overexpression as a recurrent feature of poor-prognosis, chromosomally unstable colon carcinomas.

**Figure 1.**
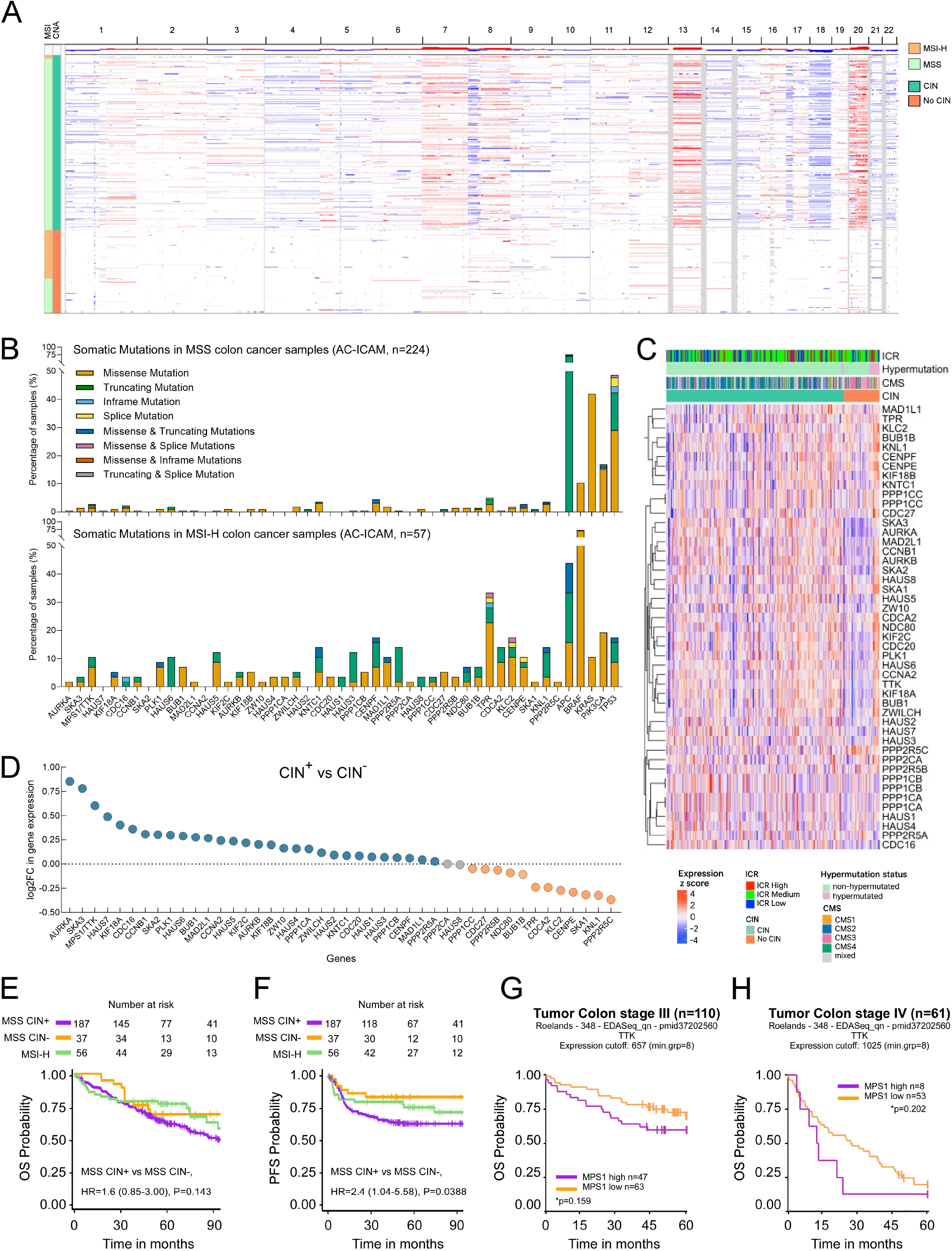
MPS1 overexpression is prevalent in chromosomally unstable colon carcinoma and is associated with a worse clinical outcome. (A) Heat map of copy number alterations (CNAs) across the 22 autosomes in the AC-ICAM cohort (N = 281, Roelands et al., 2023). Each row represents an individual sample and columns correspond to genomic positions ordered by chromosome (1– 22). Copy number gains are indicated in red and losses in blue, with colour intensity reflecting the magnitude of deviation from the baseline. Samples are grouped according to chromosomal instability (CIN) classification (CIN vs. No CIN) and categorized for microsatellite instability status (MSI-H vs MSS), as indicated by the annotation bars on the left. **(B)** Frequency of somatic mutations in 44 mitotic genes across the 281 tumours of the AC-ICAM cohort stratified by microsatellite instability status (MSI-H vs MSS). **(C)** Heat map of normalized log2-transformed expression of 44 mitotic genes across MSS colon tumours stratified by CIN status (CIN vs. No CIN) based on whole exome-wide CNA burden (row-wise z-scores; each gene z-scored across samples). Columns represent MSS tumour samples (AC-ICAM, n = 224) annotated with immune classification of response (ICR), consensus molecular subtype (CMS) and MSI status. **(D)** Log2 fold change in expression of 44 mitotic genes in CIN^+^ vs. CIN^−^ MSS tumours (AC-ICAM, n = 224). Blue, grey, and orange dots indicate genes that are upregulated, unaltered, or downregulated in CIN^+^ relative to CIN^−^ MSS tumours, respectively. **(E, F)** Kaplan–Meier plots of MSS CIN^+^, MSS CIN^-^ and MSI-H groups (AC-ICAM) for overall survival (OS; **E**) and progression-free survival (PFS; **F**). Vertical lines denote censored observations. Statistical comparisons were performed only within MSS tumours, comparing MSS CIN^+^ versus MSS CIN^−^. Hazard ratios (HRs) and 95% confidence intervals are calculated by Cox proportional hazard regression. The corresponding Cox model *P* values (Wald test) are displayed. **(G, H)** Kaplan–Meier plots of overall survival (OS) for patients with stage III **(G)** and stage IV **(H)** colon cancer from the AC-ICAM cohort, stratified by MPS1 expression. Patients were dichotomised into MPS1-high and MPS1-low groups using the expression cut-offs indicated. Vertical lines denote censored observations. Depicted *\*P* values compare overall differences in survival between expression-defined groups and were calculated using a two-sided log-rank test.

### Patient-derived colon cancer cells (PCCCs) recapitulate core hallmarks of CIN and provide disease-relevant models for mechanistic analysis

To determine whether MPS1 overexpression contributes to CIN and to dissect the underlying mechanism in a disease-relevant experimental system, we used a panel of low passage patient-derived colon cancer cell (PCCC) lines that were established from primary tumours and metastases (**Fig. 2A**). Unless otherwise indicated, experiments were performed using PCCC cultures between passages 20 and 30 to minimise potential effects of culture-associated adaptation. Previous genetic, genomic and transcriptomic profiling confirmed their molecular heterogeneity, spanning distinct molecular backgrounds, including MSS and MSI genotypes and recurrent alterations in canonical colon cancer driver genes (**Fig. 2B**; Boot et al., 2016). CNA and loss-of-heterozygosity analysis showed that the MSI line JVE059 is near-diploid, whereas the MSS lines JVE187, JVE207 and KP363T harbour varying degrees of chromosomal gains and losses (**Appendix Fig. S2A**; Boot et al., 2016). However, such karyotypic imbalances represent a genomic state and do not, by themselves, demonstrate ongoing CIN driven by elevated chromosome mis-segregation rates. We therefore introduced EGFP-αTubulin and H2B-mRFP into PCCCs to visualize spindles and chromatin during live mitosis and directly quantify chromosome segregation errors by live-cell imaging. Compared with near-diploid RPE-1 and JVE059 cells, JVE187, JVE207 and KP363T cells exhibited frequent anaphase abnormalities, principally lagging chromosomes and DNA bridges (**Fig. 2C,D**; **Movies EV1–EV8**). These ongoing segregation errors, which are established drivers of CIN (Cimini et al., 2002; Thompson and Compton, 2008), together with the genomic imbalances observed (**Appendix Fig. S2A**), led us to classify the PCCC lines JVE187, JVE207 and KP363T as chromosomally unstable (CIN^+^). High-resolution immunofluorescence of anaphase cells revealed that lagging chromosomes in CIN^+^ PCCCs harboured kinetochores attached to microtubules emanating from both spindle poles (**Fig. 2E,F**). This pattern is consistent with merotelic attachments that evade correction before anaphase onset, rather than premature anaphase entry in the presence of unattached or weakly attached kinetochores. In line with this, live-cell imaging showed that asynchronously cultured CIN^+^ PCCCs congressed chromosomes efficiently and entered anaphase only after alignment at the metaphase plate, with prometaphase and metaphase durations comparable to those of RPE-1 cells and the MSI line JVE059 (**Appendix Fig. S2B,C; Movies EV9-EV13**). Moreover, nocodazole treatment triggered a robust mitotic arrest in CIN^+^ PCCCs comparable to RPE-1 and JVE059, indicating an intact SAC (**Appendix Fig. S2D**) and further arguing against checkpoint failure as the source of lagging chromosomes. To directly assess kinetochore-microtubule attachment stability, we analysed microtubule dynamics using fluorescence dissipation after photoactivation. We quantified the turnover of photoactivatable (PA)-GFP-αTubulin in prometaphase and metaphase spindles, staging cells by chromosome alignment using DIC optics. In RPE-1 cells, the half-life of kinetochore-bound microtubules (kMTs) increased stepwise from prometaphase to metaphase (**Appendix Fig. S2E**), consistent with prior measurements and the maturation of stable end-on attachments at metaphase (Kabeche and Compton, 2013). Notably, metaphase kMT half-life was further elevated across CIN PCCCs (**Appendix Fig. S2E**), indicating hyper-stable kinetochore-microtubule attachments, a hallmark previously linked to CIN (Bakhoum et al., 2009a). Despite these hyper-stable attachments, CIN PCCCs retained the capacity for error correction. Following monastrol-induced monopolar spindle formation (produces syntelic attachments) and subsequent washout, CIN PCCCs realigned chromosomes to the metaphase plate with efficiencies comparable to RPE-1 and MSI cells, indicating effective correction of the initially erroneous attachments (**Appendix Fig. S2F**).

**Figure 2.**
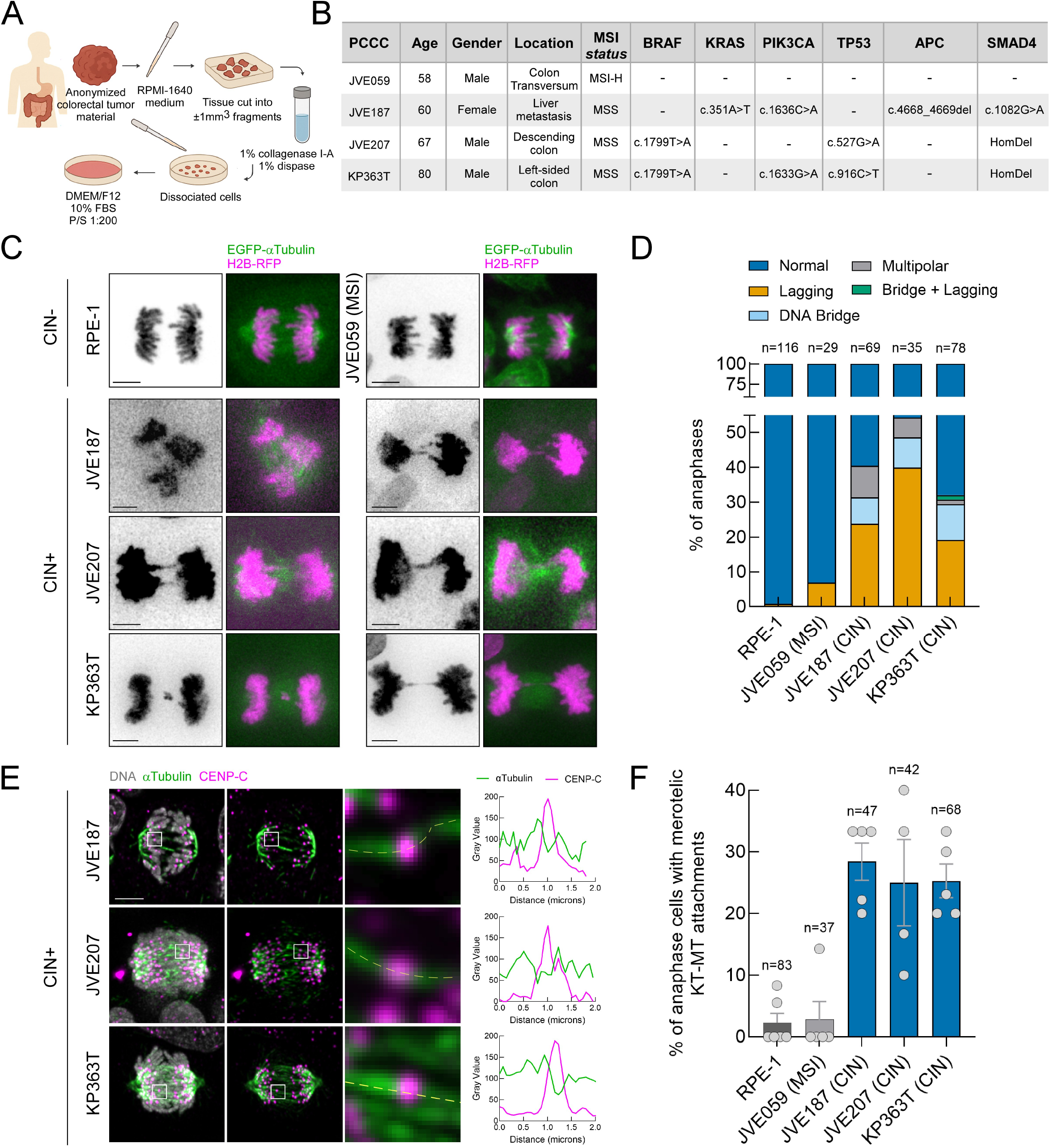
Patient-derived colon cancer cells (PCCCs) recapitulate core hallmarks of CIN and provide disease-relevant models for mechanistic analysis. (A) Schematic representation of the experimental pipeline for the establishment of the low passage patient-derived colon cancer cell (PCCC) lines from anonymized tumour material as in Boot et al. (2016). **(B)** PCCC lines characteristics and key mutation signatures, as reported by Boot et al. (2016). **(C)** Representative still frames from live-cell imaging of chromosome segregation in RPE-1 cells and PCCCs JVE059 (MSI), JVE187 (CIN), JVE207 (CIN) and KP363T (CIN). Cells stably expressed H2B–RFP (chromosomes, magenta) and EGFP– αTubulin (spindle microtubules, green). **(D)** Quantification of chromosome segregation fidelity for the cell lines shown in **(C)**. Graph bars represent the frequency of indicated anaphase output. n denotes the number of anaphase events analysed (filmed) for each cell line. **(E)** Representative immunofluorescence images of CIN^+^ PCCCs in anaphase displaying merotelic kinetochore–microtubule attachments. Insets show magnified views of the boxed regions. Line-scan intensity profiles of CENP-C and αTubulin fluorescence are plotted as a function of distance (µm) along the dashed yellow line in the inset. **(F)** Quantification of the percentage of anaphase cells exhibiting merotelic kinetochore–microtubule attachments as in **(E)**; each dot represents an independent experiment. n denotes the total number of anaphase cells analysed for each cell line. Data information: data in **(D)** are plotted as fractions of the total and data in **(F)** are presented as mean with standard error of the mean (SEM). Scale bars: 5μm.

In conclusion, the PCCC lines JVE187, JVE207 and KP363T recapitulate core hallmarks of CIN, combining copy-number imbalance with ongoing chromosome mis-segregation dominated by lagging chromosomes arising from persistent merotelic kinetochore-microtubule attachments. Importantly, these defects occur despite efficient chromosome congression, normal mitotic timing and an intact SAC. This molecularly characterised and genetically diverse patient-derived panel therefore provides a disease-relevant and experimentally tractable system to investigate if and how MPS1 overexpression contributes to CIN.

### Elevated MPS1 activity drives chromosome segregation errors in CIN^+^ PCCCs

To test whether elevated MPS1 levels drive CIN in colon carcinoma, we first asked whether MPS1 is likewise overexpressed in CIN^+^ PCCCs, as observed in CIN^+^ colon tumours. Transcriptome-wide expression estimates for all PCCC lines (17090 genes) were derived from cDNA hybridization–intensity data generated on Infinium HumanExome-12 v1 BeadChips (Boot et al., 2016). For each mitotic gene analysed above, we plotted the expression across individual CIN+ PCCC lines after normalisation to the near-diploid MSI JVE059 (**Fig. 3A**). *MPS1* was consistently upregulated in all CIN+ PCCCs relative to JVE059. Increased *MPS1* mRNA and protein levels were independently confirmed by qPCR and immunoblotting, respectively (**Fig. 3B,C**). Thus, *MPS1* is overexpressed in chromosomally unstable PCCCs, mirroring CIN^+^ colon tumours.

**Figure 3.**
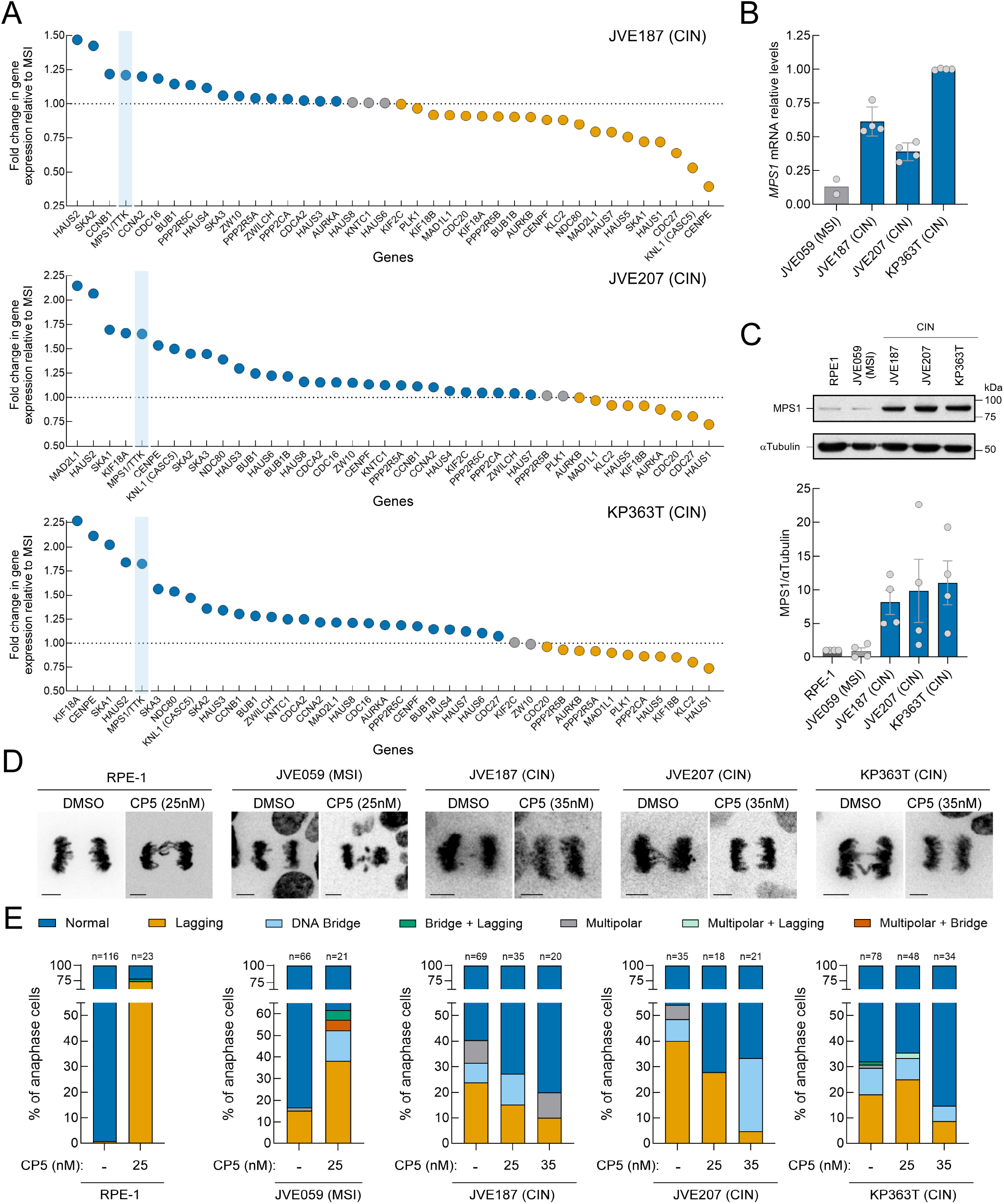
Elevated MPS1 activity drives chromosome segregation errors in CIN^+^ PCCCs. (A) Fold change in expression of mitotic genes in CIN^+^ PCCCs relative to the MSI cell line. Data are expressed as a ratio relative to the MSI cell line, arranged in descending order from left to right. Blue, grey, and orange dots indicate genes that are upregulated, unaltered, or downregulated in CIN^+^ relative to MSI (CIN^−^) PCCCs, respectively. **(B)** Relative *MPS1* mRNA levels measured by RT-qPCR. *MPS1* mRNA abundance in each experimental condition was determined relative to the mRNA abundance of the housekeeping gene *CPSF6*. All *MPS1*/*CPSF6* values were normalized to the mean value determined for KP363T cells, which was set to 1. Each dot represents an independent replicate. **(C)** Relative MPS1 protein levels measured by Western blot. MPS1 protein abundance in each experimental condition was determined relative to the loading control αTubulin. All MPS1/ αTubulin values were normalized to the mean value determined for RPE-1 cells, which was set to 1. Each dot represents an independent replicate. **(D)** Representative still frames from live-cell imaging of chromosome segregation in RPE-1 cells and PCCCs, under control conditions (DMSO) and following treatment with the MPS1 inhibitor compound 5 (CP5). Chromosomes (H2B-RFP) are represented in inverted greyscale. **(E)** Quantification of chromosome segregation fidelity for the cell lines shown in **(D)**. Graph bars represent the frequency of indicated anaphase output. n denotes the number of anaphase events analysed (filmed) for each cell line. Data information: data in **(B)** and **(C)** are presented as mean with standard deviations (SD) and data in **(E)** are plotted as fractions of the total. Scale bars: 5μm.

We next assessed whether MPS1 activity was also increased at kinetochores of CIN^+^ PCCCs. We measured MPS1 activation by immunofluorescence using an antibody against the activating Thr676 phosphorylation in the MPS1 T-loop (MPS1^pT676^). In chromosomally stable cells, kinetochore-associated MPS1 activity normally declines as chromosomes biorient and progress towards metaphase (Moura et al., 2017; Hayward et al., 2019), and accordingly RPE-1 and JVE059 cells exhibited an ∼50% reduction in kinetochore MPS1^pT676^ levels from prometaphase to metaphase (**Appendix Fig. S3A,B**). By contrast, this decline was significantly attenuated in all CIN^+^ PCCCs, indicating sustained MPS1 activity at metaphase (**Appendix Fig. S3A,B**).

Because CIN^+^ PCCCs retain elevated levels of active MPS1 at metaphase kinetochores, we then asked whether dampening MPS1 activity could suppress segregation errors in these cells without abrogating the SAC. Treatment with suboptimal doses (25 nM and 35 nM) of the MPS1 specific inhibitor CP5 (Koch et al., 2016) partially reduced kinetochore MPS1^pT676^ at metaphase (**Appendix Fig. S3A,B**) and significantly decreased the frequency of lagging chromosomes and other anaphase defects (**Fig. 3D,E**). In contrast, the same low doses of CP5 were sufficient to compromise the SAC in RPE-1 and JVE059 cells, leading to frequent chromosome mis-segregation (**Fig. 3D,E**). Collectively, these data indicate that CIN^+^ PCCCs operate with an abnormally elevated set point of kinetochore MPS1 activity at metaphase and that this heightened signalling contributes to chromosome segregation errors. Importantly, partial MPS1 inhibition reduces mis-segregation in CIN^+^ PCCCs without overt checkpoint collapse, consistent with a CIN-permissive mitotic state sustained by elevated MPS1 activity. Therefore, we conclude that elevated MPS1 is functionally linked to the CIN phenotype in PCCCs.

### MPS1 overexpression is sufficient to induce CIN in otherwise stable near diploid cells

To test whether increased MPS1 levels are sufficient to drive CIN, we generated RPE-1 and JVE059 cells ectopically expressing EGFP-MPS1 under either doxycycline-inducible promoter (P_tet_) or a constitutive promoter (P_CMV_) (**Fig. 4A** and **Appendix Fig. S4A,B**). Live-cell imaging showed that MPS1 overexpression prolonged metaphase in both RPE-1 and colon JVE059 cells (**Fig. 4B** and **Appendix Fig. S4A**). Accordingly, RPE-1 cells overexpressing MPS1 retained elevated MAD1 at metaphase kinetochores, suggesting that the metaphase delay was due to enduring SAC signalling (**Fig. EV1A-C**). Importantly, both acute (doxycycline-induced) and constitutive MPS1 overexpression significantly increased the frequency of anaphases with lagging chromosomes in RPE-1 and JVE059 cells (**Fig. 4C,D**; **Appendix Fig. S4B,C** and **Movies EV14-EV21**). Moreover, long-term constitutive overexpression of MPS1 promoted a progressive accumulation of segregation errors across passages, with lagging chromosomes remaining the most frequent abnormality (**Fig. 4E** and **Appendix Fig. S4D**). The vast majority of lagging events arose from chromosomes that were aligned at the metaphase plate, with only a small fraction originating from misaligned chromosomes (**Fig. EV1D,E**). We then monitored the fate of lagging chromosomes in RPE-1 cells that constitutively overexpressed MPS1. Live-cell imaging showed that lagging chromosomes often culminated in micronuclei formation and that these events were recurrent across passages (**Fig. 4F,G and Movie EV22**). Chromosome mis-segregation can lead to both numerical and structural aneuploidy that can further promote chromosome instability in the following cell cycles (Garribba et al, 2023). Accordingly, multicolour FISH (mFISH) of RPE-1 and JVE059 cells across passages identified recurrent whole-chromosome gains and losses andstructural rearrangements in independent EGFP-MPS1 clones compared with the parental near-diploid karyotype (**Fig. 4H**; **Fig. EV2A,B** and **Appendix Fig. S4E**). This indicates that sustained overexpression of MPS1 in RPE-1 and MSI colon cells resulted in progressive karyotypic divergence. Based on these results, we conclude that MPS1 overexpression is sufficient to induce ongoing chromosome segregation errors and to fuel karyotypic evolution in otherwise chromosomally stable epithelial cells.

**Figure 4.**
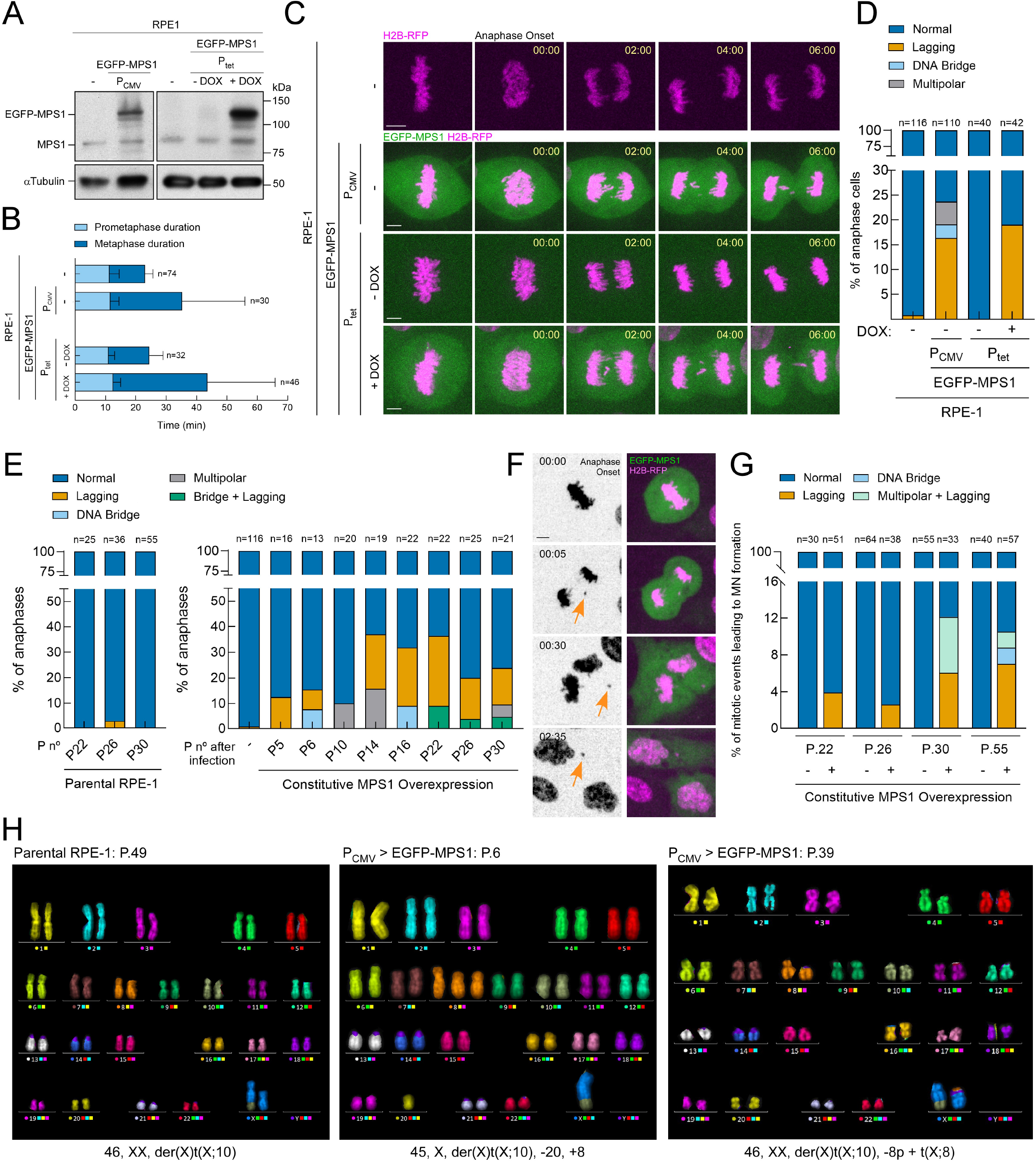
MPS1 overexpression is sufficient to induce CIN in otherwise stable near diploid RPE-1 cells. (A) Representative immunoblot of MPS1 in parental RPE-1 cells, RPE-1 cells with constitutive MPS1 overexpression (P_CMV_), and RPE-1 cells carrying a doxycycline-inducible MPS1 expression system (P_tet_). α-tubulin was used as a loading control. **(B)** Mitotic timing of cells indicated in **(A)**. Light-coloured bars represent the prometaphase duration (length of time measured between NEB and the first frame with all chromosomes aligned at the metaphase plate) and dark-coloured bars represent the metaphase duration (length of time measured between the first frame after chromosome alignment and anaphase onset). n indicates the number of cells filmed for each condition. **(C)** Selected still frames from representative live-cell imaging of chromosome segregation in RPE-1 cells, RPE-1 cells constitutively overexpressing MPS1 (P_CMV_) and RPE-1 cells carrying a doxycycline-inducible MPS1 expression system (P_tet_). All cells stably express H2B-RFP to visualize the chromosomes (magenta). MPS1 overexpression is shown in green for both P_CMV_ and P_tet_ (+DOX) conditions. Time (min) is shown relative to anaphase onset. Scale bar: 5 µm. **(D)** Quantification of chromosome segregation fidelity for the cell lines shown in **(C)**. Graph bars represent the frequency of indicated anaphase output. n denotes the number of anaphase events analysed (filmed) for each cell line. **(E)** Quantification of chromosome segregation fidelity for parental RPE-1 cells (left) and RPE-1 cells constitutively overexpressing MPS1 (right) over several passages. Graph bars represent the frequency of indicated anaphase output. n denotes the number of anaphase events analysed (filmed) for each cell line and for each passage. **(F)** Selected still frames from representative live-cell imaging of micronuclei formation in RPE-1 cells overexpressing MPS1 (green) and stably expressing H2B-RFP to visualize the chromosomes (magenta). Orange arrowheads track the lagging chromosome until it eventually forms a micronuclei. Time (min) is shown relative to anaphase onset. Scale bar: 10μm. **(G)** Quantifications of the mitotic events leading to micronuclei (MN) formation in control parental RPE-1 cells (-) and RPE-1 cells overexpressing MPS1 (+) over several passages. Graph bars represent the frequency and mitotic origin of micronuclei. n denotes the number of anaphase events analysed (filmed) for each cell line and for each passage. **(H)** Representative multicolour fluorescence in situ hybridization (mFISH) karyograms of a parental RPE-1 cell at passage 49 (P49; left) and of RPE-1 cells constitutively overexpressing CMV-driven EGFP-MPS1 analysed at passage 6 (P6; middle) and at passage 39 (P39; right). Corresponding karyotypes are indicated below each karyogram. Data information: data in **(B)** are presented as mean with standard deviations and data in **(D)**, **(E)** and **(G)** are plotted as fractions of the total.

### MPS1 overexpression promotes merotely independently of global changes in kinetochore-microtubule stability or Aurora B signalling

To define the basis of lagging chromosomes in RPE-1 cells overexpressing MPS1, we first assessed their kinetochore-microtubule attachment configurations after selective depolymerization of non-kinetochore microtubules by brief calcium treatment. Immunofluorescence analysis revealed a marked increase in anaphase cells containing chromosomes with merotelic attachments, a configuration rarely observed in control RPE-1 cells expressing endogenous MPS1 (**Fig. 5A–C**).

**Figure 5.**
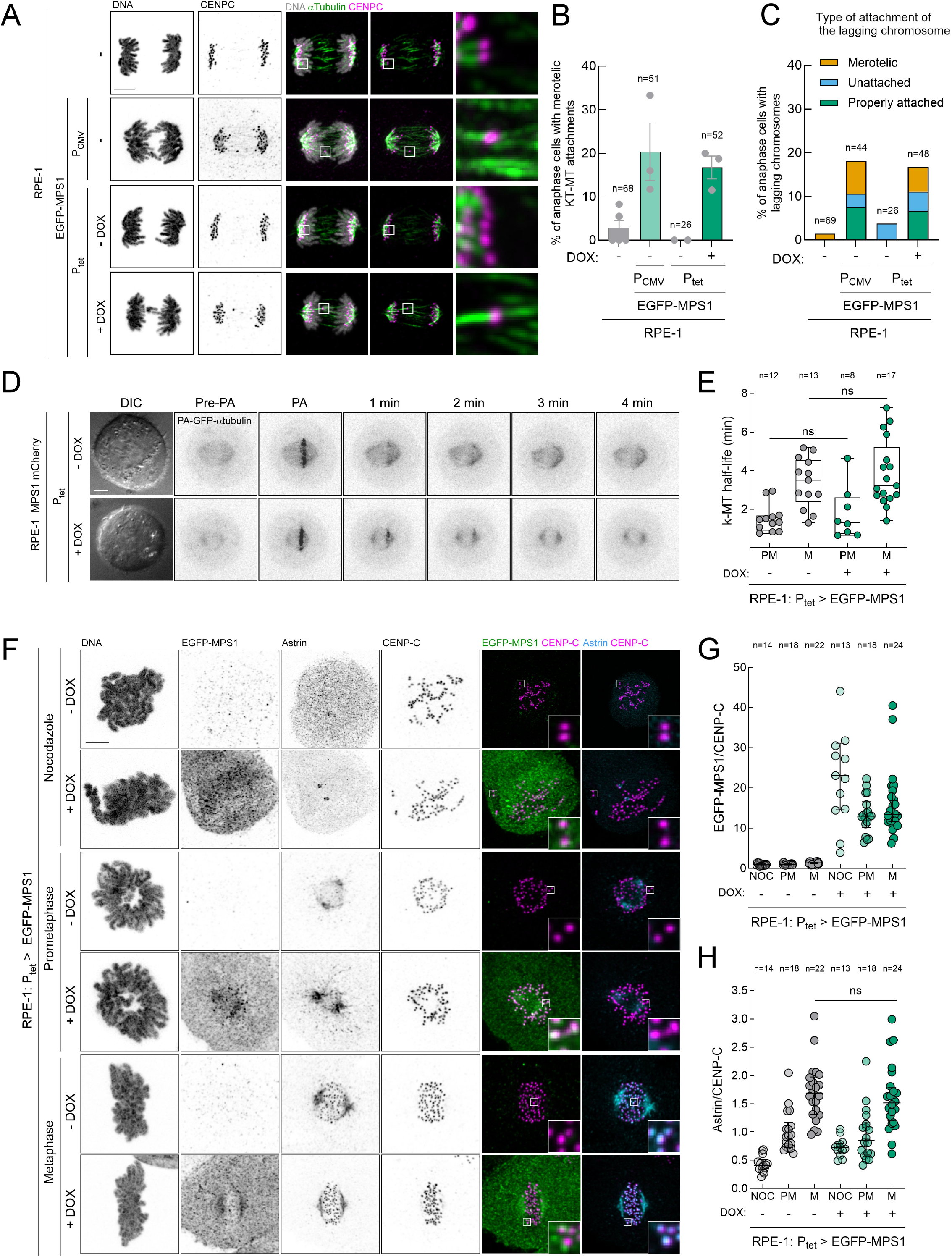
MPS1 overexpression promotes merotely independently of global changes in kinetochore-microtubule stability. (A) Representative immunofluorescence images of calcium-treated cells to visualise stable kinetochore-microtubule attachments in anaphase in parental RPE-1 cells, RPE-1 cells constitutively overexpressing MPS1 (P_CMV_) and RPE-1 cells carrying a doxycycline-inducible MPS1 expression system (P_tet_). Insets display magnifications of the outlined regions. **(B)** Quantification of the percentage of anaphase cells exhibiting merotelic kinetochore–microtubule attachments in the conditions depicted in **(A)**. Each dot represents an independent experiment. n indicates the number of cells analysed for each condition. **(C)** Frequency of anaphase cells with lagging chromosomes exhibiting either a merotelic kinetochore-microtubule attachment (orange bars), an unattached kinetochore (blue bars) or a properly attached kinetochore (green bars). n denotes the number of anaphase cells analysed for each cell line. **(D)** Representative stills of tubulin photoactivation experiments in live metaphase control RPE-1 cells (-DOX) and RPE-1 cells overexpressing MPS1 (+DOX). Panels represent DIC and the PA–GFP–αTubulin signal before photoactivation (Pre-PA), immediately after photoactivation (PA), and after photoactivation. The PA–GFP– αTubulin signal was inverted for better visualization. **(E)** Quantification of kinetochore-microtubule half-life for the cell lines in **(D)** in prometaphase (PM) and metaphase (M). Each data point represents an individual cell. The boxes represent median and interquartile range; the bars represent minimum and maximum values. n indicates the number of cells analysed for each condition. **(F-H)** Representative immunofluorescence images **(F)** and corresponding quantifications of EGFP-MPS1 levels **(G)** and Astrin levels **(H)** at kinetochores of RPE-1 cells in control (-DOX) and MPS1 overexpression (+DOX) conditions, in nocodazole (NOC), prometaphase (PM) and metaphase (M). Insets depict magnifications of selected kinetochores. EGFP-MPS1 and Astrin fluorescence intensities were determined relative to the CENP-C signal. All values in (G,H) were normalized to the mean value determined for prometaphase control RPE-1 cells, which was set to 1. Each data point represents the mean of an individual cell. Data information: data in **(B)** are presented mean with standard error (SEM) of the mean, data in **(C)** are presented as mean, and data in **(E)**, **(G)** and **(H)** are presented as median with interquartile range; ns, non-significant; Kruskal-Wallis, Dunn’s multiple comparison test in **(E)**, and Mann-Whitney test in **(H)**. Scale bars: 5μm.

We next tested whether MPS1 overexpression altered kinetochore–microtubule attachment stability. Fluorescence dissipation after photoactivation of PA–GFP– αTubulin in prometaphase and metaphase spindles showed that MPS1 induction (+DOX) did not significantly change kMT half-life compared with non-induced controls (−DOX) (**Fig. 5D,E**), indicating that elevated MPS1 does not measurably perturb kMT turnover and that cells still establish stable attachments in metaphase. In line with this, tubulin intensity within kinetochore-fibers (microtubule bundles attached to kinetochores) increased from prometaphase to metaphase to a similar extent in control and MPS1-overexpressing cells (**Appendix Fig. S5A**,**B**). Moreover, the predominant kinetochore-microtubule attachment configuration in metaphase cells remained end-on under both conditions (**Appendix Fig. S5C**), further supporting that MPS1 overexpression does not destabilize mature attachments. As an independent readout, Astrin, which selectively decorates stably end-on attached kinetochores (Mack et al., 2001; Manning et al., 2010; Conti et al., 2019), accumulated at metaphase kinetochores to similar levels in control and MPS1-overexpressing RPE-1 cells, consistent with preserved attachment maturation despite elevated MPS1 (**Fig. 5F-H**). Notably, despite normal end-on attachment formation, metaphase kinetochores in MPS1-overexpressing cells exhibited increased phosphorylation of KNL1 at Thr943 and Thr1155 (KNL1^pT943/pT1155^), established MPS1 target sites (Yamagishi et al., 2012; Vleugel et al., 2015), indicating that a kinetochore-localised pool of MPS1 remains catalytically active at metaphase under these conditions (**Appendix Fig. S6A-C**).

Given that Aurora B activity is required for the resolution of merotelic attachments (Andrews et al., 2004; Knowlton et al., 2006; Cimini et al., 2006; Liang et al., 2020), we next asked whether aberrant Aurora B signalling might account for the excess of merotelic attachments observed upon MPS1 overexpression. However, immunofluorescence analysis revealed no detectable change in Aurora B activation, as assessed by T-loop phosphorylation at Thr232 (AurB^pT232^) in MPS1-overexpressing cells compared with controls (**Appendix Fig. S6D-F**). Consistently, phosphorylation of the Aurora B outer-kinetochore substrate HEC1 at Ser44 (HEC1^pS44^), a modification required for efficient kinetochore–microtubule error correction (DeLuca et al., 2006; DeLuca et al., 2011), was also unchanged under MPS1-overexpressing conditions (**Appendix Fig. S6G-I**). These data, together with the ability to congress chromosomes efficiently and in a timely manner (**Fig. 4B**), suggest that Aurora B-mediated error correction remains largely intact in RPE-1 cells overexpressing MPS1. Collectively, these findings indicate that MPS1 overexpression promotes merotelic attachments and lagging chromosomes in RPE-1 cells through a distinct mechanism independent of global changes in kinetochore-microtubule stability or Aurora B signalling.

### MPS1 overexpression sustains fibrous corona assembly signalling at metaphase kinetochores

We next asked whether elevated MPS1 perturbs kinetochore architecture or composition in a manner that favours aberrant microtubule capture. We focused on the fibrous corona because its assembly at the outer kinetochore in early mitosis requires MPS1 activity, and Monte Carlo–based modelling predicts that incomplete corona disassembly at metaphase predisposes kinetochores to merotelic attachments (Krivov et al., 2021). We therefore tested whether MPS1 overexpression perturbs fibrous corona dynamics, which is regulated by MPS1-mediated phosphorylation of the RZZ subunit ROD. Phosphorylation of the ROD N-terminus at Thr13 and Ser15 promotes Spindly activation and its oligomerization with the RZZ complex to build the core corona RZZ:Spindly meshwork (Rodriguez-Rodriguez et al., 2018; Raisch et al., 2022). We therefore quantified kinetochore ROD phosphorylation and Spindly accumulation as cells progressed from prometaphase to metaphase. In control RPE-1 cells expressing endogenous MPS1, kinetochore levels of phosphorylated ROD (ROD^pT13/pS15^) declined markedly upon chromosome alignment at the metaphase plate, consistent with timely fibrous corona disassembly as sister kinetochores biorient during normal mitosis (**Fig. 6A,B**). In sharp contrast, MPS1-overexpressing cells retained high levels of kinetochore ROD^pT13/pS15^ at metaphase chromosomes (**Fig. 6A,B and Appendix Fig. S7A,B**). This persistence was abolished by acute treatment with the MPS1 inhibitor CP5, confirming that sustained phosphorylation of ROD at Thr13 and Ser15 reflects ongoing MPS1 kinase activity (**Fig. 6A,B**). High levels of ROD^pT13/pS15^ persisting at metaphase kinetochores upon MPS1 overexpression were accompanied by excessive accumulation of Spindly, quantified by both immunofluorescence intensity and three-dimensional signal volume (**Fig. 6C–E and Appendix Fig. S7C,D**). We next asked whether additional fibrous corona components were likewise aberrantly retained at metaphase kinetochores in MPS1-overexpressing cells. In line with Spindly, both ZW10 and CENP-F showed increased kinetochore signal intensity and volume upon MPS1 overexpression (**Fig. EV3A–F** and **Appendix Fig. S7E–H)**. By contrast, CENP-E intensity was not significantly altered (**Appendix Fig. S8A-D**), although its kinetochore signal volume showed a modest increase in MPS1-overexpressing cells (**Appendix Fig. S8E**). Collectively, these data show that MPS1 overexpression causes inappropriate retention of fibrous corona components (core RZZ:Spindly and the microtubule-binding protein CENP-F) on aligned metaphase kinetochores, consistent with delayed or incomplete corona disassembly and suggesting an architectural basis for the increased merotelic attachments observed under elevated MPS1 activity.

**Figure 6.**
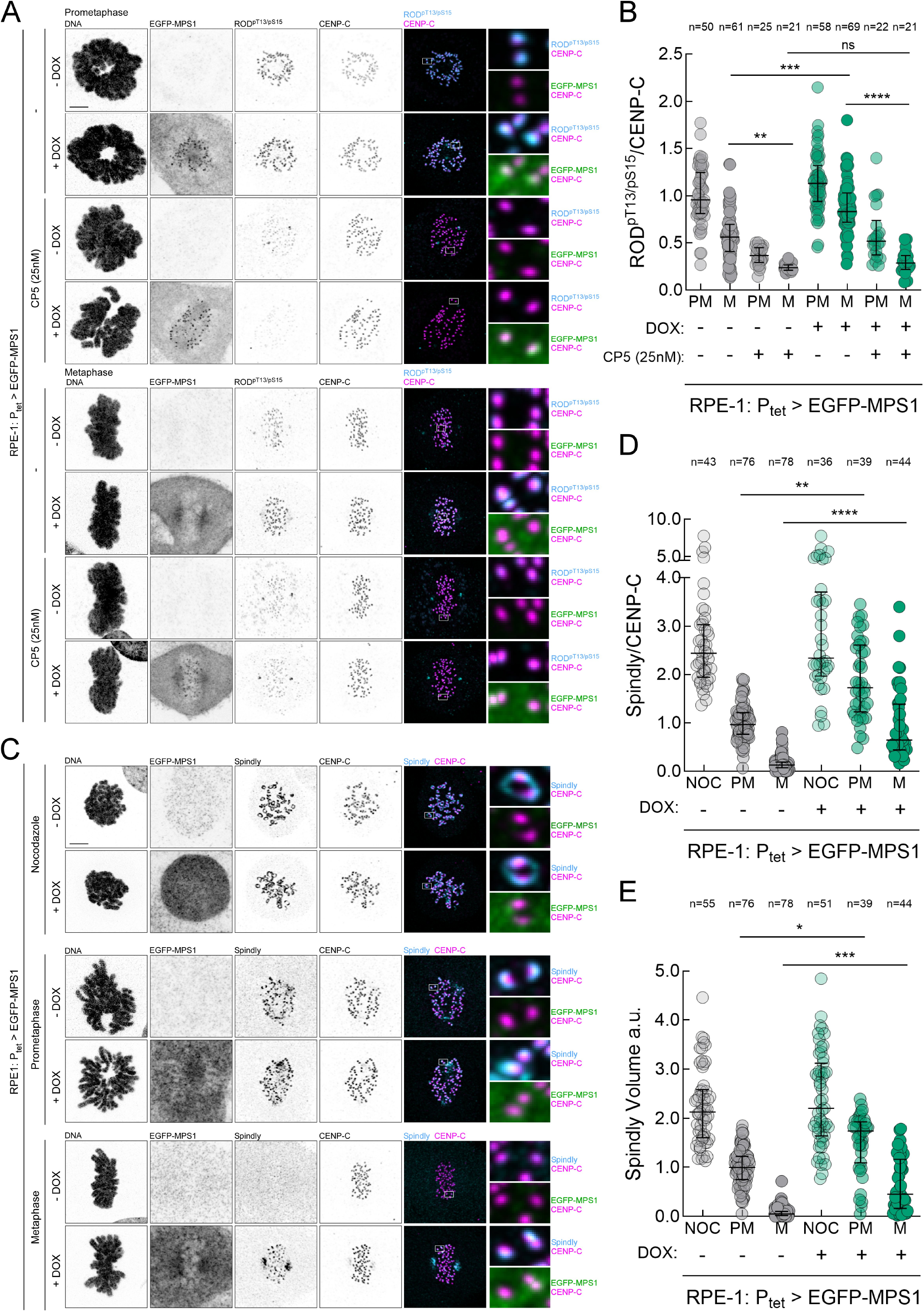
MPS1 overexpression sustains signalling for fibrous corona assembly at metaphase kinetochores. (A-B) Representative immunofluorescence images **(A)** and corresponding quantifications **(B)** of ROD phosphorylation at Thr13 and Ser15 (ROD^pT13/pS15^) levels at kinetochores of RPE-1 cells in control (-DOX) and MPS1 overexpression (+DOX) conditions, in prometaphase (PM) and metaphase (M). MPS1 was partially inhibited when indicated with CP5. Insets depict magnifications of selected kinetochores. ROD^pT13/pS15^ fluorescence intensities were determined relative to the CENP-C signal. All values in **(B)** were normalized to the mean value determined for prometaphase control RPE-1 cells, which was set to 1. Each data point represents the mean of an individual cell; n=4 independent experiments. **(C-E)** Representative immunofluorescence images **(C)** and corresponding quantifications of Spindly levels **(D)** and Spindly volumes **(E)** at kinetochores of RPE-1 cells in control (-DOX) and MPS1 overexpression (+DOX) conditions, in nocodazole (NOC), prometaphase (PM) and metaphase (M) Insets depict magnifications of selected kinetochores. Spindly fluorescence intensities were determined relative to the CENP-C signal. All values in **(D)** and **(E)** were normalized to the mean value determined for prometaphase control RPE-1 cells, which was set to 1. Each data point represents the mean of an individual cell; n=3 independent experiments. Data information: data in **(B)**, **(D)** and **(E)** are presented as medians with interquartile range; asterisks indicate that differences between mean ranks are statistically significant; \**p* < 0.05; \*\**p* < 0.01; \*\*\**p* < 0.001; \*\*\*\**p* < 0.0001; ns, non-significant; Kruskal-Wallis, Dunn’s multiple comparison test in (B), (D) and (E). Scale bars: 5μm.

### MPS1 overexpression sustains a fibrous corona at mature end-on attachments, enabling merotelic kinetochore–microtubule engagement

To determine whether the fibrous corona persists at mature end-on attachments and how it interacts with microtubules under MPS1 overexpression, we examined Spindly co-localization with Astrin and used Stimulated Emission Depletion (STED) microscopy to visualize microtubule engagement at Spindly-positive kinetochore– microtubule interfaces. In metaphase RPE-1 cells expressing endogenous MPS1 (−DOX), the vast majority of Astrin-positive kinetochores showed little or no Spindly signal (**Fig. EV4A,B**). This is consistent with normal fibrous corona dynamics, in which formation of robust end-on attachments is coupled to corona disassembly/stripping (Sacristan et al., 2018). By contrast, in MPS1-overexpressing cells (+DOX) the fraction of Astrin-positive kinetochores retaining Spindly was markedly increased (**Fig. EV4A,C**). Per-cell quantification revealed a high proportion of Astrin-positive kinetochores co-labelled with Spindly, indicating that Spindly, and by extension the fibrous corona, persists on kinetochores that have already formed mature end-on attachments under elevated MPS1 activity.

Using STED microscopy, we next examined the spatial relationship between Spindly, the inner kinetochore marker ACA, and kMTs in metaphase cells. In amphitelic attachments of both control (-DOX) and MPS1-overexpressing (+DOX) cells, k-fibers terminated at ACA-positive kinetochores, consistent with bona fide end-on attachments (**Fig. 7A**). Strikingly, however, under MPS1 overexpression, Spindly frequently formed an external domain aligned with the incoming microtubule bundle and positioned distal to ACA (**Fig. 7B**). Three-dimensional STED and rendered reconstructions revealed k-fibers extending through the Spindly-positive outer domain before terminating at the ACA-labelled kinetochore core (**Fig. EV4D** and **Movie EV23, EV24**). Together, these observations indicate that when MPS1 activity is abnormally elevated at metaphase, mature end-on attachments can still form while corona material remains retained. Notably, 3D STED imaging frequently revealed a “split-interface” attachment geometry in MPS1-overexpressing metaphase cells, in which a single kinetochore simultaneously engaged microtubules from both spindle poles through distinct kinetochore domains (**Fig. 7C,D**). In these events, the k-fiber from one pole formed a canonical end-on attachment at the kinetochore core, with microtubule tips terminating at the ACA-labelled inner kinetochore. In parallel, a microtubule bundle originating from the opposite pole terminated within the retained Spindly-positive domain, consistent with microtubule engagement of corona meshwork rather than the kinetochore core (**Fig. EV4E** and **Movie EV25, EV26**). Thus, persistence of an expanded fibrous corona provides an additional microtubule-binding interface that can capture microtubules from the opposing pole, offering a direct structural explanation for the elevated merotelic attachment frequency under MPS1 overexpression. Consistent with opposite pole-directed pulling forces acting on a single kinetochore, these merotelic configurations were accompanied by deformation of the ACA-defined kinetochore core and an apparent inward displacement of the Spindly signal relative to ACA (**Fig. EV4E** and **Movie EV25, EV26**).

**Figure 7.**
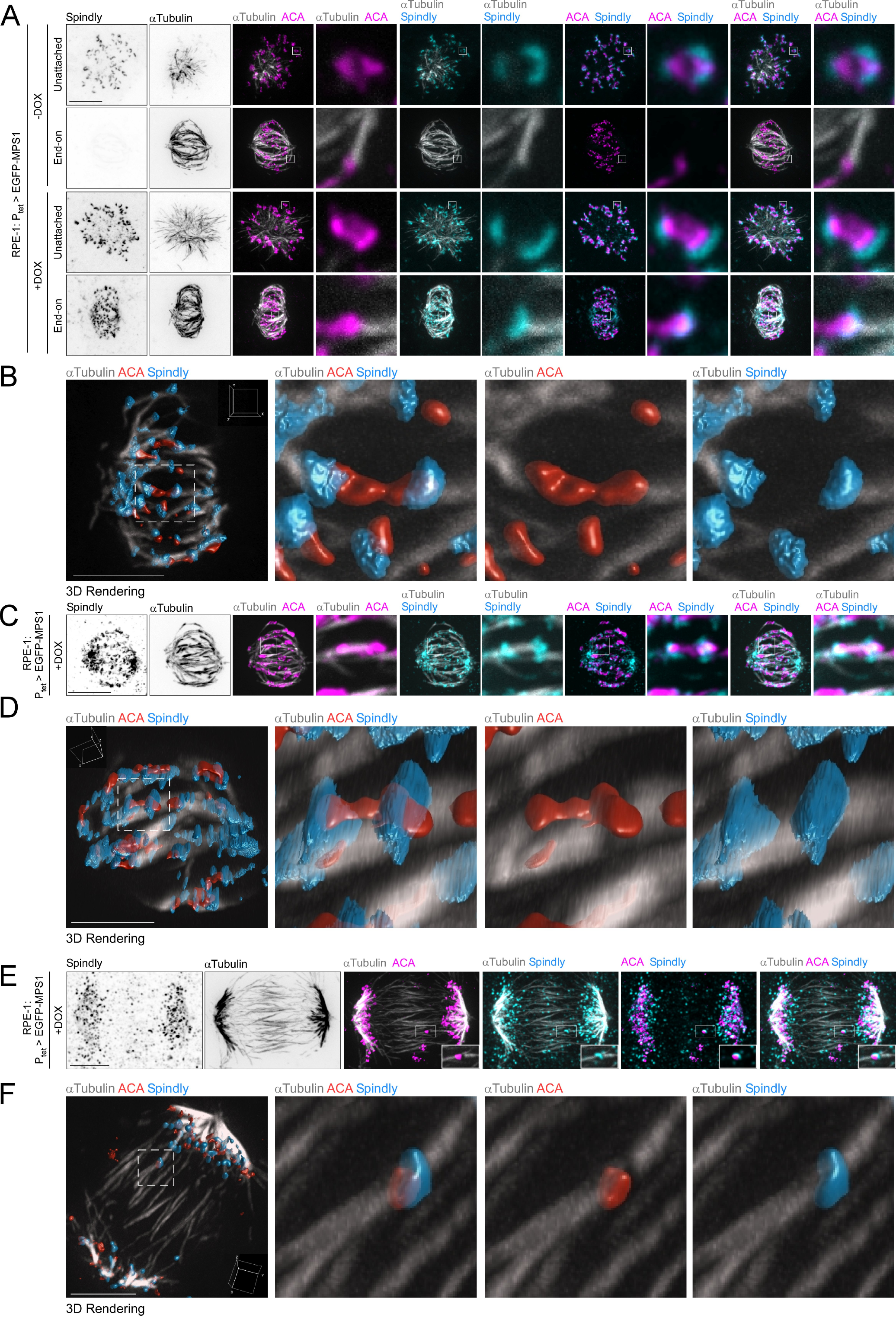
MPS1 overexpression maintains fibrous corona assembly at end-on-attached kinetochores, promoting merotelic microtubule capture. (A) Representative STED microscopy images of prometaphase and metaphase of RPE-1 cells in control (-DOX) and MPS1 overexpression conditions (+DOX). Microtubules (grey; STED), Spindly (cyan; STED) and centromeres (ACA; magenta; confocal). Insets depict magnifications of selected kinetochores. Scale bar: 5 µm. **(B)** Three-dimensional reconstruction of the metaphase RPE-1 cell overexpressing MPS1 shown in **(A)** (selected Z-slices). Insets depict magnifications of selected sister-kinetochores in amphitelic attachment configuration. Microtubules (grey), Spindly (blue) and centromeres (ACA; red). Scale bar: 5 µm. **(C, D)** Representative STED microscopy images **(C)** and respective three-dimensional reconstruction (selected Z-slices, different perspective) **(D)** of a merotelic kinetochore-microtubule attachment in a metaphase RPE-1 cell overexpressing MPS1 (+DOX). Microtubules (grey; STED), Spindly (cyan; STED) and centromeres (ACA; magenta; confocal). Insets depict magnifications of selected kinetochores. Scale bar: 5 µm. **(E, F)** Representative STED microscopy images **(E)** and respective three-dimensional reconstruction (selected Z-slices, different perspective) **(F)** of a merotelic kinetochore-microtubule attachment in an anaphase RPE-1 cell overexpressing MPS1 (+DOX). Microtubules (grey; STED), Spindly (cyan; STED) and centromeres (ACA; magenta; confocal). Insets depict magnifications of selected kinetochores. Scale bar: 5 µm.

The same “split-interface” merotelic geometry was readily detected at lagging chromosomes of MPS1-overexpressing cells undergoing anaphase, with one microtubule bundle terminating within the retained Spindly-positive domain and microtubules from the opposite spindle pole terminating at the ACA-labelled inner kinetochore core (**Fig. 7E,F; Fig. EV4F** and **Movie EV27, EV28**). Consistent with substantial mechanical strain at these merotelic kinetochores, the inner kinetochore marker CENP-C displayed an elongated/stretched organization specifically at Spindly-positive lagging kinetochores, indicative of kinetochore deformation under opposing pole-directed forces (**Fig. EV4G,H**).

Together, these observations provide direct structural evidence that fibrous corona retention under elevated MPS1 activity creates an additional microtubule-engagement interface capable of capturing microtubules from the opposing pole, thereby generating merotelic attachments that can persist into anaphase and give rise to lagging chromatids.

### Preventing fibrous corona assembly at metaphase kinetochores restores anaphase fidelity in RPE-1 cells overexpressing MPS1

Having established that MPS1 overexpression sustains fibrous corona material at metaphase kinetochores and promotes merotelic engagement, we next tested whether persistent corona assembly is required for the elevated lagging-chromosome phenotype. We reasoned that if ongoing corona maintenance underlies merotely in MPS1-overexpressing cells, then suppressing corona persistence at metaphase should restore chromosome segregation fidelity in anaphase. We first used a pharmacological strategy to reduce MPS1 activity toward control levels by treating MPS1-overexpressing (+DOX) cells with a suboptimal dose of the MPS1 inhibitor CP5. Consistent with sustained corona assembly being driven by continued MPS1-mediated phosphorylation of ROD (**Fig. 6A,B**), partial MPS1 inhibition restored near-normal Spindly dynamics. Spindly was efficiently cleared from aligned metaphase kinetochores in MPS1-overexpressing cells treated with CP5, resembling the pattern observed in controls expressing endogenous (-DOX) MPS1 (**Fig. 8A-C** and **Fig. EV5A,B**). Importantly, this restoration of corona clearance upon normalizing MPS1 activity was accompanied by a significant reduction in the frequency of anaphases with lagging chromosomes (**Fig. 8D,E** and **Movies EV29-EV31**).

**Figure 8.**
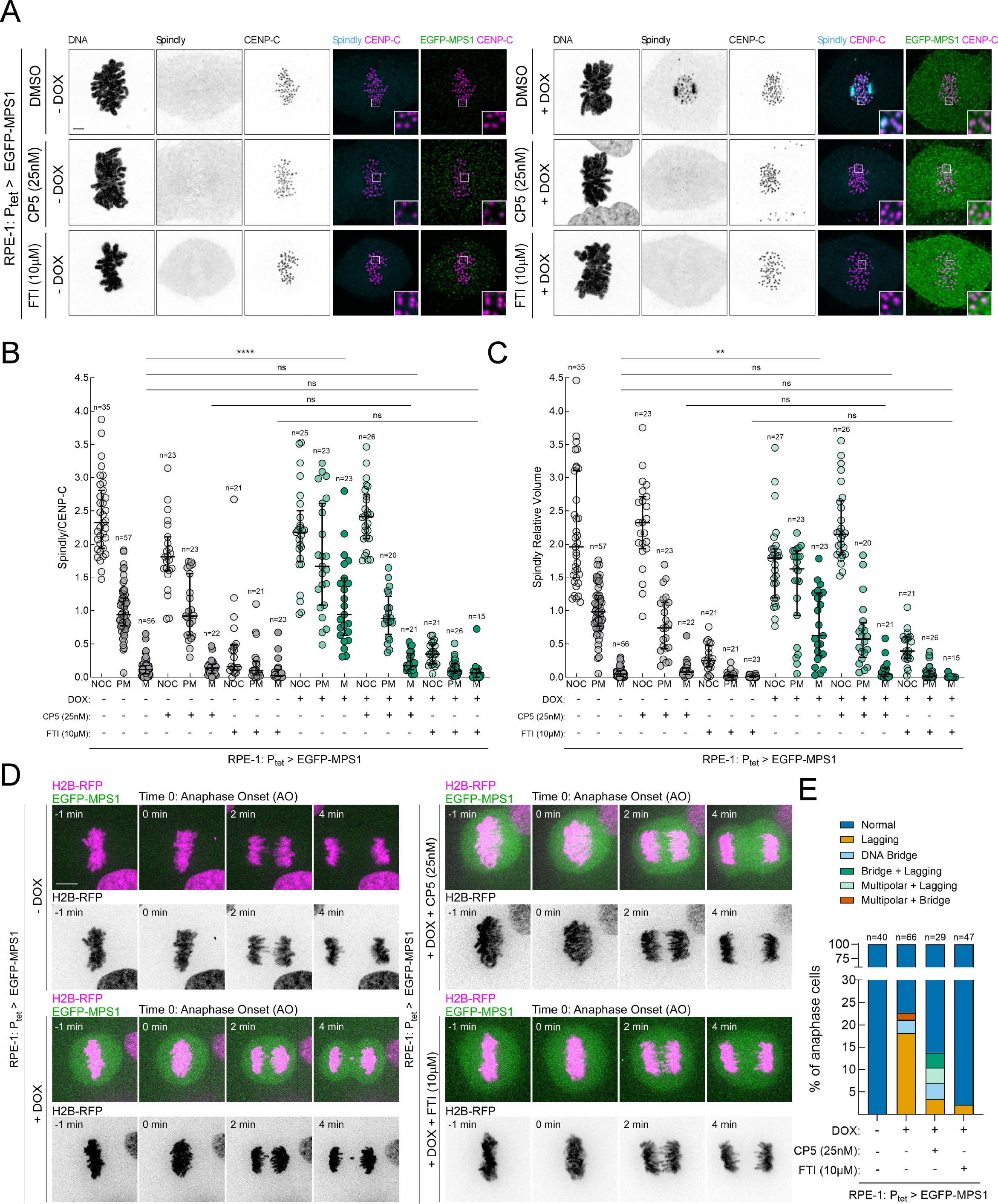
Preventing abnormal accumulation of Spindly at metaphase kinetochores restores anaphase fidelity in RPE-1 cells overexpressing MPS1. (A) Representative immunofluorescence images of Spindly kinetochore localization at metaphase in RPE-1 control (-DOX) and MPS1 overexpressing (+DOX) cells. MPS1 was partially inhibited when indicated with CP5. Spindly farnesylation was prevented when indicated with FTI-277. Insets depict magnifications of selected kinetochores. **(B,C)** Quantification of Spindly levels **(B)** and volumes **(C)** at kinetochores of RPE-1 cells in control (-DOX) and MPS1 overexpression (+DOX) conditions, in nocodazole (NOC), prometaphase (PM) and metaphase (M). MPS1 was partially inhibited when indicated with CP5. Spindly farnesylation was prevented when indicated with FTI-277. Spindly fluorescence intensities were determined relative to the CENP-C signal. All values in **(B)** and **(C)** were normalized to the mean value determined for prometaphase control RPE-1 cells, which was set to 1. Each data point represents the mean of an individual cell; n≥2 independent experiments. Scale bar: 5μm. See also **Fig. EV5A**. **(D)** Selected still frames from representative live-cell imaging of mitotic progression of RPE-1 cells stably expressing H2B-RFP (magenta) in control (-DOX) and MPS1 overexpression (+DOX) conditions. EGFP-MPS1 overexpression is depicted in green. Time (min) is shown relative to anaphase onset (AO). MPS1 was partially inhibited when indicated with CP5. Spindly farnesylation was prevented when indicated with FTI-277. Scale bar: 5μm. **(E)** Quantification of chromosome segregation fidelity for the cell lines and conditions shown in **(D)**. Graph bars represent the frequency of indicated anaphase output. n denotes the number of anaphase events analysed (filmed) for each condition. Data information: data in **(B)** and **(C)** are presented as medians with interquartile range; asterisks indicate that differences between mean ranks are statistically significant; \*\**p* < 0.01; \*\*\*\**p* < 0.0001; ns, non-significant; Kruskal-Wallis, Dunn’s multiple comparison test.

We next disrupted corona assembly independently by preventing Spindly kinetochore recruitment using the farnesyl-transferase inhibitor FTI-277, which blocks Spindly farnesylation required for its kinetochore targeting and corona assembly (Moudgil et al., 2015; Holland et al., 2015; Raisch et al., 2022). Accordingly, FTI-277 abrogated kinetochore localization of Spindly and suppressed fibrous corona formation in both control (-DOX) and MPS1-overexpressing (+DOX) RPE-1 cells (**Fig. 8A-C** and **Fig. EV5A,B**). As expected from the role of the corona in the initial microtubule capture and chromosome congression, FTI-277 treatment prolonged prometaphase, with cells requiring additional time to align their chromosomes at the metaphase plate and achieve biorientation (**Fig. EV5C,D**). Strikingly, however, once alignment was achieved, cells were not delayed in metaphase relative to their respective untreated controls (−FTI) and proceeded into anaphase with high segregation fidelity, resulting in near-complete suppression of lagging chromosomes and other anaphase defects in MPS1-overexpressing cells (**Fig. 8D,E**; **Fig. EV5C,D** and **Movie EV32**). To exclude off-target effects of FTI-277, we independently impaired corona assembly by siRNA-mediated depletion of Spindly, which similarly delayed chromosome congression but restored segregation fidelity in MPS1-overexpressing cells (**Appendix Fig. S9A-C** and **Movies EV33-EV35**).

Together, these orthogonal interventions demonstrate that sustained fibrous corona assembly is required for the chromosome segregation errors induced by MPS1 overexpression. These findings support a model in which elevated MPS1 maintains fibrous corona material on aligned metaphase kinetochores, expanding the microtubule-interaction interface and favouring merotelic attachment formation that can persist into anaphase to cause lagging chromatids.

### Preventing fibrous corona assembly improves the fidelity of chromosome segregation in chromosomally unstable PCCCs

Next, we asked whether the mechanism uncovered in engineered RPE-1 cells could explain the strong association between MPS1 overexpression and CIN in colon cancer by testing whether fibrous corona persistence contributes to lagging chromosomes in chromosomally unstable (CIN^+^) PCCCs. We first quantified Spindly kinetochore dynamics across mitotic progression. In chromosomally stable RPE-1 cells and the near-diploid colon MSI line JVE059, kinetochore Spindly levels declined sharply from prometaphase to metaphase, consistent with normal fibrous corona clearance upon biorientation (**Fig. 9A** and **Appendix Fig. S10**). By contrast, a substantial fraction of kinetochores in all three CIN^+^ PCCC lines retained Spindly at metaphase (**Fig. 9A** and **Appendix Fig. S10**). Co-staining with Astrin confirmed that Spindly persists on kinetochores bearing mature end-on attachments in CIN^+^ PCCCs (**Fig. 9A,B**), indicating delayed or incomplete fibrous corona disassembly despite attachment maturation.

**Figure 9.**
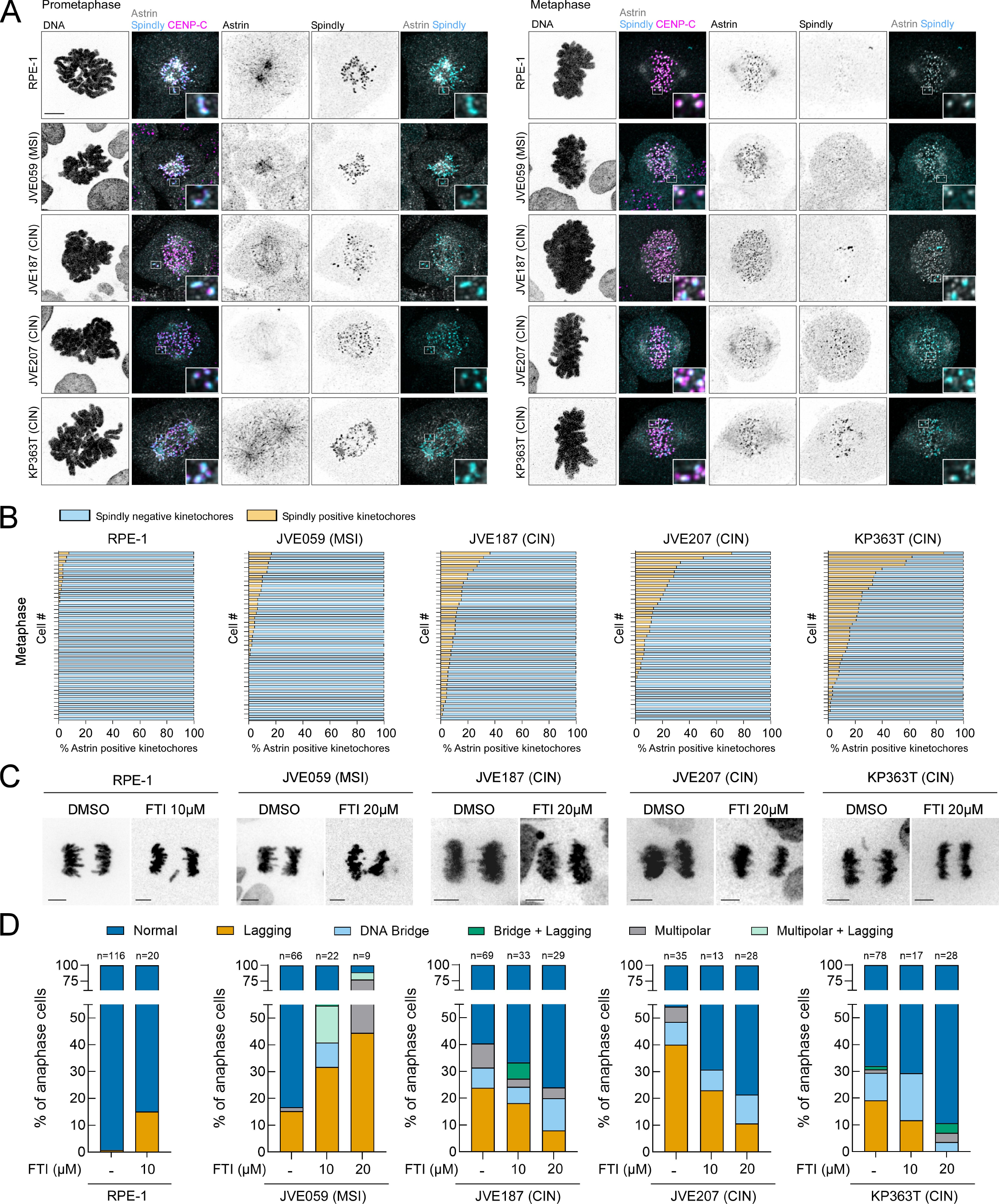
Preventing fibrous corona assembly improves the fidelity of chromosome segregation in chromosomally unstable PCCCs. (A) Representative immunofluorescence images of Spindly and Astrin localization at prometaphase and metaphase kinetochores in RPE-1 cells and PCCCs JVE059 (MSI), JVE187 (CIN), JVE207 (CIN) and KP363T (CIN). CENP-C was used as kinetochore reference. Insets depict magnifications of selected kinetochores. **(B)** Histograms depicting the percentage of Astrin-positive metaphase kinetochores per cell that are also positive (orange bars) or negative (blue bars) for Spindly in the cell lines indicated in **(A)**. Each bar represents one individual cell; n=2 independent experiments. **(C)** Representative still frames from live-cell imaging movies of chromosome segregation in RPE-1 cells and PCCCs, under control conditions (DMSO) and following treatment with the farnesyl-transferase inhibitor FTI-277 (FTI). Chromosomes (visualized by H2B-RFP) are represented in inverted greyscale. **(D)** Quantification of chromosome segregation fidelity for the cell lines and conditions indicated. Graph bars represent the frequency of indicated anaphase output. n denotes the number of anaphase events analysed (filmed) for each condition. Data information: data in **(B)** and **(D)** are plotted as fractions of the total. Scale bar: 5μm.

We next tested whether metaphase Spindly retention in CIN^+^ PCCCs depends on MPS1 activity. Partial inhibition of MPS1 with CP5 significantly reduced Spindly levels at metaphase kinetochores across all three CIN^+^ PCCC lines (**Appendix Fig. S11A,B**), restoring the fraction of Astrin-positive kinetochores that co-retained Spindly to near-control levels (**Appendix Fig. S11C**). Independently, blocking fibrous corona assembly with the farnesyl-transferase inhibitor FTI-277 suppressed kinetochore Spindly already in prometaphase across RPE-1, JVE059, and CIN^+^ PCCC lines (**Appendix Fig. S12A-C**), and by metaphase, Spindly was largely absent from kinetochores in all FTI-treated conditions, including CIN^+^ PCCCs (**Appendix Fig. S12A-C**). Importantly, live-cell imaging of CIN^+^ PCCC lines revealed that FTI-277 treatment markedly reduced the frequency of lagging chromosomes during anaphase (**Fig. 9C,D**). Together, these data demonstrate that CIN^+^ PCCCs inappropriately retain fibrous corona material at metaphase kinetochores in an MPS1-dependent manner, and that preventing corona assembly significantly suppresses lagging chromosomes. These findings directly link metaphase fibrous corona persistence to chromosome mis-segregation in patient-derived CIN colon cancer cells, providing a mechanistic explanation for the association between elevated MPS1 activity and CIN in colon carcinomas.

## DISCUSSION

Chromosomal instability (CIN) is a major source of intratumour heterogeneity and tumour evolution, yet the proximal molecular events that sustain recurrent chromosome mis-segregation remain poorly defined in most cancers. This gap is particularly evident in colon cancer, where the CIN pathway accounts for the majority of MSS tumours but direct mechanistic links between canonical colon cancer driver mutations and recurrent chromosome mis-segregation have been difficult to establish (Lengauer et al., 1997; Shih et al., 2001; Hermsen et al., 2002; Giaretti et al., 2004a; Giaretti et al., 2004b; Trautmann et al., 2006; Pino and Chung, 2010). Here, we propose a model in which transcriptional rewiring of mitotic control, rather than recurrent mutation of the core mitotic machinery, establishes a CIN-permissive state in colon cancer (**Fig. 1A-C** and **Appendix Fig. S1A**). Our cohort-level analyses revealed a prominent upregulation of *AURKA*, *SKA3* and *MPS1/TTK* genes in CIN^+^ MSS tumours (**Fig. 1D**). MPS1 is a regulator of both Aurora A (Leça et al., 2025) and the SKA complex (Maciejowski et al., 2017) and therefore represents an upstream node capable of modulating multiple pathways that control kinetochore–microtubule attachment dynamics and error correction. Beyond this central mitotic positioning, MPS1 is broadly upregulated across human cancers and has been proposed to exhibit cancer-germline features, consistent with its classification among genes whose expression is normally enriched in the germline but aberrantly reactivated in tumours (cancer-testis/cancer-germline genes) (Singh et al., 2021; Nin and Deng, 2023). Moreover, MPS1 is included in the CIN70 chromosomal instability expression signature, a gene set constructed around transcripts whose overexpression correlates with aneuploidy and CIN (Carter et al., 2006; Slee et al., 2014). Finally, prior work further suggests that aneuploid cells that overexpress MPS1 tend to become dependent on its activity to survive high mitotic stress (Daniel et al., 2011; Maire et al., 2013; Slee et al., 2014; Maia et al., 2015; Zhang et al., 2016; Choi et al., 2017; Anderhub et al., 2019; Chandler et al., 2020). Collectively, these observations support a close relationship between elevated MPS1 and CIN. However, what has remained unresolved, is whether elevated MPS1 is simply a compensatory adaptation to pre-existing CIN (for example, to bolster SAC signalling in a stressed mitotic environment) or whether it can actively drive the errors that generate and sustain CIN. Our study provides evidence for the latter by showing that MPS1 overexpression disrupts the temporal remodelling of the outer-kinetochore architecture by continuously promoting fibrous corona assembly.

Fibrous corona assembly is normally restricted to the period in which unattached kinetochores require an enlarged surface for microtubule capture and checkpoint signalling. As biorientation is established, declining MPS1 activity, reduced ROD phosphorylation and Dynein- and phosphatase-dependent clearance dismantle this capture platform, leaving the core NDC80-based interface to support load-bearing end-on attachment (Griffis et al., 2007; Gassmann et al., 2010; Sacristan et al., 2018; Rodriguez-Rodriguez et al., 2018; Barbosa et al., 2020; d’Amico et al., 2022; Raisch et al., 2022; Eibes et al., 2023; Cmentowski et al., 2023; Ide et al., 2023). Elevated MPS1 shifts this balance by maintaining ROD phosphorylation at Thr13 and Ser15, thereby prolonging an assembly-competent RZZ:Spindly state beyond its normal window (**Fig. 6**; **Fig. 9A,B and Fig. EV4**). The consequence is an uncoupling of two events that are normally coordinated. The core kinetochore acquires a mature end-on attachment while an outer corona-based capture surface remains available.

This uncoupling provides a structural explanation for merotely. A retained corona creates a second microtubule-engagement domain spatially distinct from the canonical kinetochore core, allowing one microtubule bundle to terminate at the end- on attachment interface while microtubules from the opposite pole engage the corona (**Fig. 7A-D** and **Fig. EV4A-E**). Such “split-interface” attachments explain how chromosomes that appear aligned and carry mature end-on attachments at metaphase can nevertheless become merotelic and lag during anaphase. Consistent with this model, the same split-interface geometry is detected at lagging chromosomes, accompanied by deformation of inner-kinetochore markers suggestive of opposing poleward forces acting on a single kinetochore (**Fig. 7E,F** and **Fig. EV4F-H**). The suppression of lagging chromosomes when corona assembly is reduced, either by partially inhibiting MPS1 to restore corona clearance or by preventing Spindly kinetochore recruitment, supports a causal role for corona persistence rather than a mere association with elevated MPS1 (**Fig. 8**; **Fig. 9** and **Appendix Fig. S9**). Thus, MPS1 overexpression provides a direct mechanistic link between tumour-associated transcriptional rewiring and the chromosome segregation errors that sustain CIN by maintaining an otherwise transient corona-based microtubule-capture surface at metaphase. This increases the likelihood of forming merotelic attachments that persist into anaphase and generate lagging chromatids.

The molecular basis through which the retained corona supports load-bearing microtubule engagement remains however to be defined. CENP-F is a plausible contributor because it contains two microtubule-binding regions, interacts with Dynein-regulatory modules and is capable of high-affinity microtubule binding (Feng et al., 2006; Musinipally et al., 2013; Volkov et al., 2015; Auckland et al., 2020). CENP-E, whose kinetochore-associated volume increased at metaphase despite only modest changes in signal intensity, may also cooperate with CENP-F and other microtubule-interacting factors embedded within the multivalent RZZ:Spindly meshwork. By concentrating several individually weak or transient microtubule-binding activities, and orienting them along the incoming k-fiber, an expanded corona could generate sufficient avidity to stabilise otherwise short-lived contacts and convert them into a mechanically competent, load-bearing interface. Thus, microtubule engagement by the persistent corona may emerge from cooperative multivalent binding rather than from the activity of any single corona component. Direct perturbation of CENP-F, CENP-E and defined elements of the RZZ:Spindly architecture will be required to establish which components bear force and which primarily organise the platform.

How can MPS1, which normally promotes error correction by destabilising improper attachments, permit robust end-on attachments when overexpressed? Our findings suggest that MPS1 output is not defined simply by kinase abundance. Instead, it is likely determined by its spatial positioning, substrate accessibility and the local kinase– phosphatase balance. Under physiological conditions, MPS1 is enriched at unattached or incorrectly attached kinetochores, where it promotes error-correction pathways through regulation of the SKA complex, Aurora B targeting and Aurora A activation (Jelluma et al., 2008b; van der Waal et al., 2012; Maciejowski et al., 2017; Hayward et al., 2022; Leça et al., 2025). As biorientation is established, kinetochore-associated MPS1 declines and phosphatase activity promotes attachment stabilisation and SAC silencing (Nijenhuis et al., 2014; Moura et al., 2017; Hayward et al., 2022). However, when MPS1 is overexpressed, our results show that an active kinetochore-associated pool persists into metaphase without causing global destabilisation of end-on attachments (**Fig. 5**; **Appendix Fig. S3 and Appendix Fig. S5**). Instead, elevated MPS1 sustains an expanded fibrous corona that acts as an additional microtubule-engagement surface. This apparent selectivity can be explained in several, non-mutually exclusive ways: (i) metaphase kinetochores may spatially restrict the excess kinase from the key destabilizing substrates within the core attachment interface; (ii) the metaphase-retained MPS1 pool may be functionally biased toward the fibrous corona branch, where it efficiently phosphorylates ROD to sustain RZZ:Spindly oligomerization, rather than continuing to engage SKA/Aurora-dependent destabilization pathways; (iii) metaphase phosphatases may preferentially oppose MPS1’s destabilization outputs at the core kinetochore while leaving the ROD/corona-assembly pathway comparatively less antagonized when MPS1 recruitment persists; and (iv) the accompanying metaphase delay imposed by MPS1 overexpression may further allow end-on attachments to mature despite sustained MPS1 activity. Thus, we propose that MPS1 overexpression creates a substrate-biased signalling state that uncouples two processes normally coordinated during biorientation: stabilization of the core end-on attachment interface and disassembly of the outer fibrous corona. This uncoupling explains how MPS1, although required for error correction at physiological levels, can promote stable merotelic attachments when overexpressed.

The coexistence of robust end-on attachments with an expanded fibrous corona under MPS1 overexpression presents however a second apparent paradox, as the RZZ complex has previously been shown to inhibit NDC80-dependent end-on binding (Cheerambathur et al., 2013; Amin et al., 2018). We suggest this is reconciled by viewing RZZ as a context-dependent regulator of attachment timing and maturation rather than as an absolute inhibitor of the NDC80 complex. In early mitosis, the RZZ:Spindly corona favours lateral microtubule capture and delays premature stabilisation of end-on attachments, thereby gating the transition to biorientation. Once a load-bearing end-on attachment is established, however, NDC80 engagement, tension-dependent kinetochore remodelling and recruitment of maturation factors such as Astrin may render the core interface comparatively resistant to RZZ activity. Consistent with this interpretation, our 3D STED reconstructions show that the retained corona appears to form an external, permeable domain through which k-fibres can pass before terminating at the kinetochore core, rather than physically occluding microtubule access to NDC80 (**Fig. 7** and **Fig. EV4**). Moreover, the metaphase-retained corona may be compositionally or conformationally distinct from its early-mitotic counterpart and therefore preserve microtubule-capture capacity without fully maintaining its inhibitory effect on end-on attachment. Finally, biorientation itself imposes a strongly pro-attachment regime (increased microtubule occupancy, tension-dependent kinetochore architecture, and maturation of end-on binding), which may overcome any residual kinetic or steric penalty imposed by a partially retained corona. Thus, rather than universally preventing end-on connections, RZZ/corona appears to modulate when and how end-on attachments mature. MPS1 overexpression disrupts this temporal coordination, allowing the core attachment interface to mature while preserving an outer corona-based capture surface that increases susceptibility to merotely.

MPS1-driven persistence of the metaphase fibrous corona seems to be sufficient to convert otherwise karyotypically stable, p53-proficient RPE-1 and JVE059 cells into a CIN state characterised by recurrent merotelic lagging chromosomes, micronucleus formation and progressive karyotypic divergence (**Fig. 4**; **Fig. EV2** and **Appendix Fig. S4**). This is consistent with evidence that mild numerical aneuploidies can be propagated despite intact p53 signalling (Santaguida et al., 2017; Soto et al., 2017; Hintzen et al., 2022). Notably, however, chronic MPS1 overexpression also produced structural aneuploidies (translocations), which are generally thought to be more strongly constrained in p53-proficient cells (Santaguida et al., 2017; Soto et al., 2017; Hintzen et al., 2022). A possible reconciling explanation is that p53 imposes a stringent but non-absolute barrier whose effectiveness depends on how and when structural lesions arise and on their cellular cost. In line with that, it should be noted that p53-proficient RPE-1 cells stably carry the well-documented structural rearrangement der(X)t(Xq;10q) involving duplicated 10q material translocated onto chromosome X (Volpe et al., 2025). Likewise, the near-diploid colorectal cancer cell line HCT116, which retains wild-type p53, can propagate stable structural lesions, including a 10q+ marker and translocations involving chromosomes 10/16, and 8/16 depending on the specific clonal population, demonstrating that structurally altered karyotypes can be compatible with robust proliferation in a p53-proficient setting (Masramon et al., 2000; Abdel-Rahman et al, 2001; Roschke et al., 2002). We therefore hypothesise that the structural abnormalities detected after MPS1 overexpression arise through secondary genome evolution driven by repeated micronucleus formation and DNA damage, rather than through the immediate propagation of acutely generated segmental lesions. Chronic, moderate CIN may additionally select for relatively tolerable configurations that evade a sustained p53 response, allowing rare surviving clones to expand over time. In this view, p53 strongly restricts the propagation of many segregation-error-derived structural abnormalities but does not absolutely prevent the retention of lesions whose origin, magnitude and fitness cost are compatible with long-term proliferation.

Anaphase lagging chromosomes arising from erroneous merotelic attachments are viewed as major contributors to genomic instability in human cancer, in part because, even if many lagging events are transient or ultimately corrected, a subset of lagging chromosomes fails to be incorporated into daughter nuclei and generate micronuclei (Cimini et al., 2002; Cimini et al., 2004; Thompson and Compton 2011; Orr et al., 2021; Sen et al.,2021). Micronuclei are of particular interest because they have been implicated as key intermediates of chromothripsis and other catastrophic genome-rearrangement processes, such as breakage–fusion–bridge cycles and chromoanasynthesis, that can accelerate tumour evolution, drug resistance, and oncogene activation (Crasta et al., 2012; Hatch et al., 2013; Zhang et al., 2015; Mackenzie et al., 2017; Bakhoum et al., 2018; Ly et al., 2019; Umbreit et al., 2020; Krupina et al., 2021). Recent work has shown that micronuclei can also originate from chronically misaligned chromosomes that silence the SAC and mispartition directly (Gomes et al., 2022; Tucker et al., 2023; Koprivec et al., 2026). Together with our findings, these studies support a model in which the relative contributions of “misaligned-chromosome” versus “lagging-chromosome” micronuclei are genetically and mechanistically determined. Perturbations that compromise congression and alignment mechanics, such as defects in CENP-E/KIF18A-dependent pathways, bias toward micronuclei seeded by misaligned chromosomes, whereas perturbations that expand or inappropriately retain the fibrous corona, such as MPS1 overexpression, bias toward persistent, load-bearing merotelic attachments in otherwise aligned metaphase chromosomes, producing lagging chromatids that are prone to micronucleus formation. Viewed together, these data argue that cancer cells can exploit multiple, mechanistically distinct routes to generate micronuclei and sustain CIN, and that micronucleus origin should be considered an informative readout of the underlying mitotic failure mode with potential consequences for the genomic and signalling outcomes of micronucleated progeny.

Several questions remain open. It is presently unclear whether lagging chromosomes under elevated MPS1 arise preferentially from chromosomes that are intrinsically prone to fibrous corona retention and merotelic attachment, or whether corona persistence and the ensuing merotely occur stochastically. This issue is underscored by the heterogeneous retention of metaphase corona material in both MPS1-overexpressing cells and CIN+ PCCCs (**Fig. 9A,B**; **Fig. EV4A-C** and **Appendix Fig. S10**). Among kinetochores bearing mature end-on attachments, only a subset retains detectable Spindly. Such mosaicism could reflect chromosome-intrinsic determinants of corona persistence, including kinetochore size and centromere architecture, or a probabilistic process in which local attachment history, geometry or microtubule–kinetochore force balance transiently biases individual kinetochores towards a corona-retained state.

In RPE-1 cells, MPS1 overexpression promotes merotelic attachments and lagging chromosomes without globally increasing kinetochore–microtubule stability (**Fig. 5D,E**), indicating that persistent corona-mediated microtubule capture is sufficient to generate segregation errors in an otherwise stable near-diploid background. By contrast, CIN+ PCCCs exhibit hyperstable kinetochore–microtubule attachments (**Appendix Fig. S2E**), a common feature of chromosomally unstable cancer cells. These defects may therefore cooperate in colon tumour cells to promote chromosome mis-segregation: persistence of the fibrous corona may increase the probability of merotelic attachments, whereas hyperstable attachments may favour their persistence into anaphase. Whether this mechanism operates across CIN malignancies remains unknown. A converse phenotype was recently reported in several breast cancer cell lines, where reduced kinetochore localisation of ROD, ZW10 and Zwilch, was accompanied by attenuated corona-dependent microtubule nucleation and lower MPS1 activity (Ishikawa et al., 2025). Corona dysregulation in cancer may therefore be context-dependent and bidirectional: insufficient corona function before attachment may impair microtubule capture, whereas sustained assembly signalling after end-on attachment may preserve an ectopic microtubule-engagement surface that promotes merotely. MPS1-driven corona persistence should thus be viewed as a defined route to CIN in MPS1-high tumours, rather than a universal cancer phenotype. This context dependence may reflect broader rewiring of the mitotic machinery. *AURKA* and *SKA3* are also strongly upregulated in CIN+ MSS tumours. Defining whether and how Aurora A and SKA3 cooperate with MPS1 to tune attachment stability, corona dynamics and tolerance of segregation errors will further refine the mitotic circuitry that sustains CIN *in vivo*.

Overall, we demonstrate that excessive MPS1 activity drives CIN by promoting untimely expansion of the fibrous corona in metaphase, providing a structural basis for merotely and chromosome mis-segregation in colon cancer. This work therefore positions MPS1 as a molecular link between the oncogenic landscape of colon cancer and the mitotic errors that define CIN, raising the possibility that tumour cells exploit MPS1 overexpression not merely as a byproduct of transformation, but as an active mechanism to generate the chromosomal diversity required for adaptation and clonal evolution. These findings not only advance our understanding of the mechanisms driving CIN but also underscore the potential significance of targeting MPS1 and the fibrous corona as therapeutic strategies in colon cancer.

## METHODS

### Atlas and Compass of Immune-CAncer-Microbiome Interactions in Colon Cancer (AC-ICAM)

Genomic and transcriptomic data for the AC-ICAM colon cancer cohort were obtained from the study by Roelands et al., 2023, in which fresh-frozen primary colon tumours and matched normal colon tissue were collected and profiled by RNA-seq and whole-exome sequencing (WES), and WES-derived copy-number profiles were generated. All laboratory procedures, sequencing, primary processing, and quality control were performed as described in Roelands et al. (2023). For the gene mutation and expression analyses, we included the subset of 281 tumours that passed the original study QC for tumour–normal concordance and contamination, as defined by Roelands et al., 2023. Chromosomal instability (CIN) status was inferred from the WES-derived genome-wide copy-number alteration (CNA) burden. CIN^+^ and CIN^−^ designations were taken directly from the AC-ICAM CNA burden-based classification reported in Roelands et al., 2023. For the expression analysis of mitotic genes, RNA-seq gene expression values were obtained from the AC-ICAM processed expression matrix generated and normalised in Roelands et al., 2023 (EDASeq within-lane and between-lane normalisation followed by quantile normalisation). Expression of 44 mitotic genes was analysed in MSS tumours and compared between CIN^+^ and CIN^−^ groups. For each gene, the log2 fold change in expression (CIN^+^ relative to CIN^−^) was calculated and plotted.

### Somatic mutation analysis of colon tumours

Somatic mutations were analysed for a predefined panel of 44 mitotic genes using the AC-ICAM (Roelands et al., 2023) and The Cancer Genome Atlas (TCGA) Pan-Cancer Atlas colorectal adenocarcinoma (TCGA COAD-READ) datasets available through cBioPortal (v6.3.5; https://www.cbioportal.org). Patients were stratified into microsatellite instability (MSI) and microsatellite stability (MSS) groups. For the TCGA COAD-READ dataset, the classification was based on two complementary MSI scoring metrics, the MANTIS score (> 0.6, MS; 0.4< indeterminate < 0.6; < 0.4, MSS) and the MSIsensor score (> 10, MSI and < 10, MSS). Samples with discordant classifications between the two metrics or with missing data were excluded from the analysis. For the AC-ICAM dataset, the MANTIS score was used to classify samples either as MSI-H (MANTIS score > 0.4) or MSS (MANTIS score ≤ 0.4) (Roelands et al., 2023). Mutations were classified into five primary categories: missense, truncating, inframe, splice, and fusion. Combinations of these categories were additionally recorded to account for samples harbouring multiple mutation types within the same gene. The frequency of each mutation type per gene was calculated by dividing the number of samples harbouring that mutation type by the total number of samples in each group (MSI or MSS).

### Survival analysis

Overall survival (OS) and progression-free survival (PFS) were analysed in the AC-ICAM colon cancer cohort using Kaplan–Meier estimation and Cox proportional hazards regression. Tumours were stratified by microsatellite instability status (MSI-H vs MSS) and by chromosomal instability (CIN) status, which was inferred from GISTIC data by summing up the total event of amplifications or deletion (shallow and deep) per sample. A cutoff of <5000 for CIN^-^ and >5000 CIN^+^ was used based on the distribution of values (inspection of density curve). To minimise confounding from the mismatch-repair–driven hypermutator phenotype and distinct tumour–immune biology of MSI-H cases, all statistical comparisons of CIN-associated outcome were restricted to MSS tumours, comparing MSS CIN^+^ versus MSS CIN^−^. Kaplan-Meier plots were generated using ggsurvplot from R package survminer (v.0.4.9). Hazard ratios (HRs) and the corresponding 95% confidence intervals (95% CI) and P values were estimated using a Cox proportional hazards model implemented in the R package survival (v2.41–3), with CIN status as the sole predictor (univariable Cox model). Cox regression was performed only when each comparison group contained at least 10 patients. Censoring was handled by standard right-censoring at last follow-up.

The R2:Genomics Analysis and Visualisation Platform (http://r2.amc.nl) was used to analyse the overall survival and disease-free survival (DFS) in AC-ICAM and GSE17538 (Smith et al., 2010) cohorts after stratifying patients by tumour stage (stage III and stage IV) and by MPS1 expression. For each stage, patients were dichotomised into MPS1-high and MPS1-low groups using the expression cutoff indicated on the plots (selected with a minimum group size constraint of min.grp = 8). Survival analyses were performed with R2’s KaplanScanner tool, after evaluating every possible expression threshold as a potential cutoff to divide the cohort into high- and low-expression groups. For each threshold, a log-rank test was applied to compare survival outcomes, and the cutoff that produces the most statistically significant separation (lowest p-value) was selected. The resulting Kaplan–Meier curves were then plotted accordingly. The overall P value comparing survival between expression-defined groups was computed using a two-sided log-rank test.

### Cell lines and cell culture conditions

Patient-derived colon cancer cells JVE059, JVE187, JVE207 and KP363T were previously established (Boot et al., 2016). In brief, anonymized tumour material was extensively rinsed in RPMI-1640 medium under vigorous tapping. Tissue was subsequently minced into approximately 1 mm³ fragments which were enzymatically dissociated using a solution containing 1% collagenase I-A (Sigma) and 1% dispase (Gibco Life Technologies). The resulting dissociated cells were washed with RPMI-1640, and primary cultures were initiated in DMEM/F12 supplemented with 10% fetal bovine serum (FBS), 100 U/mL penicillin, and 100 μg/mL streptomycin. All anonymized human samples were handled in accordance with the medical ethical guidelines described in the Code Proper Secondary Use of Human Tissue established by the Dutch Federation of Medical Sciences.

All cell lines, including hTERT RPE-1 (American Type Culture Collection) were grown in DMEM-F12, HEPES (Invitrogen, Cat. No. 31330038) supplemented with 10% fetal bovine serum (FBS; Life Technologies, Cat. No. A5256801) and penicillin/streptomycin (100 IU/mL and 100 µg/mL; Life Technologies, Cat. No. 15140-122). Cells were kept at 37°C in a humidified incubator with 5% CO_2_ and were routinely tested for mycoplasma contamination using a mycoplasma-specific PCR assay with the primers set: MGSO: 5’-TGC ACC ATGTGTCACTCTGTTAACCTC – 3’ and GPO1: 5’-ACTCCTACGGGAGGC AGCAGT A – 3’.

### Exome and transcriptome analysis of PCCC lines

Genomic and transcriptomic data of PCCC lines were obtained from the study by Boot et al., 2016. Exome profiles of PCCC lines were obtained by hybridizing DNA to Infinium HumanExome-12v1 BeadChips (Boot et al., 2016). DNA isolation was performed using the Wizard Genomic DNA Purification Kit (Promega, Madison, WI, USA) according to the manufacturer’s instructions. BeadChips were used with an input of 200 ng DNA. Copy number profiles and group copy number analysis was performed using the DNAcopy package (Venkatraman et al., 2007).

Transcriptomic profiles of PCCC lines were obtained by hybridizing cDNA to Infinium HumanExome-12v1 BeadChips (Boot et al., 2016). RNA isolation was performed using TRIzol® Reagent (Life Technologies). DNAse treatment was performed in suspension using rDNAse (Macherey Nagel GmbH & Co. KG, Düren, Germany) and 500 ng of RNA was converted to cDNA using the DyNAmo™ cDNA Synthesis Kit (Thermo Scientific, Waltham, MA, USA). cDNA was then purified using the QIAquick PCR purification kit (Qiagen, Germantown, Maryland, USA) and eluted in 15 μL MQ water. Five μL of purified cDNA was used as input for the Infinium protocol. Gene expression data was generated using the intensity signal of the cDNA hybridization on the BeadChips. Intronic probes and probes which were heterozygous in any sample were removed. Subsequently, intensity data related to the colour of the genotyped allele was extracted for each probe. After quantile normalisation using the Limma package (Ritchie et al., 2015) the average probe intensity per gene was calculated and gene expression was reported in log2 expression values per gene per sample. In total 17090 genes were assayed. Raw data and pre-processed intensities per probe are available via the Gene Expression Omnibus (GEO) under accession numbers GSE67773 and GSE67774.

### Drug treatments

When required, cultured cells were subjected to several drug treatments before being collected and processed for the desired analysis. Microtubule depolymerization was induced by treatment with 3.3 μM nocodazole (Sigma-Aldrich, CAS 31430-18-9) for 4 hours prior fixation for immunofluorescence experiments, or for 2 hours prior live-cell imaging acquisition. To induce MPS1 overexpression, 1µg/mL of doxycycline (Sigma-Aldrich, D9891-1G) was added to the cells 48h before the experiment. For MPS1 partial inhibition, 25nM or 35nM of CP5 (Koch et al., 2016; gift from René Medema, Oncode Institute, The Netherlands Cancer Institute, The Netherlands), was added to the cell culture media 2.5h before fixation. For farnesylation inhibition, 10 µM or 20 µM of FTI-277 (Merck Life Science S.L.U., F9803-1MG) was added to the cell culture media 48h before fixation. For monastrol washout experiments, cells were treated with 100 µM monastrol (Tocris, Bioscience, Cat. No.1305) for 14h. After this period, cells were washed three times with pre-warmed PBS to remove monastrol and then incubated for 60 min at 37 °C in pre-warmed culture medium supplemented with 5 µM MG132 (Sigma-Aldrich; CAS 133407-82-6), prior to fixation and processing for immunostaining. For all live-cell experiments, when required, drugs were added directly into the imaging medium and remained during the experiment.

### siRNA-mediated depletion of Spindly

For siRNA-mediated knockdown, cells were transfected with 40 nM siRNA targeting Spindly transcripts using Lipofectamine RNAiMAX (Thermo Fisher Scientific) in Opti-MEM (Thermo Fisher Scientific) and incubated for 48 h prior live-cell imaging. The siRNA sequences were: siSPINDLY, 5′-GAAAGGGUCUCAAACUGAA-3′, and a non-targeting control siRNA, 5′-UGGUUUACAUGUCGACUAA-3′ (Sigma-Aldrich).

### Transgene constructs

Plasmids pLKO.1-LV-H2B-RFP (Addgene #26001, Beronja et al., 2010), pRRL-CMV-EGFP-αTubulin, mRFP-αTubulin and pLVx-PA-GFP–αTubulin (Ganem et al., 2005) were previously generated. The pLIX_403 backbone plasmid was obtained from Addgene (#41395). Linear vector and insert fragments were generated by PCR using the following primer pairs: pLIX_403 FC Fw (5’-CTTATACACAGCCAGTCTGCAGGTC-3’) and pLIX_403 FC Rv (5’-CCTTAGCTCCTGAAAATCTCGACGG-3’) for linearisation of the pLIX_403 backbone, and EGFP-MPS1 pLIX FC Fw (5’-CCGTCGAGATTTTCAATGGTGAGCAAGGG-3’) and EGFP-MPS1 pLIX FC Rv (5’-CTGGCTGTGTATAAGACAAGCTTGGTACCGCAT-3’) for amplification of the EGFP-MPS1 construct. PCR products were purified using the PCR Clean up DNA extraction from agarose gels kit (Macherey-Nagel). The expression vector pLIX-EGFP-MPS1 was then assembled by circular polymerase extension cloning (CPEC) (Quan & Tian, 2009). To generate the expression vector pLVx-EGFP-MPS1, the EGFP-MPS1 cDNA was amplified by PCR and inserted into the pLVx-PA-GFP–αTubulin construct by FastCloning (Li et al., 2011), replacing the previously removed PA-GFP–αTubulin.

### Lentiviral transduction

All lentiviral particles were produced by co-transfection of HEK293T cells with the lentiviral vectors, psPAX2 (Gag, Pol, Rev, and Tat expressing packaging vector), pMD2.G (VSV-G expressing envelope vector) and with plasmids bearing the gene of interest: pLKO.1-LV-H2B-RFP (Addgene #26001, Beronja et al., 2010), pRRL-CMV-EGFP-αTubulin, pLVx-PA-GFP–α-Tubulin, pLIX-EGFP-MPS1 and pLVx-EGFP-MPS1. After 72h, supernatants containing lentiviral particles were collected. For lentiviral transduction, lentiviral particles were added to RPE-1 and PCCC cell lines with DMEM 10% FBS with 1:1000 Polybrene (Sigma-Aldrich) for 24h. The lentiviruses were used individually, giving time for cells to recover between transductions. Stable lines with sufficient fluorescence intensity were selected by FACS sorting.

### RT-qPCR

The RNA isolation and purification was performed using the RNA isolation kit, NucleoSpin® RNA from Macherey-Nagel according to the manufacturer’s instructions. Concentration measurements were performed using the nanodrop device. Reverse transcriptase reaction was performed with 1 µg of total RNA with random hexamers and oligo(dT) primers. RT-qPCR was performed using IQ Sybr. Master mix (Bio-Rad) on a CFX 96 Real-Time PCR System (Bio-Rad Laboratories) following standard cycling conditions. Primer sequences used for MPS1 RT–qPCR were as follows: qPCR_MPS1_Forw 5’-GATTCTCAGGTTGGCACAGTT-3’ and qPCR_MPS1_Rev 5’-CATCCTAAGGACCAAACATCACT-3’. Relative gene expression levels were calculated using the ΔΔCt method (Livak & Schmittgen, 2001) and normalized to the housekeeping gene CPSF6. The results were analysed using CFX Manager, Microsoft Excel and GraphPad Prism v10.0.

### Cell lysates and Western blotting

For the preparation of cell lysates to measure protein levels, cells were collected after trypsinization and centrifuged at 1200 rpm for 5 min at 4°C. After washing the pellets in PBS 1×, cells were pelleted by centrifugation at 1200 rpm for 5 min at 4°C. The pellets were then resuspended in 50 μL of lysis buffer (50 mM Tris HCl pH 7.4; 150 mM NaCl; 1 mM EDTA; 0.5% Triton; 0.1% Digitonin; 10 mM MgCl_2_; 30 µg/mL RNase; 20 µg/mL DNase; supplemented with 1× protease inhibitor cocktail (Roche, Basel, Switzerland) and 1× phosphatase inhibitor cocktail (Sigma-Aldrich, St. Louis, MO, USA)) and kept on ice for 30 min, before immersion in liquid nitrogen for cell disruption. Protein extracts in the supernatant were recovered by centrifugation at 14000 rpm for 10 min at 4°C and protein concentration determined by Bradford protein assay (Bio-Rad, Hercules, CA, USA). Samples were resuspended in Laemmli sample buffer and heated at 95°C for 5 min before being resolved in SDS–PAGE and probed for proteins of interest through Western blotting. For Western blotting analysis, resolved proteins were transferred to a nitrocellulose membrane, using the iBlot Dry Blotting System (Thermo Fisher Scientific) according to the manufacturer’s instructions. Membranes were incubated for at least 1h at room temperature in blocking solution (5% powder milk in PBS1×, 0.05% Tween 20). All primary and secondary antibodies were diluted in the blocking solution. Membranes were incubated with primary antibody solutions overnight at 4°C under constant stirring and washed three times in PBS1×, 0.05% Tween 20 for 10 min each. Then, membranes were incubated with secondary antibody conjugated to HRP (Santa Cruz Biotechnology) solutions for 1h at room temperature under constant stirring. Blots were developed with ECL Chemiluminescent Detection System (Amersham) according to the manufacturer’s protocol and detected on x-ray film (Fuji Medical).

### Photoactivation experiments

Microtubule turnover rates were measured in cells stably expressing mRFP-αTubulin and PA–GFP–αTubulin. Cells were cultured on glass-bottom dishes (MatTek) coated with Fibronectin (25 μg/mL; F1141-1MG; Sigma-Aldrich) using DMEM-F12, HEPES, without phenol red (Invitrogen, Cat. No. 11039047) supplemented with 10% fetal bovine serum (FBS; Life Technologies, Cat. No. A5256801) and penicillin/streptomycin (100 IU/mL and 100 µg/mL; Life Technologies, Cat. No. 15140-122). Timelapse imaging was performed in a heated chamber (37°C) using a 100× 1.4 NA plan-apochromatic differential interference contrast (DIC) objective mounted on an inverted microscope (Nikon TE2000U) equipped with a CSU-X1 spinning-disk confocal head (Yokogawa Corporation of America) and with two laser lines (488 nm and 561 nm). Images were detected with an iXonEM+ EM-CCD camera (Andor Technology). Photoactivation was performed with a Mosaic digital mirror device– based patterning system (Andor) equipped with a 405-nm diode laser. Cells in prometaphase and metaphase were selected using DIC acquisition. A region of interest corresponding to spindle microtubules for photoactivation was selected using a line segment placed perpendicular to the main axis in one of the sides adjacent to the metaphase plate. The region of interest was locally photoactivated by pulsed near-UV irradiation (405-nm laser; 500-ms exposure time) and fluorescence images (seven 1-µm separated z-planes centered at the middle of the mitotic spindle) were captured every 15 s for 4.5 min with a 100× oil-immersion 1.4 NA planapochromatic objective. To quantify fluorescence dissipation after photoactivation, spindle poles were aligned horizontally, and whole-spindle, sum-projected kymographs were generated as previously described (Pereira & Maiato, 2010). Fluorescence intensities were quantified for each time point (custom-written routine in Matlab, “LAPSO” software) (Girão & Maiato, 2020) and normalized to the first time point after photoactivation for each cell following background subtraction and correction for photobleaching. Correction for photobleaching was performed by normalizing to the values of fluorescence loss obtained from whole-cell (including the cytoplasm), sum-projected images for each individual cell. This method allows for the precise measurement of photobleaching for each cell at the individual level. Under these conditions, the photoactivated region dissipates, yet the photoactivated molecules are retained within the cellular boundaries defined by the cytoplasm. To calculate microtubule turnover, the sum intensity at each time point was fit to a double exponential curve A1*exp(−k1*t) + A2*exp(−k2*t) using Matlab (Mathworks), in which t is time, A1 represents the less stable microtubule population (non-kMT), and A2 the more stable microtubule population (kMT) with decay rates of k1 and k2, respectively. When running the routine to fit the data points to the model curve, the rate constants are obtained as well as the percentage of microtubules for the fast (typically interpreted as the fraction corresponding to non-kMT) and the slow (typically interpreted as the fraction corresponding to kMT) processes. The half-life for each process was calculated as ln2/k for each population of microtubules. All experiments were performed in the presence of MG132 (5µM), which was added 30 min before imaging to ensure that cells were in metaphase and prevent mitotic exit.

### Immunofluorescence analysis

For the immunofluorescence experiments, cells were seeded 48h or 72h in coverslips coated with fibronectin (25 μg/mL; F1141-1MG; Sigma-Aldrich). Unless specified, cells were fixed with 4% paraformaldehyde in PTEM buffer (20 mM PIPES pH 6.8, 0.2% Triton X-100, 10 mM EGTA, 1 mM MgCl2) for 10 minutes at room temperature, following a wash with PBS 1x. For assessing Aurora B pT232, cells were fixed for 10 minutes at room temperature in 4% paraformaldehyde following an extraction with 0.3% Triton X-100 in PBS for 5 min. To determine MAD1 levels at kinetochores, cells were pre-extracted with 0.1% Triton X-100 in PEM (100 mM Pipes, pH 6.8, 1 mM MgCl_2_ and 5 mM EGTA) for 1 min and then fixed in 4% paraformaldehyde for 10 minutes. For immunofluorescence with the Spindly antibody, cells were fixed in 4% paraformaldehyde in PHEM buffer (60 mM PIPES pH 6.8, 25 mM HEPES, 10 mM EGTA, 2 mM MgCl_2_) for 7 min 30 sec, followed by an extraction with 0.1% Triton X-100 in PHEM for 2 min. Cells were washed 1x with PHEM and 1x with PBS 1x. For assessing MPS1 pT676, cells were fixed in 4% paraformaldehyde in PHEM buffer (60 mM PIPES pH 6.8, 25 mM HEPES, 10 mM EGTA, 2 mM MgCl_2_) containing 0.1% Triton X-100 for 12 min. Cells were washed 1x with PBS 1x. Blocking was performed in 1× PBS, 10% fetal bovine serum for 1h. Primary antibody incubations were prepared in blocking solution overnight at 4°C, followed by three 5-min washes in 1× PBS. Secondary antibody incubations were performed as described for the primary antibodies, including the three 5-min washes at the end. Slides were then mounted using Vectashield mounting medium for fluorescence with DAPI (Vector Laboratories, Burlingame, CA). To analyse kinetochore microtubule attachments, cells were subjected to a calcium treatment for 2.5 min with 100 mM PIPES pH 6.8, 1 mM MgCl_2_, 1 mM CaCl_2_, 0.5% Triton X-100. Cells were fixed with 0.1% glutaraldehyde and 4% paraformaldehyde (Electron Microscopy Sciences) for 10 min. Autofluorescence was quenched by 0.1% sodium borohydride (Sigma-Aldrich) after aldehyde fixation and cells permeabilized with 0.5% Triton X-100 (Sigma-Aldrich) for another 10 min. Blocking was performed in PBS 0.05% Triton X-100, 10% fetal bovine serum for 1h. Primary antibody incubations were prepared in blocking solution overnight at 4°C, followed by three 5-min washes in PBS 0.05% Triton X-100. Secondary antibody incubations were performed as described for the primary antibodies, including the three 5-min washes at the end. Slides were mounted as described above. Images were collected in a Zeiss Axio Imager microscope (Carl Zeiss) or a Leica TCS SP8 laser scanning confocal microscope (Leica Microsystems, Germany). 3D deconvolution was performed using Huygens Professional (Scientific Volume Imaging) employing the CMLE algorithm (100 iterations; quality threshold 0.1) and a theoretical point spread function computed from the acquisition metadata. For quantification of fluorescence intensities, an ImageJ macro was used as previously described (Saurin et al., 2011). Briefly, kinetochores were identified based on constitutive marker (CENP-C or ACA) and segmented using the threshold function. The segmentation mask was enlarged by one pixel to ensure complete kinetochore selection and subsequently applied across all channels to quantify the mean fluorescence intensity of each kinetochore protein of interest. This approach yielded the mean pixel intensity for all kinetochores within a single cell. Background fluorescence was determined by thresholding the lowest 5% of signal intensity values across the image using the setThreshold function. Thresholds were held constant within an experiment. In cases where staining produced a substantial cytoplasmic background signal, background was quantified from empirically defined ROI placed within the cytoplasmic compartment, excluding kinetochore regions. The intensities of the kinetochore proteins were quantified relative to CENP-C fluorescence. Control values were averaged and used for normalisation of values determined in the different biological conditions tested.

For quantification of kinetochore-microtubule attachment configurations (end-on, lateral and unattached), kinetochore positions were manually identified based on CENP-C staining from maximum projected raw images. To verify kinetochore-microtubule attachment, the corresponding raw images were analysed in a z-stack-by-stack basis and for each previously defined kinetochore region of interest, the presence or absence of associated microtubules was determined. End-on attachment was defined as a kinetochore directly engaged with the plus-end of a k-fibre. Lateral attachment was assigned when a kinetochore was associated with the lateral surface of the microtubule lattice rather than the plus-end. Kinetochores were classified as unattached when no detectable association with microtubules was observed.

Co-localization of Spindly and Astrin at kinetochores was quantified based on kinetochore segmentation using CENP-C as a reference marker. Individual kinetochores were identified and segmented from the CENP-C channel using Fiji software (https://fiji.sc/), and fluorescence intensities for Spindly and Astrin were measured at each segmented kinetochore region. Kinetochores were classified as positive or negative for each marker using intensity thresholds defined empirically based on background signal and marker-specific reference levels: prometaphase signal was used as the reference for Spindly, reflecting its maximal recruitment at unattached kinetochores, whereas metaphase signal was used as the reference for Astrin, reflecting its maximal recruitment at bi-oriented, end-on attached kinetochores.

Thresholds were applied consistently across all images and conditions. For each cell, the proportion of kinetochores positive for Spindly alone, or positive for both Spindly and Astrin, was calculated relative to the total number of segmented kinetochores.

### Volumes determination at kinetochores

Raw TIFF image stacks were converted to IMS format using Imaris v10.2.0 (Oxford Instruments). Volumetric quantification of corona protein signal at kinetochores was performed using the automated batch processing function with pre-defined parameters applied uniformly across all images and experimental conditions. Nuclear boundaries were defined using DAPI fluorescence, and corona protein surfaces were segmented within the nuclear compartment. Segmentation thresholds were empirically determined to maximise the corona protein signal overlapping spatially with CENP-C, while minimising background, and were kept constant across all conditions. A 3D DAPI mask was applied to exclude non-specific signal outside the nuclear volume. Segmented surfaces smaller than 0.05 μm³ were discarded to eliminate background signal, and positional filters in the XY plane were applied to restrict segmentation to the kinetochore-occupied nuclear region, as confirmed by CENP-C signal. The total corona protein surface volume per cell was normalised to the number of kinetochores, determined by manual counting of CENP-C foci using Fiji software (https://fiji.sc/).

### STED Microscopy and 3D rendering analysis

Cells were fixed using PFA (Electron Microscopy Sciences) in Cytoskeleton Buffer (CB) pH 6.1 (274 mM NaCl, 10 mM KCl, 2.2 mM Na_2_HPO_4_, 0.8 mM KH_2_PO_4_, 4 mM EGTA, 10 mM PIPES, 4 mM MgCl_2_, 10 mM glucose) for 10 min. For autofluorescence quenching it was used 0.1% sodium borohydride solution (Sigma-Aldrich) diluted in PBS, for 10 min. Extraction was performed with CB-0.5%Triton (Sigma-Aldrich), for 10 min. Blocking was performed for 1h at RT with cytoskeleton buffer 10% FBS and 0.05% Tween 20. Primary antibodies used: human anti-centromere antiserum at 1:1000 (ACA; Fitzgerald), chicken anti-GFP at 1:2000 (1:2000; ab13970; Abcam), rabbit anti-spindly at 1:2500 (Arshad Desai OD:174), mouse anti-α-tubulin clone B-512 at 1:3000 (SigmaAldrich) diluted in cytoskeleton buffer with 10% FBS and 0.05% Tween 20. Secondary antibodies used were anti-rabbit STAR 580 (Abberior Instruments); anti-mouse STAR 647 (Abberior Instruments) at 1:100; Alexa Fluor anti-chicken 488 (Invitrogen) at 1:2000 and Alexa Fluor anti-human 405 (Invitrogen) at 1:1000, diluted in CB-0.1% Tween with 10% FBS. Coverslips were mounted using mounting medium. For STED imaging, an Inverted microscope Leica Stellaris 8 with STED and Falcon module equipped with a DMI8 motorized inverted microscope (Leica Microsystems, Germany), with a HC PL APO 93x/1,30 GLYC motCORR STED WHITE (working distance: 300um) objective lens. All acquisition channels (confocal and STED) were performed using a 1 Airy unit pinhole. Fixed cell images were acquired using excitation wavelengths at 405nm, 488nm, 561nm and 640nm with a collection window of 412-479, 494-571, 593-624 and 642-750, respectively. Excited volumes were doughnut-depleted with a single laser at 775 nm set at 30% and adjusted at 20% AOTF. All images in the figures represent maximum intensity projections of the entire cell (unless specified otherwise) collected with a z-stack of 0.214µm. Quantifications and panel construction were performed with Fiji software (https://fiji.sc/).

Three-dimensional rendering of the STED image stacks was performed using Huygens Professional v25.10 (Scientific Volume Imaging, The Netherlands). The ACA channel (405 nm excitation, confocal), was deconvolved using the Classic Maximum Likelihood Estimation (CMLE) algorithm (100 iterations; quality threshold 0.1) with a theoretical point spread function computed from the acquisition metadata. For visualization of Spindly and ACA, the Huygens Surface Renderer was employed. Surface reconstruction was performed independently for each channel, generating two separate segmentation groups. Initial intensity thresholds and seed values were suggested automatically by the software and subsequently adjusted empirically by comparison with the raw STED image to optimally capture the signal of each marker. ACA and Spindly surfaces were rendered in red and blue, respectively.

To visualize microtubules, the tubulin channel was first rendered using the Huygens MIP (Maximum Intensity Projection) Renderer (displayed in white), providing an overview of the microtubule bundle architecture within the region of interest. To further resolve the 3D spatial relationship between tubulin and each kinetochore-associated protein, a third segmentation group was generated for the tubulin channel using the Surface Renderer (displayed in yellow), with threshold and seed values independently tuned to best represent the tubulin signal. This tertiary surface reconstruction enabled direct visualization of the interface between the microtubule bundle and ACA or Spindly.

To comprehensively assess the relative positioning between ACA, Spindly, and the microtubule bundle, all renderings were performed on a cropped kinetochore– microtubule attachment region of interest. Rendered images were captured at multiple viewing angles by rotating the 3D volume, allowing visualization of the spatial relationships from different orientations.

### Live-cell imaging

For live cell imaging, human cells were cultured on glass-bottom dishes (MatTek) coated with Fibronectin (25 μg/mL; F1141-1MG; Sigma-Aldrich) using DMEM-F12, HEPES, without phenol red (Invitrogen, Cat. No. 11039047) supplemented with 10% fetal bovine serum (FBS; Life Technologies, Cat. No. A5256801) and penicillin/streptomycin (100 IU/mL and 100 µg/mL; Life Technologies, Cat. No. 15140-122). 4D datasets were collected using spinning disc confocal systems: Andor Revolution XD (Andor Technology, Oxford Instruments, UK) equipped with an inverted motorized IX81 microscope (Olympus, Japan), a EMCCD iXonEM+ DU-897 camera (Andor Technology, Oxford Instruments, UK), a CSU-22 confocal scanner unit (Yokogawa Electric Corporation, Japan, and a 488 and a 561 nm laser line, driven by iQ software (Andor Technology, Oxford Instruments, UK). EGFP was excited using the 488 nm laser line in combination with a 488 dichroic mirror and a 488-emission filter. mRFP was excited using the 561 nm laser line, together with a Semrock dual 488/561 dichroic mirror and a Semrock dual 488/561 emission filter. Images were acquired with a 40x /1.00 oil immersion objective. Andor BC43 unit (Andor Technology, Oxford Instruments, UK), equipped with a 40x/0.95 Plan Apochromat (INS-OBJ-40D-095), a 60x/1.42 Plan Apochromat oil immersion (INS-OBJ-60D-142-O), as well as 488 nm and a 561 nm laser lines. EGFP was excited using the 488 nm laser line in combination with a quad dichroic mirror (405/488/561/640) and a 488 emission filter. mRFP was excited using the 561 nm laser line, together with a quad dichroic mirror (405/488/561/640) and a 561-emission filter. Images were acquired using a 2040x2040 pixel format at 16-bit depth. The system was controlled using Fusion software (version 2.7), and image data were saved in .ims format.

Time-lapse imaging of z stacks with 1-µm steps covering the entire volume of the cell were collected on both microscopes. All imaging was performed under controlled environmental conditions (37 °C, 5 % CO₂), using constant laser power and exposure time settings within each experiment.

For the experiments involving siRNA-mediated depletion and corresponding controls, imaging was performed on an inverted Zeiss Axio Observer Z1 inverted microscope (Marianas Imaging Workstation, 3i – Intelligent Imaging Innovations Inc.) equipped with a CSU-X1 spinning-disk confocal head (Yokogawa Corporation of America) and four laser lines (405, 488, 561, and 640 nm). A Plan-Apochromat DIC 63×/1.4 NA oil immersion objective was used, and imaging was conducted in an environmental chamber maintained at 37 °C with controlled humidity and 5% CO₂. Image acquisition was performed using an iXon Ultra 888 EM-CCD camera (Andor Technology) and SlideBook 6.0.24 software (3i – Intelligent Imaging Innovations, Inc.). RPE-1 cells were seeded in 35 mm glass-bottom dishes (MatTek) and imaged at 2-minute intervals. Z-stacks were acquired with 1 µm step size to capture the full extent of the mitotic spindle.

### Multicolor-FISH Karyotyping of RPE-1 cells

Cells were processed for mFISH karyotyping adapting the protocol described by Trott et al, 2017. RPE-1 cell cultures were grown to 80% confluency in 10cm dishes, then incubated in 100 ng/mL Colcemid for 3h. Cell media was collected in a falcon tube, cells were washed once with PBS. Trypsin-EDTA diluted 1:1 in PBS was added to the cells following an incubation at 37°C for 1.5 min. A mitotic shake-off was performed to collect the mitotic cells. Cells were centrifuged 5 minutes at 200g. The supernatant was aspirated and the cells were resuspended dropwise in 5mL of 75mM KCl hypotonic solution pre-warmed at 37°C with low power vortex and incubated on water bath at 37°C for 15 min. Carnoy fixation solution was added dropwise and the cells were centrifuged 200g 5min. The cells were fixed with 5mL Carnoy fixation solution added dropwise on low power vortex and stored at -20°C. Cell suspensions were applied onto glass slides and allowed to air dry. The mFISH procedure was carried out in accordance with the Metasystem manufacturer’s instructions for the use of the 24Xcyte human chromosome probe. Chromosome spreads were automatically captured using the Metafer imaging system and subsequent analyses were conducted with Isis software.

### COBRA-FISH Karyotyping of JVE059 cells

COBRA-FISH was performed as described in Szuhai and Tanke (2006). In brief, metaphase suspensions were dropped on microscopy glass slides and airdried overnight. Slides with metaphase chromosomes were pre-treated at 37 °C for 10 min with 100 μg/ml RNase A (Roche; Cat. No.: 10154105103) in 2x saline-sodium citrate (SSC; Sigma-Aldrich; Cat. No.: S0902) and were then incubated with 0.005% pepsin (Sigma-Aldrich; Cat. No.: P0525000) in 0.1 M HCl for 5 min at 37 °C. After a 10-min fixation at RT with 1% (v/v) formaldehyde (Merck; Cat. No.: 1.03999.1000) in PBS (pH 7.4), the specimens were dehydrated by three 3-min incubations in ethanol at increasing concentrations, 70%, 90% and 100%. Next, the coverslips were air-dried and exposed to whole-chromosome painting probes fluorescently labelled with the dyes 7-diethylaminocoumarin-3-carboxylic acid (DEAC), Cy3, Cy5 and rhodamine green by using the Universal Linkage System (ULS) kit (Kreatech Biotechnology, The Netherlands). After DNA denaturation at 80°C for 75 s, hybridizations were let to proceed for 3 days at 37°C in a humidified chamber. Unbound probes were eliminated by first washing with 0.1% Tween-20 (Promega; Cat. No.: PRH5152) in 2 × SSC, and then with 50% formamide (Merck; Cat. No.: 1.09684.1000) in 2 × SSC pH 7.0 at 44 °C followed by incubation at 60 °C in 0.1× SSC. Each washing step was done twice for 5 min. After dehydration by exposure to the above-mentioned increasing concentrations of ethanol, the specimens were air-dried and embedded in Citifluor AF1/DAPI (400 ng ml^−1^) solution (Aurion; Cat. No.: E17970). Stained chromosomes were visualized with a Leica DMRA fluorescence microscope (Leica, Wetzlar, Germany) and images were captured with the aid of a CoolSnap HQ2 camera (Photometrics, Tucson, USA). A minimum of 35 metaphase cells were analyzed from each sample and reported in the International System for Human Cytogenomic Nomenclature (ISCN) format.

### Antibodies

The following primary antibodies were used for immunofluorescence studies: guinea pig anti-CENP-C (1:1000; MBL International Corporation), chicken anti-GFP (1:2000; ab13970; Abcam), mouse anti–αTubulin clone B-512 (1:2000; Sigma-Aldrich), rabbit anti–Aurora B pT232 (1:1000; Rockland Immunochemicals), rabbit MPS1 pT676 (1:1000; a gift from Geert Kops, Hubrecht Institute, Utrecht, Netherlands; (Jelluma et al., 2008a), rabbit anti-ROD pT13/pS15 (1:1000; a gift from Prasad Jallepalli, Memorial Sloan Kettering Cancer Center) (Maciejowski et al., 2017), rabbit anti-Astrin (1:1000; a gift from Duane Compton, Geisel School of Medicine, Dartmouth) (Mack & Compton, 2001), mouse anti-Astrin clone C-1 (1:200; Sigma-Aldrich MABN2487), rabbit anti-Hec1 pS44 (1:1000; a gift from Jennifer DeLuca, Colorado State University) (DeLuca et al., 2011), rabbit anti-Spindly (1:5000; a gift from Arshad Desai), Sheep anti-CENP-F (1:800; a gift from Stephen Taylor), rabbit polyclonal anti-KNL1 pT943/pT1155 (1:1000; ab23) (Nijenhuis et al., 2014), rabbit anti-ZW10 (1:300; a gift from Andrea Musacchio), mouse anti-CENP-E clone1H12 (1:500; Abcam, Cat No.: ab5093), mouse anti-MAD1 (1:1000; Millipore MABE867). The following primary antibodies were used for Western blotting studies: mouse anti–α-tubulin DM1A (1:10,000; RRID:AB_477593; Sigma-Aldrich) and mouse anti-MPS1 (1:500; clone 4-112-3; Merck Millipore).

### Statistical analysis

Statistical analysis was performed using GraphPad Prism V10.0 (GraphPad Software, Inc.). All results presented in this article were obtained by pooling data from at least two independent experiments, unless stated otherwise. Values were considered statistically different whenever P < 0.05. P values: ns, not significant; *< 0.05; **< 0.01; ***< 0.001; ****< 0.0001. The normality of the samples was determined with a D’Agostino & Pearson test. Statistical analysis for two sample comparison, with normal or non-normal distribution, was performed with a t-test or Mann-Whitney test, respectively. For multiple group comparison a parametric one-way analysis of variance (ANOVA) or a nonparametric Kruskal-Wallis was used for samples with normal or non-normal distribution, respectively. All pairwise multiple comparisons were subsequently analyzed using either Student t-test (parametric) or Dunn’s (nonparametric) tests.

Kaplan-Meier plots were generated using ggsurvplot from R package survminer (v.0.4.9), and differences between groups were assessed using the log-rank test. Hazard ratios (HRs) and the corresponding 95% confidence intervals (95% CI) and P values were estimated using a Cox proportional hazards model implemented in the R package survival (v2.41–3), with CIN status as the sole predictor (univariable Cox model). Cox regression was performed only when each comparison group contained at least 10 patients. Censoring was handled by standard right-censoring at last follow-up. Detailed information on statistical significance for each condition can be found in the figures and their legends.

## DATA AVAILABILITY

All data supporting the findings of this study, including all raw data and complete statistical analysis files, are available as downloadable ZIP archives via the EMBO Journal submission system. The AC-ICAM datasets are deposited in the public repository FigShare: https://doi.org/10.6084/m9.figshare.16944775. The SNP array data of PCCCs are deposited in the public repository NCBI Gene Expression Omnibus (GEO) under accession numbers GSE67773 and GSE67774.

## AUTHOR CONTRIBUTION

**Ana Pinto-Teixeira:** Data curation; Formal analysis; Validation; Investigation; Visualization; Methodology; Writing - original draft. **André Oliveira**: Data curation; Formal analysis; Investigation; Visualization; Methodology. **Gonçalo Gouveia**: Formal analysis; Investigation; Visualization; Methodology. **Ana Sousa**: Formal analysis; Investigation; Visualization; Methodology. **Jessica Roelands**: Data curation; Formal analysis; Resources; Methodology; Software. **Marco Grillo**: Investigation; Visualization; Methodology. **Sergi Rodriguez-Calado**: Investigation; Visualization; Methodology. **Carlos Resende**: Investigation; Visualization; Methodology. **Helena Xavier-Ferreira**: Investigation; Visualization; Methodology. **Mariana Osswald**: Software. **Hugo Girão**: Investigation; Visualization; Methodology. **Patrícia Mesquita**: Investigation; Visualization; Methodology. **Raquel Almeida**: Resources; Supervision; Funding acquisition. **Marin Barisic**: Resources; Supervision; Funding acquisition; Writing-review and editing. **José Carlos Machado**: Resources; Supervision; Funding acquisition. **Claudio Sunkel**: Resources; Supervision; Funding acquisition. **Tom van Wezel**: Resources; Supervision; Funding acquisition; Methodology. **Daniele Fachinetti**: Resources; Supervision; Funding acquisition; Methodology; Writing-review and editing. **Karoly Szuhai**: Resources; Investigation; Formal analysis; Funding acquisition; Methodology; Writing-review and editing. **Noel de Miranda**: Data curation; Resources; Investigation; Formal analysis; Funding acquisition; Methodology; Writing-review and editing. **Carlos Conde:** Conceptualization; Project administration; Investigation; Data curation; Resources; Supervision; Funding acquisition; Visualization; Writing-original draft; Writing-review and editing.

## ACKNOWLEDGEMENTS

We thank Rene Medema (Oncode Institute, The Netherlands Cancer Institute, The Netherlands) for the CP5, Geert Kops (Hubrecht Institute, the Netherlands) for the phospho-specific MPS1^pT676^ antibody, Prasad Jallepalli (Memorial Sloan Kettering Cancer Center, USA) for the phospho-specific ROD^pT13/pS15^ antibody, Jennifer DeLuca (Colorado State University, USA) for the phospho-specific HEC1^pS44^ antibody, Arshad Desai (University of California, San Diego, USA) for the Spindly antibody, Stephen Taylor (University of Manchester, UK) for the CENP-F antibody, Adrian Saurin (University of Dundee, UK) for the phospho-specific KNL1^pT943/pT1155^ antibody, Duane Compton (Geisel School of Medicine, Dartmouth, USA) for the Astrin antibody, and Andrea Musacchio (Max Planck Institute of Molecular Physiology, Germany) for the ZW10 antibody. The authors acknowledge the i3S Scientific Platform ALM, a member of the national infrastructure Portuguese Platform of Bioimaging, for excellent support. Work in C.C. and C.S. lab was funded by National Funds through FCT-Fundação para a Ciência e a Tecnologia, I.P., under the projects UIDB/04293/2020 and UID/4293/2025 and by the project “Cancer Research on Therapy Resistance: From Basic Mechanisms to Novel Targets” - NORTE-01-0145-FEDER-000051, supported by Norte Portugal Regional Operational Programme (NORTE 2020), under the PORTUGAL 2020 Partnership Agreement, through the European Regional Development Fund (ERDF) and the project “How to build a mitotic checkpoint that cells can rely on?” - 2023.13952.PEX, supported by Fundação para a Ciência e a Tecnologia. C.C. was supported by a Scientific Employment Stimulus contract (2020.00067.CEECIND) and A.P-T was supported by a PhD fellowship (SFRH/BD/136334/2018) from Fundação para a Ciência e a Tecnologia. Work in M.B. lab was funded by the Lundbeck Foundation (R434-2023-431) and the Danish Cancer Society (BK25-01526).

## EXPANDED VIEW FIGURE LEGENDS

**Figure EV1.**
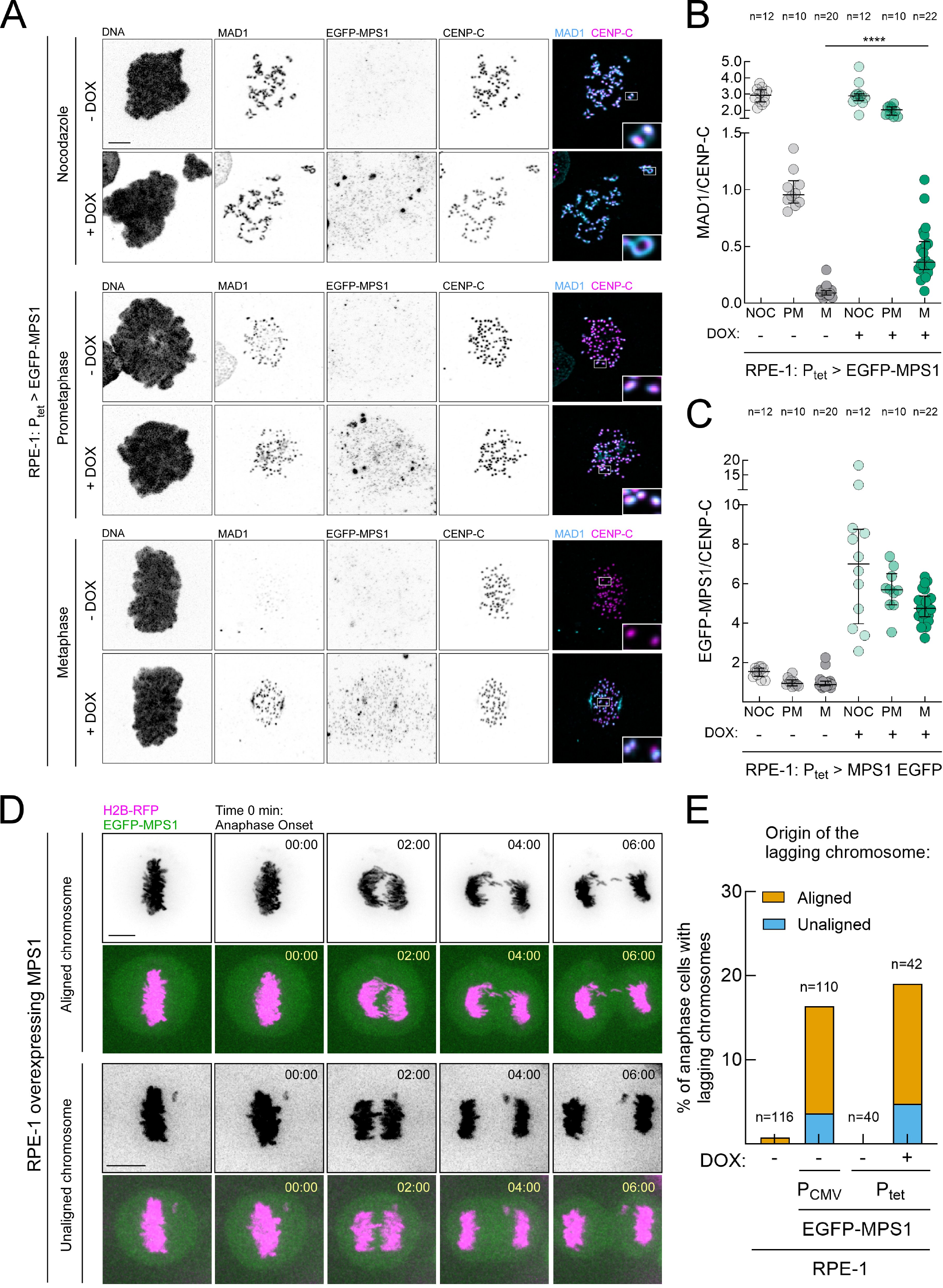
MPS1 overexpression maintains the SAC active at metaphase kinetochores and produces lagging chromatids originating from metaphase-aligned chromosomes. Related to Fig. 4. (A-C) Representative immunofluorescence images **(A)** and corresponding quantifications of MAD1 levels **(B)** and EGFP-MPS1 levels **(C)** at kinetochores of RPE-1 cells in control (-DOX) and MPS1 overexpression (+DOX) conditions, in nocodazole (NOC), prometaphase (PM) and metaphase (M). Insets depict magnifications of selected kinetochores. MAD1 and EGFP-MPS1 fluorescence intensities were determined relative to the CENP-C signal. All values in (B,C) were normalized to the mean value determined for prometaphase control RPE-1 cells, which was set to 1. Each data point represents the mean of an individual cell. **(D)** Selected still frames from representative live-cell imaging of mitotic progression of RPE-1 cells stably expressing H2B–RFP and EGFP–MPS1. Time (min) is shown relative to anaphase onset. **(E)** Frequency of anaphase events with lagging chromosomes that derived either from chromosomes that were aligned at the metaphase plate (aligned, orange bars) or from chromosomes that never aligned at the metaphase plate (unaligned, blue bars) in parental RPE-1 cells, RPE-1 cells constitutively overexpressing MPS1 (P_CMV_) and RPE-1 cells carrying a doxycycline-inducible MPS1 expression system (P_tet_). n denotes the number of anaphase cells analysed (filmed) for each condition. Data information: data in **(B)** and **(C)** are presented as median with interquartile range and data in **(E)** are presented as mean; asterisks indicate that differences between mean ranks are statistically significant; ∗∗∗∗p < 0.0001; Mann-Whitney test. Scale bars: 5μm.

**Figure EV2.**
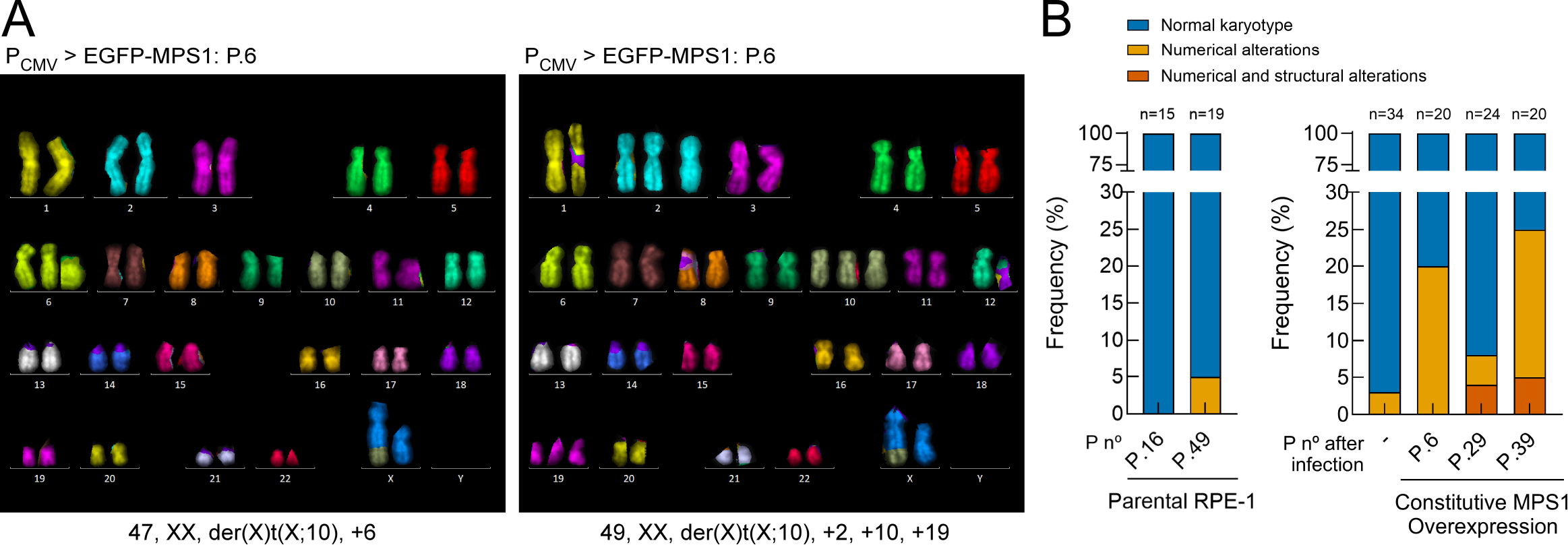
Constitutive MPS1 overexpression promotes karyotypic diversification in RPE-1 cells. Related to Fig. 4. (A) Representative multicolour fluorescence in situ hybridization (mFISH) karyograms of RPE-1 cells constitutively expressing CMV-driven EGFP–MPS1, analysed six passages after viral infection (P.6). The examples show a cell with a gain of chromosome 6 (left) and a cell with gains of chromosomes 2, 10 and 19 (right). Corresponding karyotypes are indicated below each karyogram. **(B)** Frequencies of cells retaining the parental control RPE-1 karyotype (blue), harbouring numerical chromosome alterations only (yellow), or harbouring both numerical and structural chromosome alterations (orange). Parental RPE-1 cells were analysed at passages 16 and 49. The baseline population before infection (–) and EGFP–MPS1-expressing cells at passages 6, 29 and 39 after infection were analysed as indicated. Bars represent the percentage of mitotic cells in each category; n denotes the number of mitotic spreads analysed. The constitutive der(X)t(X;10) rearrangement present in parental RPE-1 cells is considered the normal control karyotype and was not scored as an acquired structural alteration.

**Figure EV3.**
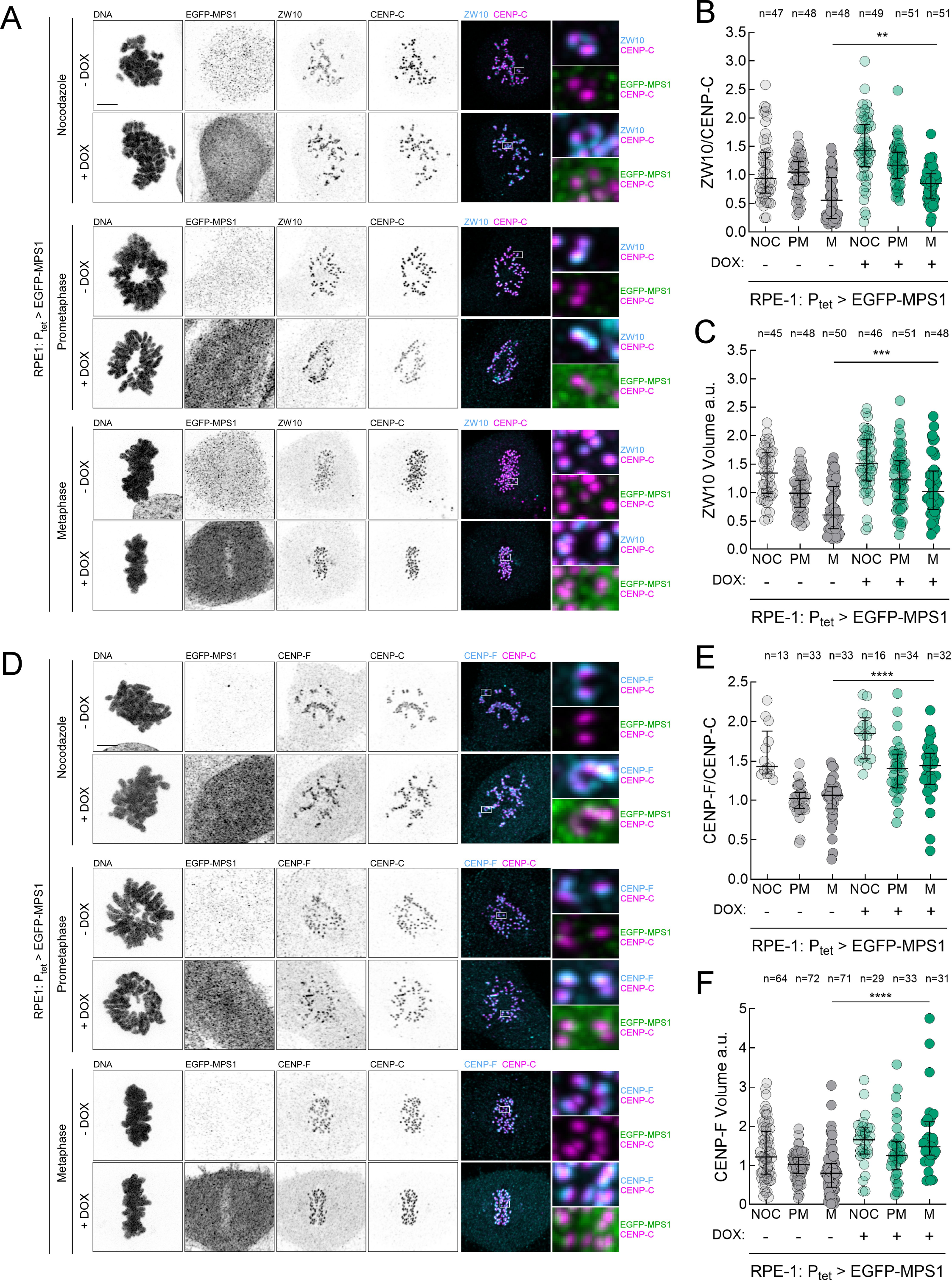
MPS1 overexpression increases the levels of ZW10 and CENP-F at metaphase kinetochores. Related to Fig. 6. (A-C) Representative immunofluorescence images **(A)** and corresponding quantifications of ZW10 levels **(B)** and ZW10 volumes **(C)** at kinetochores of RPE-1 cells in control (-DOX) and MPS1 overexpression (+DOX) conditions, in nocodazole (NOC), prometaphase (PM) and metaphase (M). Insets depict magnifications of selected kinetochores. ZW10 fluorescence intensities were determined relative to the CENP-C signal. All values in **(B)** and **(C)** were normalized to the mean value determined for prometaphase control RPE-1 cells, which was set to 1. Each data point represents the mean of an individual cell; n=2 independent experiments **(D-F)** Representative immunofluorescence images **(D)** and corresponding quantifications of CENP-F levels **(E)** and CENP-F volumes **(F)** at kinetochores of RPE-1 cells in control (-DOX) and MPS1 overexpression (+DOX) conditions, in nocodazole (NOC), prometaphase (PM) and metaphase (M). Insets depict magnifications of selected kinetochores. CENP-F fluorescence intensities were determined relative to the CENP-C signal. All values in **(E)** and **(F)** were normalized to the mean value determined for prometaphase control RPE-1 cells, which was set to 1. Each data point represents the mean of an individual cell; n≥3 independent experiments Data information: data in **(B)**, **(C)**, **(E)** and **(F)** are presented as medians with interquartile range; asterisks indicate that differences between mean ranks are statistically significant; \*\**p* < 0.01; \*\*\**p* < 0.001; \*\*\*\**p* < 0.0001; Mann-Whitney test. Scale bars: 5μm.

**Figure EV4.**
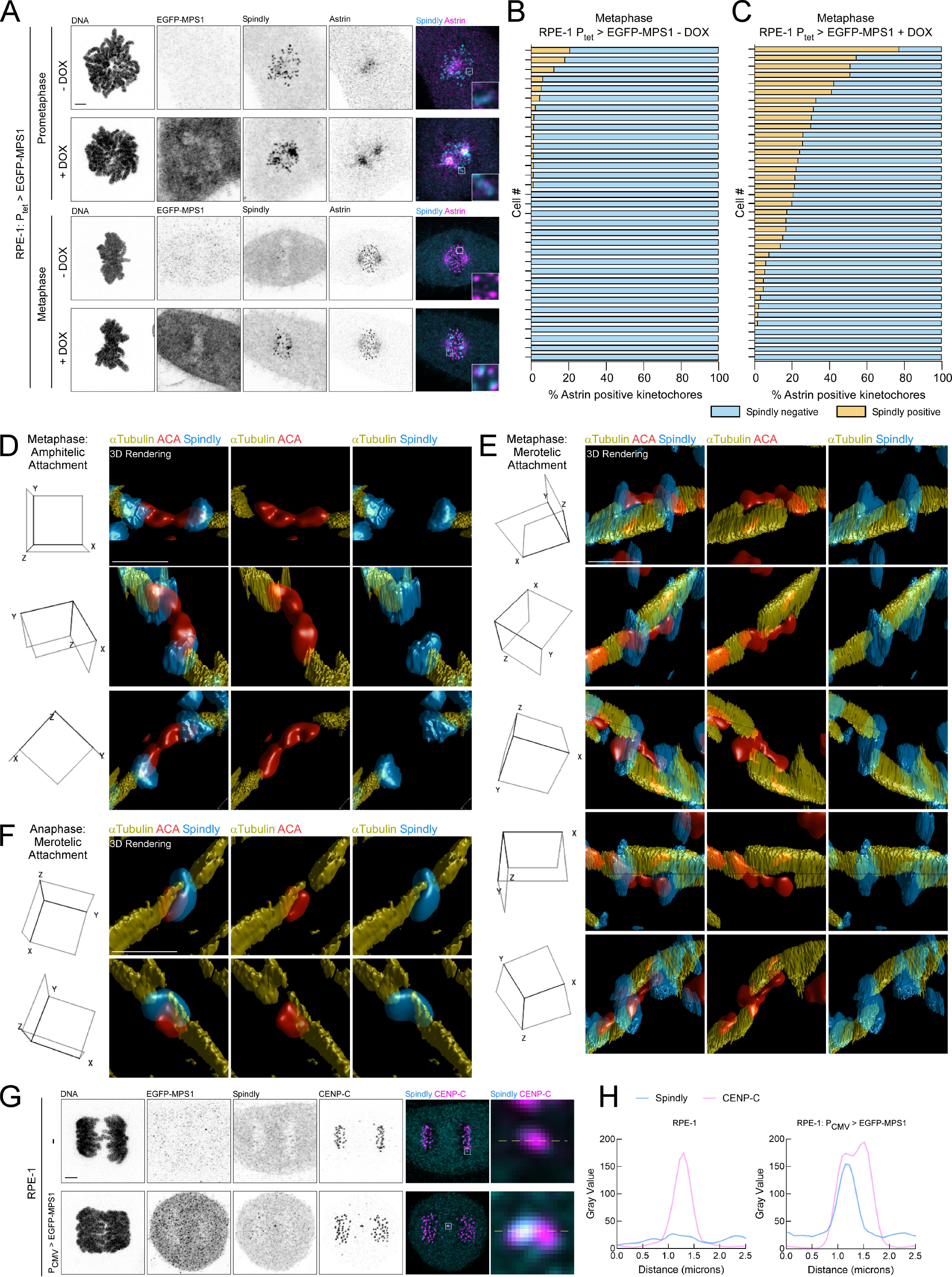
MPS1 overexpression sustains fibrous corona assembly at mature end-on attachments, promoting merotelic kinetochore–microtubule engagement. Related to Fig. 7. (A) Representative immunofluorescence images of Spindly and Astrin in prometaphase and metaphase of RPE-1 cells in control (-DOX) and MPS1 overexpression conditions (+DOX). Insets depict magnifications of selected kinetochores. **(B, C)** Histograms depicting the percentage of Astrin-positive metaphase kinetochores per cell that are also positive (orange bars) or negative (blue bars) for Spindly in control (-DOX) **(B)** and MPS1 overexpression (+DOX) **(C)** conditions as in **(A)**. Each bar represents one individual cell; n=2 independent experiments. **(D-F)** Three-dimensional reconstructions of the RPE-1 cells overexpressing MPS1 presented in **Fig. 7B**, **7D** and **7F** (selected Z-slices). Different perspectives of the same 3D reconstruction are shown, as indicated by the orientation cube displayed in the left of each panel. Microtubules (tubulin; yellow), Spindly (blue) and centromeres (ACA; red). Scale bars: 1 µm. **(G)** Representative immunofluorescence images of anaphases in control parental RPE-1 cells (-) and RPE-1 cells overexpressing MPS1 (P_CMV_). Spindly (cyan), CENP-C (magenta). Insets display magnifications of the outlined regions. **(H)** Plotted profiles display the variation in CENP-C and Spindly signals as a function of distance (µm) measured along the dashed yellow line indicated in the inset in **(G)**.

**Figure EV5.**
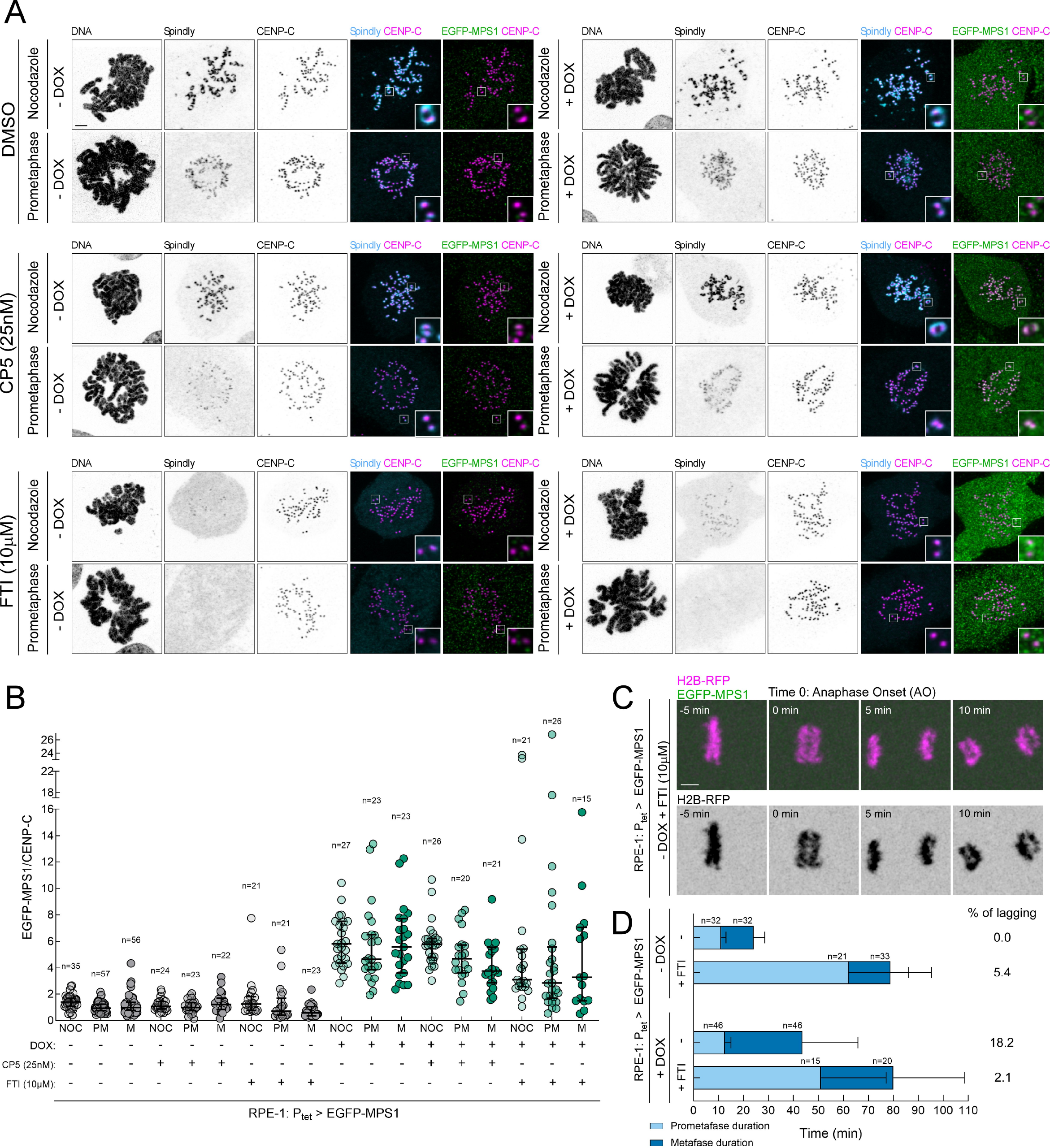
Partial inhibition of MPS1 and inhibition of Spindly farnesylation hinder Spindly kinetochore accumulation in RPE-1 cells overexpressing MPS1. Related to Fig. 8. **(A)** Representative immunofluorescence images of Spindly kinetochore localization in prometaphase and nocodazole-treated RPE-1 cells under control (-DOX) and MPS1 overexpression (+DOX) conditions. MPS1 was partially inhibited when indicated with CP5. Spindly farnesylation was prevented when indicated with FTI-277 (FTI). Insets depict magnifications of selected kinetochores. The corresponding quantification of Spindly levels and volumes at kinetochores are presented in Fig. 8B and 8C, respectively. **(B)** Quantification of EGFP-MPS1 levels at kinetochores of RPE-1 cells in control (-DOX) and MPS1 overexpression (+DOX) conditions, in nocodazole (NOC), prometaphase (PM) and metaphase (M). MPS1 was partially inhibited when indicated with CP5. Spindly farnesylation was prevented when indicated with FTI-277 (FTI). Representative immunofluorescence images of this quantification are depicted in (A) and in Fig. 8A. EGFP-MPS1 fluorescence intensities were determined relative to the CENP-C signal. Each data point represents the mean of an individual cell; n≥2 independent experiments. **(C)** Selected still frames from representative live-cell imaging of chromosome segregation in RPE-1 cells (-DOX) stably expressing H2B–RFP and treated with FTI-277 (FTI). Time (min) is shown relative to anaphase onset (AO). **(D)** Mitotic timing of RPE-1 cells under control (-DOX) and MPS1 overexpression (+DOX) conditions. Cells were treated when indicated with FTI-277 (+FTI). Light-coloured bars represent the prometaphase duration (length of time measured between NEB and the first frame with all chromosomes aligned at the metaphase plate); and dark-coloured bars represent the metaphase duration (length of time measured between the first frame after chromosome alignment and anaphase onset). The percentage of cells with lagging chromosomes is indicated for each condition. n indicates the number of cells filmed for each condition. Data information: data in **(B)** are presented as medians with interquartile range and data in **(D)** are presented as mean with standard deviation. Scale bar: 5μm.

