## Appendix for "Elevated MPS1 converts the fibrous corona into a source of chromosomal instability in colon cancer"

### Table of contents:

**Appendix Figures S1-S12,**

**Appendix Figures S1-S12 legends**

**Movies EV1-EV35 legends**

**Appendix references**

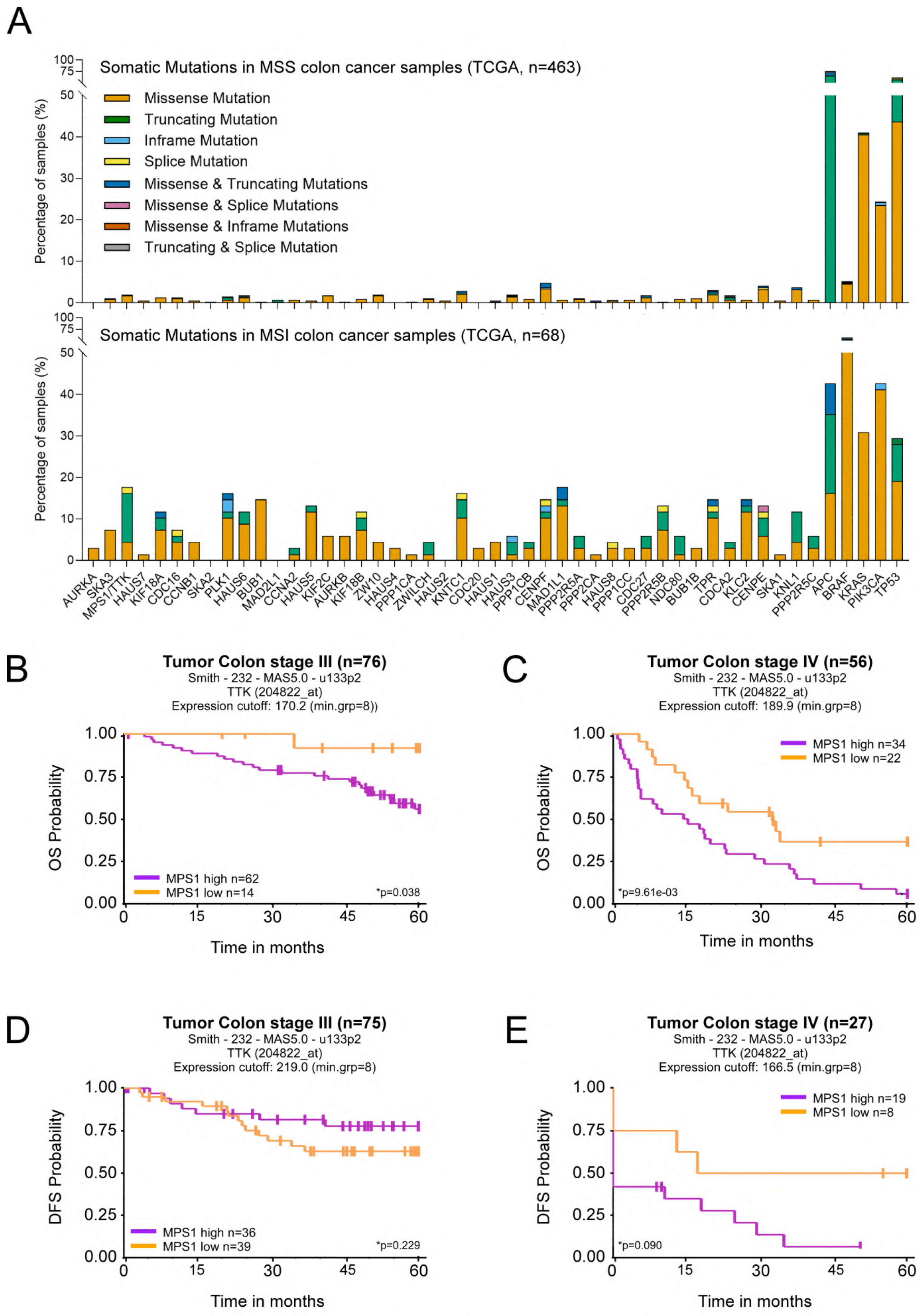


Appendix Figure S1 Pinto-Teixeira et al

**Appendix Figure S1. MPS1 overexpression correlates with a worse prognosis. Related to Figure 1.**

**(A)** Frequency of somatic mutations in 44 mitotic genes across 531 tumours of the TCGA COAD-READ cohort stratified by microsatellite instability status (MSI vs MSS).

**(B-E)** Kaplan–Meier plots of overall survival (OS), **(B)** and **(C)** and disease-free survival (DFS), **(D)** and **(E)** for patients with stage III and stage IV colon cancer from the GSE17538 cohort (Smith et al., 2010), stratified by MPS1 expression. Patients were dichotomised into MPS1-high and MPS1-low groups using the expression cut-offs indicated. Vertical lines denote censored observations. Depicted **P* values compare overall differences in survival between expression-defined groups and were calculated using a two-sided log-rank test.

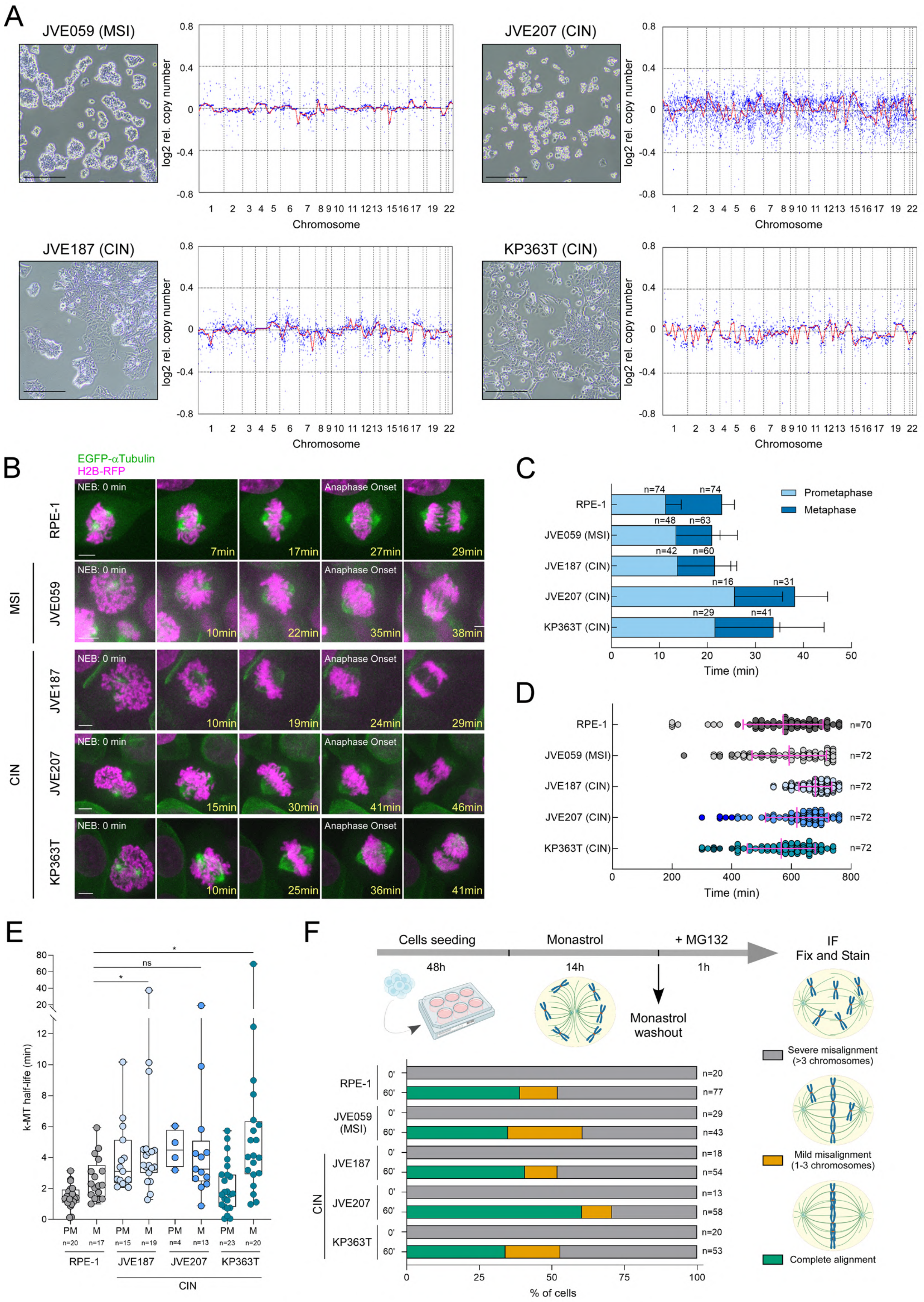


Appendix Figure S2 Pinto-Teixeira et al

**Appendix Figure S2. Characterization of patient-derived colon cancer cells. Related to Figure 2.**

**(A)** Representative brightfield images of patient-derived colon cancer cells (PCCCs) and corresponding genome-wide copy-number alteration (CNA) profiles determined by array comparative genomic hybridization and segmented using circular binary segmentation. Scale bar: 200μm.

**(B)** Selected still frames from representative live-cell imaging movies of mitotic progression of RPE-1, JVE059 (MSI), and the $CIN^+$ PCCCs JVE187, JVE207, and KP363T stably expressing H2B–RFP and EGFP–αTubulin. Time (min) is shown relative to nuclear envelope breakdown (NEB). Scale bar: 5 μm.

**(C)** Mitotic timing of cells indicated in **(B)**. Light-coloured bars represent the prometaphase duration (length of time measured between NEB and the first frame with all chromosomes aligned at the metaphase plate); and dark-coloured bars represent the metaphase duration (length of time measured between the first frame after chromosome alignment and anaphase onset). n indicates the number of cells filmed for each cell line.

**(D)** Mitotic duration of RPE-1, JVE059 (MSI), and the $CIN^+$ PCCCs in the presence of nocodazole. Mitotic progression was monitored by time-lapse microscopy with frames acquired at 20-minute intervals. The mitotic duration was defined by the length of time measured between nuclear envelope breakdown and chromatin decondensation. $n \geq 70$ cells for each condition.

**(E)** Quantification of kinetochore-microtubule half-life of RPE-1 and the $CIN^+$ PCCCs JVE187, JVE207, and KP363T in prometaphase (PM) and metaphase (M). Each data point represents an individual cell. The boxes represent median and interquartile interval; the bars represent minimum and maximum values.

**(F)** Schematic representation of the experimental pipeline for the monastrol washout experiment. The graph represents chromosome congression after monastrol washout for RPE-1, JVE059 (MSI), and the $CIN^+$ PCCCs JVE187, JVE207, and KP363T. Green indicates complete alignment, defined as full congression of all chromosomes to the metaphase plate; orange indicates mild misalignment, defined as the presence of 1–3 chromosomes that failed to congress; and grey indicates severe misalignment, defined as more than 3 chromosomes that failed to congress to the metaphase plate ($n \geq 2$ independent experiments).

Data information: data in **(C)** and **(D)** are presented as mean with standard deviations, and data in (F) are plotted as fractions of the total; asterisks indicate that differences between mean ranks are statistically significant; $^*p < 0.05$; ns, non-significant; Kruskal-Wallis, Dunn's multiple comparison test.

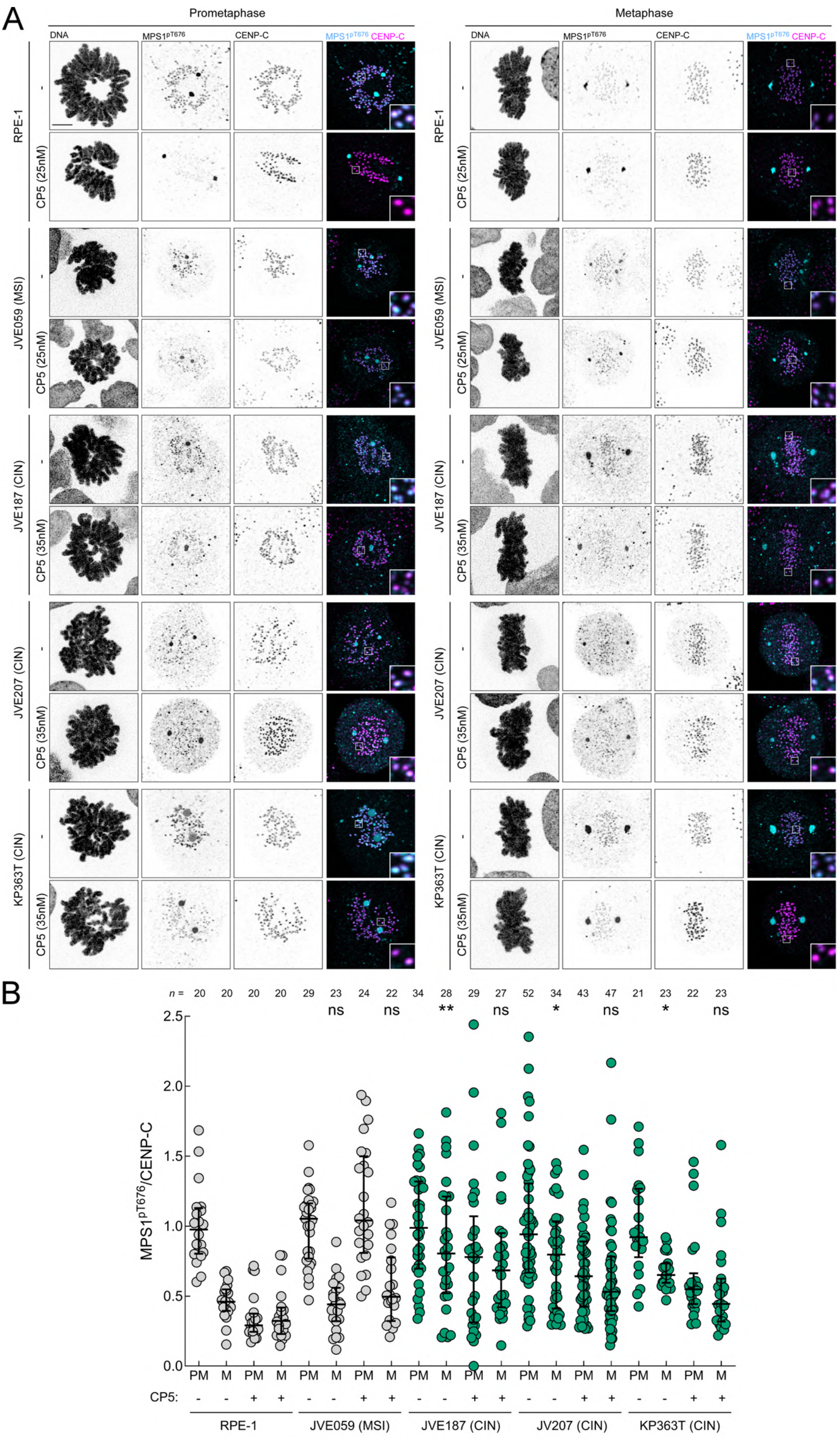


Appendix Figure S3 Pinto-Teixeira et al

**Appendix Figure S3. CIN$^+$ patient-derived colon cancer cells exhibit increased levels of MPS1 activity at kinetochores in metaphase. Related to Figure 3.**

**(A,B)** Representative immunofluorescence images **(A)** and corresponding quantifications **(B)** of MPS1 T676 phosphorylation (MPS1$^{pT676}$) levels at kinetochores in prometaphase (PM) and metaphase (M) of RPE-1 cells and PCCCs JVE059 (MSI), JVE187 (CIN$^+$), JVE207 (CIN$^+$) and KP363T (CIN$^+$). MPS1$^{pT676}$ intensities were determined relative to CENP-C signal. MPS1 was partially inhibited (when indicated) with CP5. Insets depict magnifications of selected kinetochores. All values in **(B)** were normalized to the mean value determined for prometaphase cells within each respective cell line, which was set to 1. Each data point represents the mean of an individual cell. n=3 independent experiments.

Data information: data in **(B)** are presented as median with interquartile range; asterisks indicate that differences between mean ranks are statistically significant relative to RPE-1 metaphase control; $^{*}p < 0.05$; $^{**}p < 0.01$; ns, non-significant; Kruskal-Wallis, Dunn's multiple comparison test. Scale bar: 5μm.

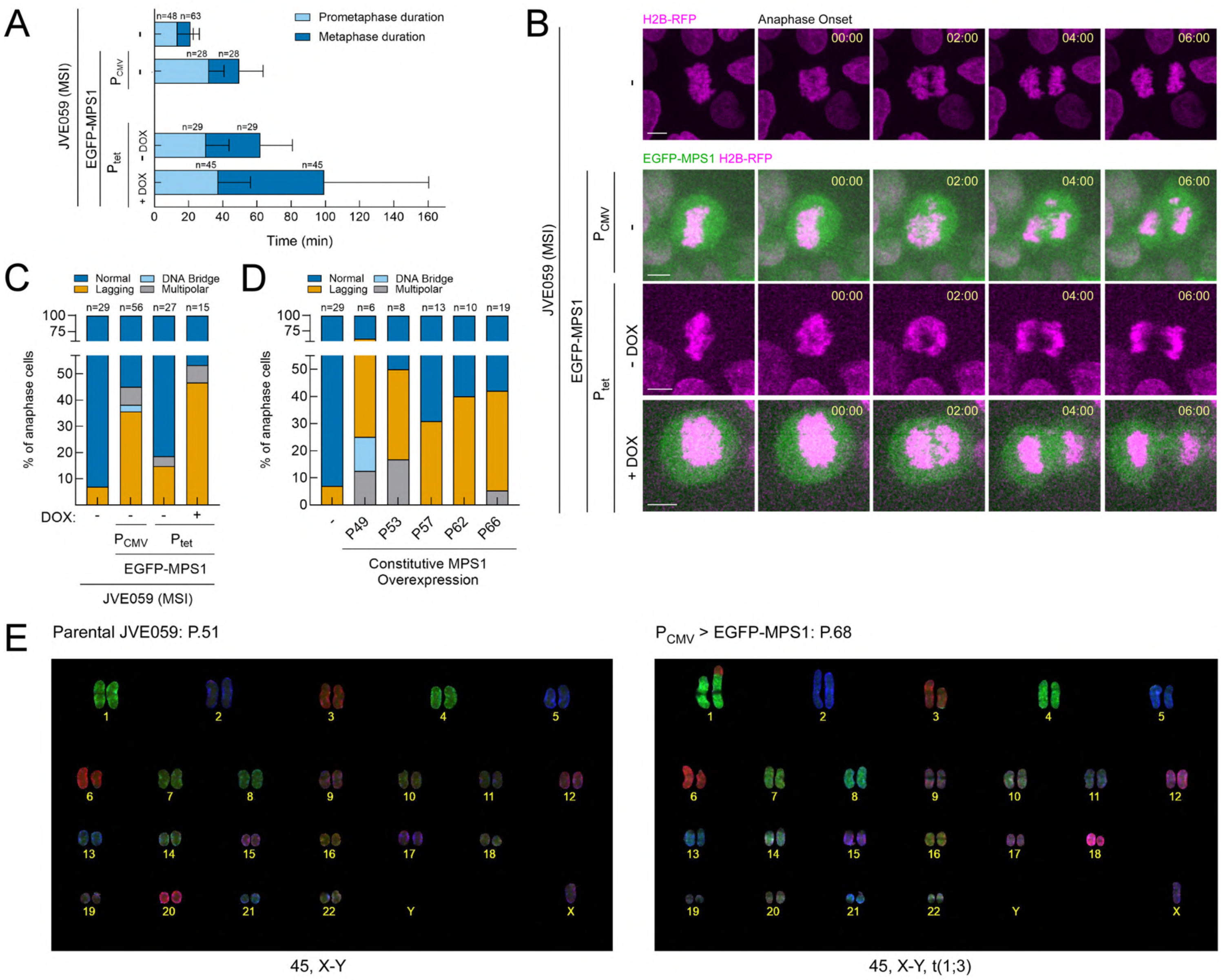


Appendix Figure S4 Pinto-Teixeira et al

**Appendix Figure S4. MPS1 overexpression increases the rate of chromosome mis-segregation in colon cancer MSI JVE059 cells. Related to Figure 4.**

**(A)** Mitotic timing of JVE059 control cells, JVE059 cells with constitutive MPS1 overexpression ($P_{CMV}$), and JVE059 cells carrying a doxycycline-inducible (DOX) MPS1 expression system ($P_{tet}$). Light-coloured bars represent the prometaphase duration (length of time measured between NEB and the first frame with all chromosomes aligned at the metaphase plate); and dark-coloured bars represent the metaphase duration (length of time measured between the first frame after chromosome alignment and anaphase onset). n indicates the number of cells filmed for each condition.

**(B)** Selected still frames from representative live-cell imaging of mitotic progression of JVE059 cells, JVE059 cells constitutively overexpressing MPS1 ($P_{CMV}$) and JVE059 cells carrying a doxycycline-inducible MPS1 expression system ($P_{tet}$). All cells stably express H2B-RFP to visualize the chromosomes (magenta). MPS1 overexpression is shown in green for both $P_{CMV}$ and $P_{tet}$ (+DOX) conditions. Time (min) is shown relative to anaphase onset.

**(C)** Quantification of chromosome segregation fidelity for the cell lines shown in **(B)**. Graph bars represent the frequency of indicated anaphase output. n denotes the number of anaphase events analysed (filmed) for each cell line.

**(D)** Quantification of chromosome segregation fidelity for JVE059 cells constitutively overexpressing MPS1 over several passages. Graph bars represent the frequency of indicated anaphase output. n denotes the number of anaphase events analysed (filmed) for each cell line and for each passage.

**(E)** Representative images of COBRA-FISH analysis of chromosome spreads of parental JVE059 cells (left) and JVE059 cells constitutively overexpressing MPS1 (right) at the indicated passage number. The corresponding karyotypes are indicated below the images.

Data information: data in **(A)** are presented as mean with standard deviations and data in **(C)** and **(D)** are plotted as fractions of the total. Scale bar: 5 µm.

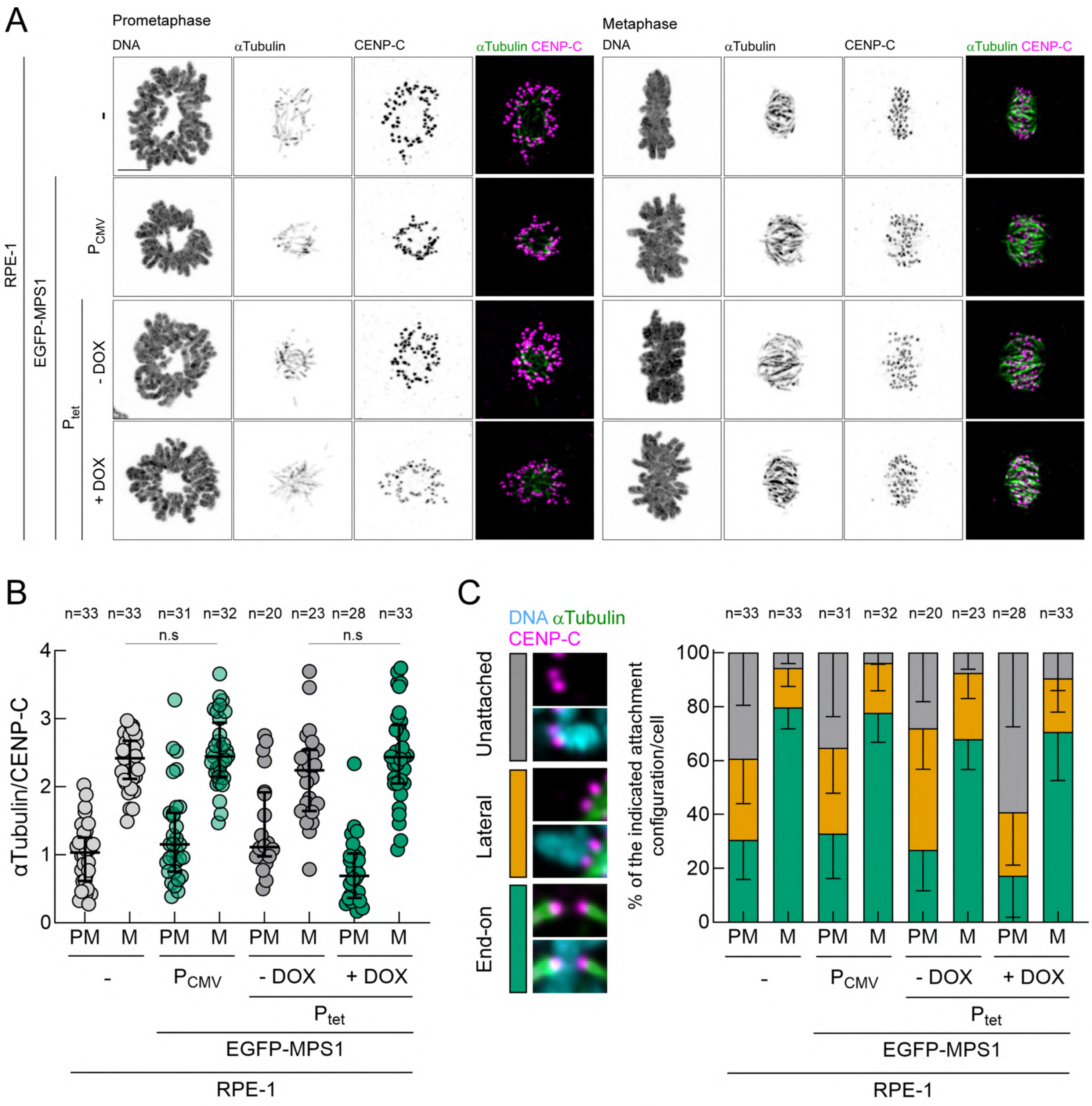


Appendix Figure S5 Pinto-Teixeira et al

**Appendix Figure S5. MPS1 overexpression does not decrease the stability of kinetochore-microtubule attachments in RPE-1 cells. Related to Fig. 5.**

**(A-B)** Representative immunofluorescence images **(A)** and corresponding quantifications of αTubulin levels **(B)** interacting with kinetochores in parental RPE-1 cells, RPE-1 cells constitutively overexpressing MPS1 ($P_{CMV}$) and RPE-1 cells carrying a doxycycline-inducible MPS1 expression system ($P_{tet}$), in prometaphase (PM) and metaphase (M). αTubulin fluorescence intensities were determined relative to the CENP-C signal. All values in **(B)** were normalized to the mean value determined for prometaphase control parental RPE-1 cells, which was set to 1. Each data point represents the mean of an individual cell; n=2 independent experiments.

**(C)** Percentage of the indicated attachment configurations per cell in the experimental conditions indicated in **(A)**. Green bars indicate end-on kinetochore-microtubule attachment, orange bars indicate lateral kinetochore-microtubule attachment and grey bars indicate unattached kinetochore. n denotes the number of prometaphase (PM) and metaphase (M) cells analysed for each cell line.

Data information: data in **(B)** are presented as median with interquartile range, data in **(C)** are presented as mean with standard deviations; ns, non-significant; Kruskal-Wallis, Dunn's multiple comparison test. Scale bars: 5μm.

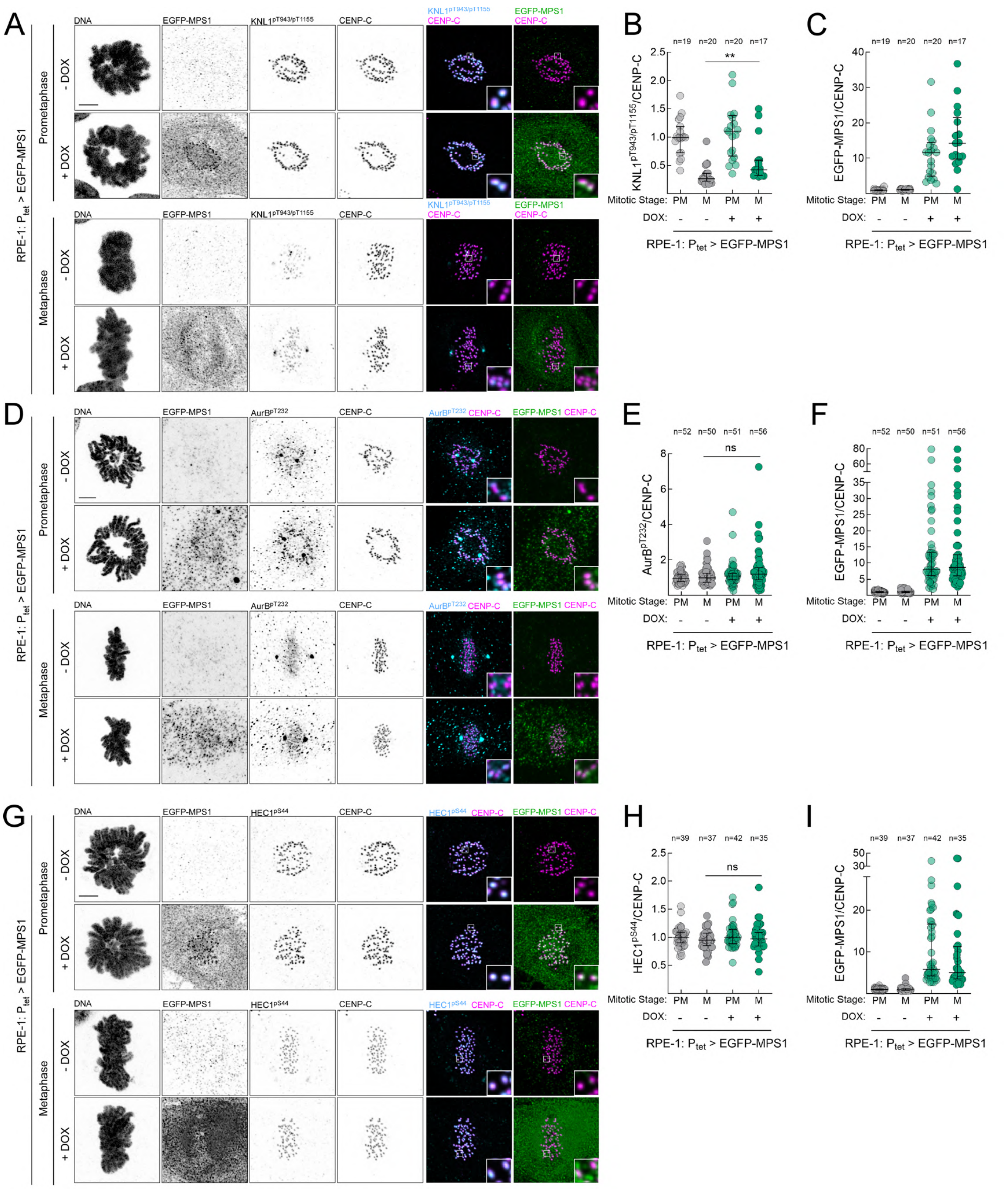

Appendix Figure S6 Pinto-Teixeira et al

**Appendix Figure S6. MPS1 overexpression promotes merotely independently of global changes in Aurora B signalling. Related to Figure 5.**

**(A-C)** Representative immunofluorescence images **(A)** and corresponding quantifications of KNL1 T943 and T1155 phosphorylation ($KNL1^{pT943/pT1155}$) levels **(B)** and EGFP-MPS1 levels **(C)** at kinetochores of RPE-1 cells in control (-

DOX) and MPS1 overexpression (+DOX) conditions, in prometaphase (PM) and metaphase (M) Insets depict magnifications of selected kinetochores. KNL1$^{pT943/pT1155}$ and EGFP-MPS1 fluorescence intensities were determined relative to the CENP-C signal. All values in **(B)** and **(C)** were normalized to the mean value determined for prometaphase control RPE-1 cells, which was set to 1. Each data point represents the mean of an individual cell; n=2 independent experiments.

**(D-F)** Representative immunofluorescence images **(D)** and corresponding quantifications of Aurora B T232 phosphorylation (AurB$^{pT232}$) levels **(E)** and EGFP-MPS1 levels **(F)** at kinetochores of RPE-1 cells in control (-DOX) and MPS1 overexpression (+DOX) conditions, in prometaphase (PM) and metaphase (M). Insets depict magnifications of selected kinetochores. AurB$^{pT232}$ and EGFP-MPS1 fluorescence intensities were determined relative to the CENP-C signal. All values in **(E)** and **(F)** were normalized to the mean value determined for prometaphase control RPE-1 cells, which was set to 1. Each data point represents the mean of an individual cell; n=5 independent experiments.

**(G-I)** Representative immunofluorescence images **(G)** and corresponding quantifications of HEC1 S44 phosphorylation (HEC1$^{pS44}$) levels **(H)** and EGFP-MPS1 levels **(I)** at kinetochores of RPE-1 cells in control (-DOX) and MPS1 overexpression (+DOX) conditions, in prometaphase (PM) and metaphase (M). Insets depict magnifications of selected kinetochores. HEC1$^{pS44}$ and EGFP-MPS1 fluorescence intensities were determined relative to the CENP-C signal. All values in **(H)** and **(I)** were normalized to the mean value determined for prometaphase control RPE-1 cells, which was set to 1. Each data point represents the mean of an individual cell; n=5 independent experiments.

Data information: data in **(B)**, **(C)**, **(E)**, **(F)**, **(H)** and **(I)** are presented as medians with interquartile range; asterisks indicate that differences between mean ranks are statistically significant; $^{**}p < 0.01$; ns, non-significant; Mann-Whitney test. Scale bars: 5μm.

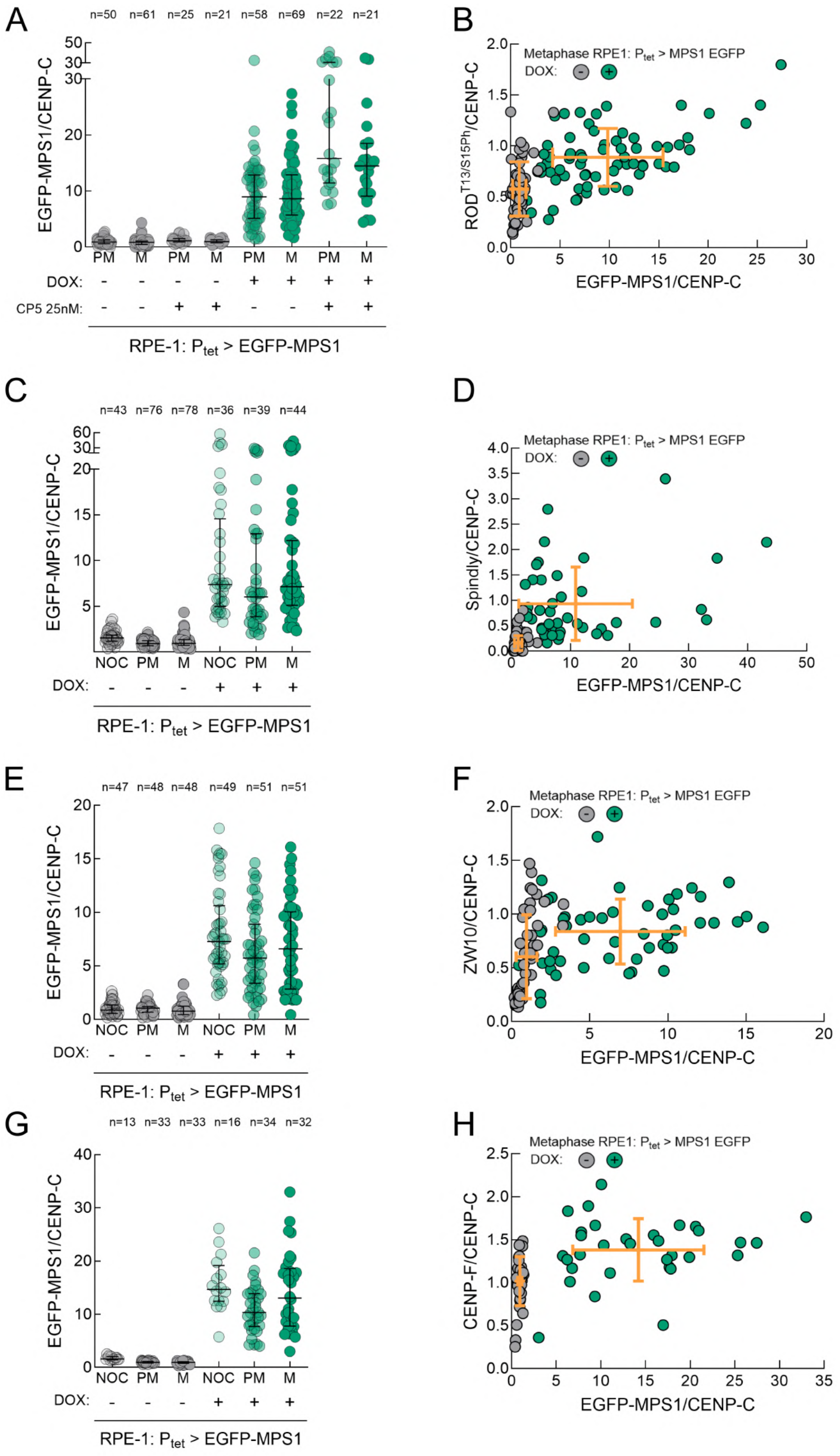


Appendix Figure S7 Pinto-Teixeira et al

**Appendix Figure S7. MPS1 overexpression increases the levels of ROD$^{pT13/pS15}$ and promotes abnormal accumulation of Spindly, ZW10 and CENP-F at metaphase kinetochores. Related to Figure 6 and Figure EV3.**

**(A)** Quantifications of EGFP-MPS1 levels at kinetochores of RPE-1 cells in control (-DOX) and MPS1 overexpression (+DOX) conditions, in prometaphase (PM) and metaphase (M), as presented in Figure 6A. MPS1 was partially inhibited when indicated with CP5. EGFP-MPS1 fluorescence intensities were determined relative to the CENP-C signal. Each data point represents the mean of an individual cell; n=4 independent experiments.

**(B)** ROD phosphorylation at T13 and S15 (ROD$^{pT13/pS15}$) at metaphase kinetochores, as presented in Figure 6B, plotted over EGFP-MPS1 kinetochore levels. ROD$^{pT13/pS15}$ and EGFP-MPS1 fluorescence intensities were determined relative to the CENP-C signal.

**(C)** Quantifications of EGFP-MPS1 levels at kinetochores of RPE-1 cells in control (-DOX) and MPS1 overexpression (+DOX) conditions, in nocodazole (NOC), prometaphase (PM) and metaphase (M), as presented in Figure 6C. EGFP-MPS1 fluorescence intensities were determined relative to the CENP-C signal. Each data point represents the mean of an individual cell; n=3 independent experiments.

**(D)** Spindly levels at metaphase kinetochores, as presented in Figure 6D, plotted over EGFP-MPS1 kinetochore levels. Spindly and EGFP-MPS1 fluorescence intensities were determined relative to the CENP-C signal.

**(E)** Quantifications of EGFP-MPS1 levels at kinetochores of RPE-1 cells in control (-DOX) and MPS1 overexpression (+DOX) conditions, in nocodazole (NOC), prometaphase (PM) and metaphase (M), as presented in Figure EV3A. EGFP-MPS1 fluorescence intensities were determined relative to the CENP-C signal. Each data point represents the mean of an individual cell; n=2 independent experiments.

**(F)** ZW10 levels at metaphase kinetochores, as presented in Figure EV3B, plotted over EGFP-MPS1 kinetochore levels. ZW10 and EGFP-MPS1 fluorescence intensities were determined relative to the CENP-C signal.

**(G)** Quantifications of EGFP-MPS1 levels at kinetochores of RPE-1 cells in control (-DOX) and MPS1 overexpression (+DOX) conditions, in nocodazole (NOC), prometaphase (PM) and metaphase (M), as presented in Figure EV3D. EGFP-MPS1 fluorescence intensities were determined relative to the CENP-C signal. Each data point represents the mean of an individual cell; n≥3 independent experiments.

**(H)** CENP-F levels at metaphase kinetochores, as presented in Figure EV3E, plotted over EGFP-MPS1 kinetochore levels. CENP-F and EGFP-MPS1 fluorescence intensities were determined relative to the CENP-C signal.

Data information: data in **(A)**, **(C)**, **(E)** and **(G)** are presented as medians with interquartile range, and data in **(B)**, **(D)**, **(F)** and **(H)** are presented as mean with standard deviation (orange bars).

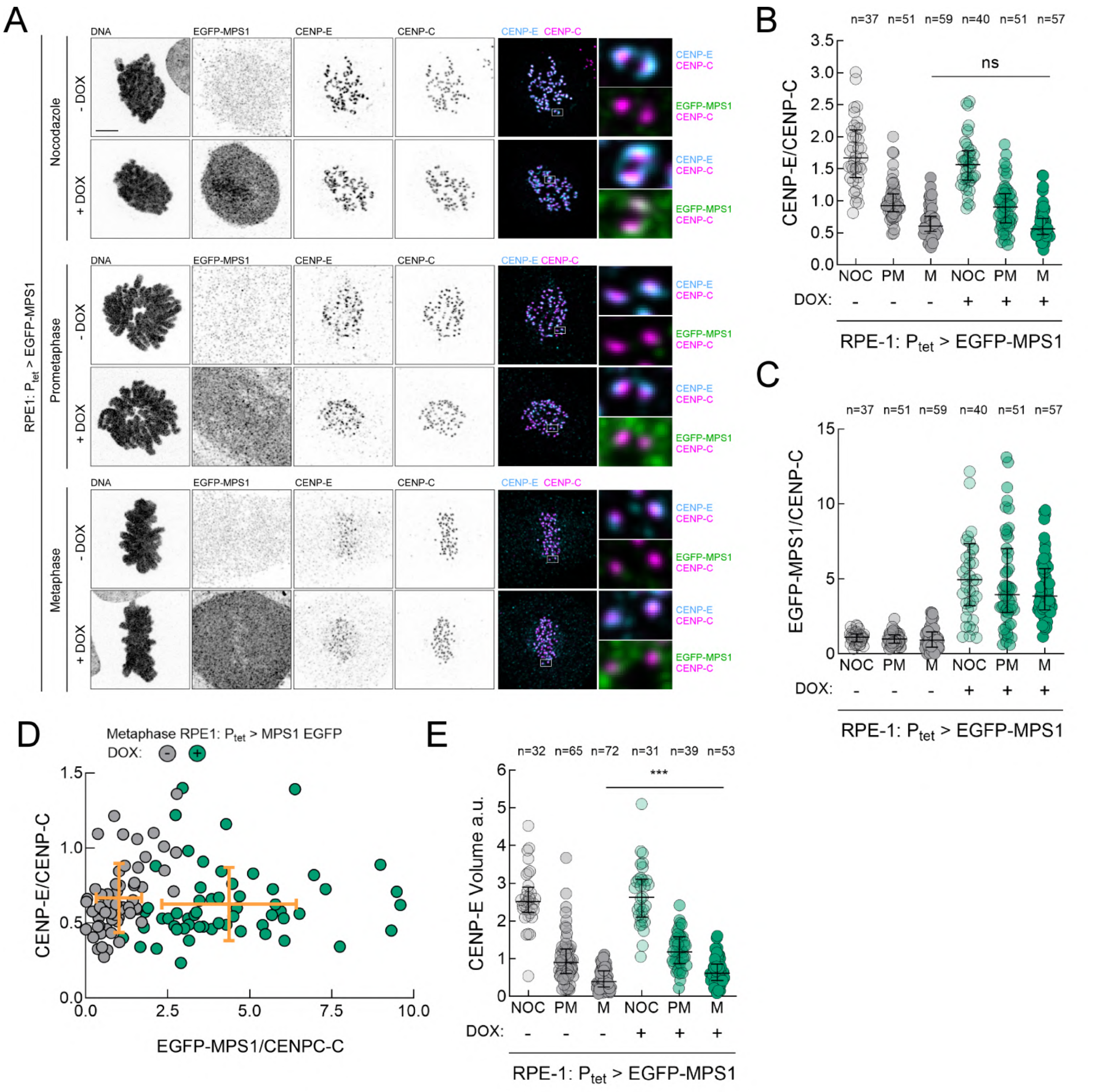


Appendix Figure S8 Pinto-Teixeira et al

**Appendix Figure S8. MPS1 overexpression increases the volume of CENP-E at metaphase kinetochores. Related to Figure 6 and Figure EV3.**

**(A-C)** Representative immunofluorescence images **(A)** and corresponding quantifications of CENP-E **(B)** and EGFP-MPS1 levels **(C)** at kinetochores of RPE-1 cells in control (-DOX) and MPS1 overexpression (+DOX) conditions, in nocodazole (NOC), prometaphase (PM) and metaphase (M). Insets depict magnifications of selected kinetochores. CENP-E and EGFP-MPS1 fluorescence intensities in **(B)** and **(C)** were determined relative to the CENP-C signal. **(D)** CENP-E levels at metaphase kinetochores, as presented in **(A)** and quantified in **(B)**, plotted over EGFP-MPS1 kinetochore levels as presented in **(A)** and quantified in **(C)** . All values in **(B)** and **(C)** were normalized to the mean value determined for prometaphase control RPE-1 cells, which was set to 1. Each data point represents the mean of an individual cell. n=3 independent experiments.

**(E)** CENP-E volumes at kinetochores of RPE-1 cells in control (-DOX) and MPS1 overexpression (+DOX) conditions, in nocodazole (NOC), prometaphase (PM) and metaphase (M). All values were normalized to the mean value determined for prometaphase control RPE-1 cells, which was set to 1. Each data point represents the mean of an individual cell. n=3 independent experiments.

Data information: data in **(B)**, **(C)** and **(E)** are presented as medians with interquartile range and data in **(D)** are presented as mean with standard deviation (orange bars); asterisks indicate that differences between mean ranks are statistically significant; $^{***}p < 0.001$; ns, non-significant; Mann-Whitney test. Scale bar: 5μm.

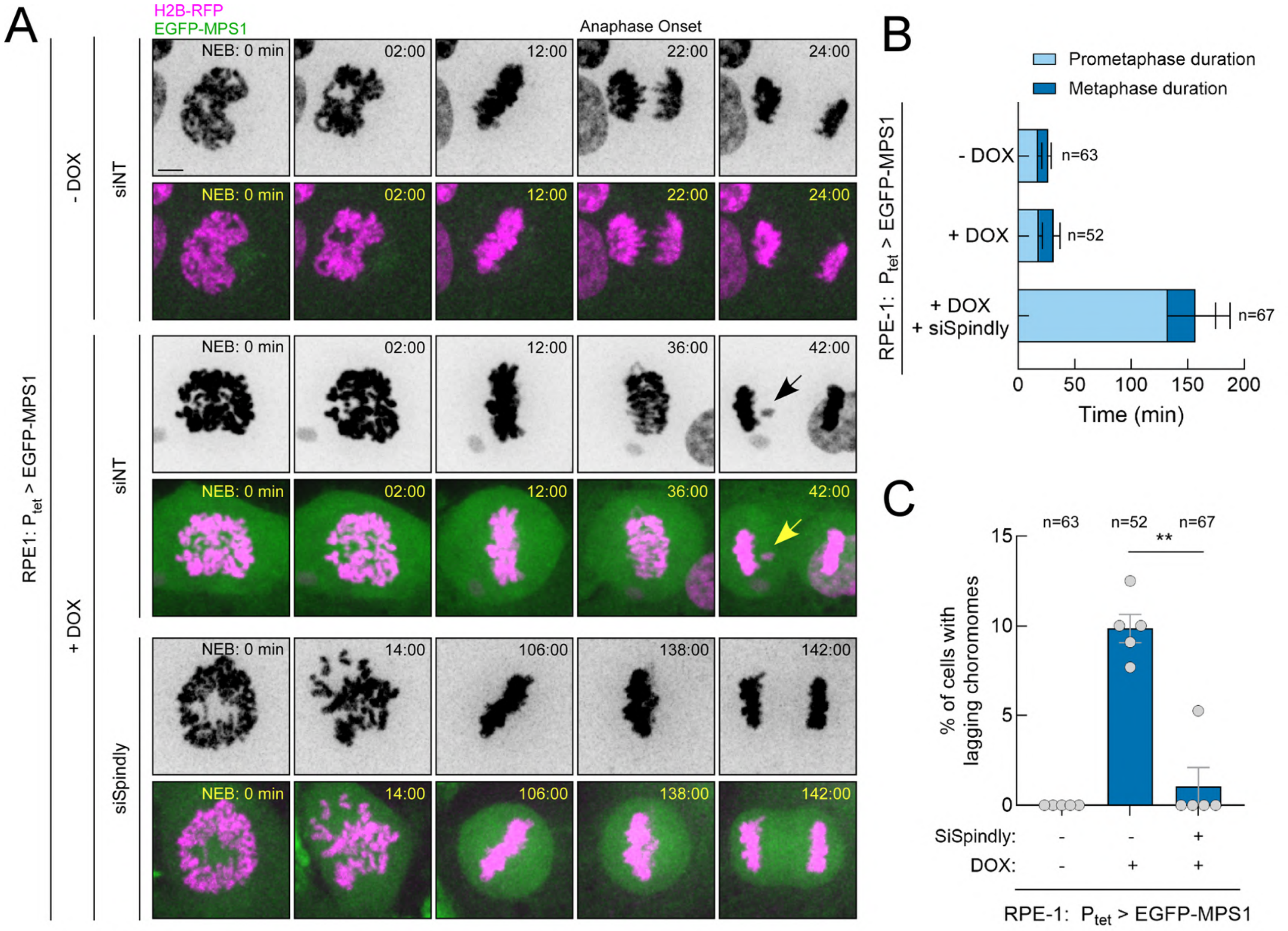


Appendix Figure S9 Pinto-Teixeira et al

**Appendix Figure S9. Spindly depletion restores anaphase fidelity in RPE-1 cells overexpressing MPS1. Related to Figure 8**

**(A)** Selected still frames from representative live-cell imaging of mitotic progression of RPE-1 cells stably expressing H2B-RFP (magenta) in control (-DOX) and MPS1 overexpression (+DOX) conditions. EGFP-MPS1 is depicted in green. Time (min) is indicated relative to nuclear envelope breakdown (NEB).

**(B)** Mitotic timing of cells indicated in **(A)**. Light-coloured bars represent the prometaphase duration (length of time measured between NEB and the first frame with all chromosomes aligned at the metaphase plate); and dark-coloured bars represent the metaphase duration (length of time measured between the first frame after chromosome alignment and anaphase onset). n indicates the number of cells filmed for each cell line; n=5 independent experiments.

**(C)** Quantification of the percentage of anaphase cells exhibiting lagging chromosomes as in **(A)**. each dot represents an independent experiment (n=5). n indicates the number of cells filmed for each condition.

Data information: data in **(B)** are presented as mean with standard deviation and data in **(C)** are presented as mean with standard error of the mean; asterisks indicate that differences between mean ranks are statistically significant; $^{**}p < 0.01$; Mann-Whitney test. Scale bar: 5µm.

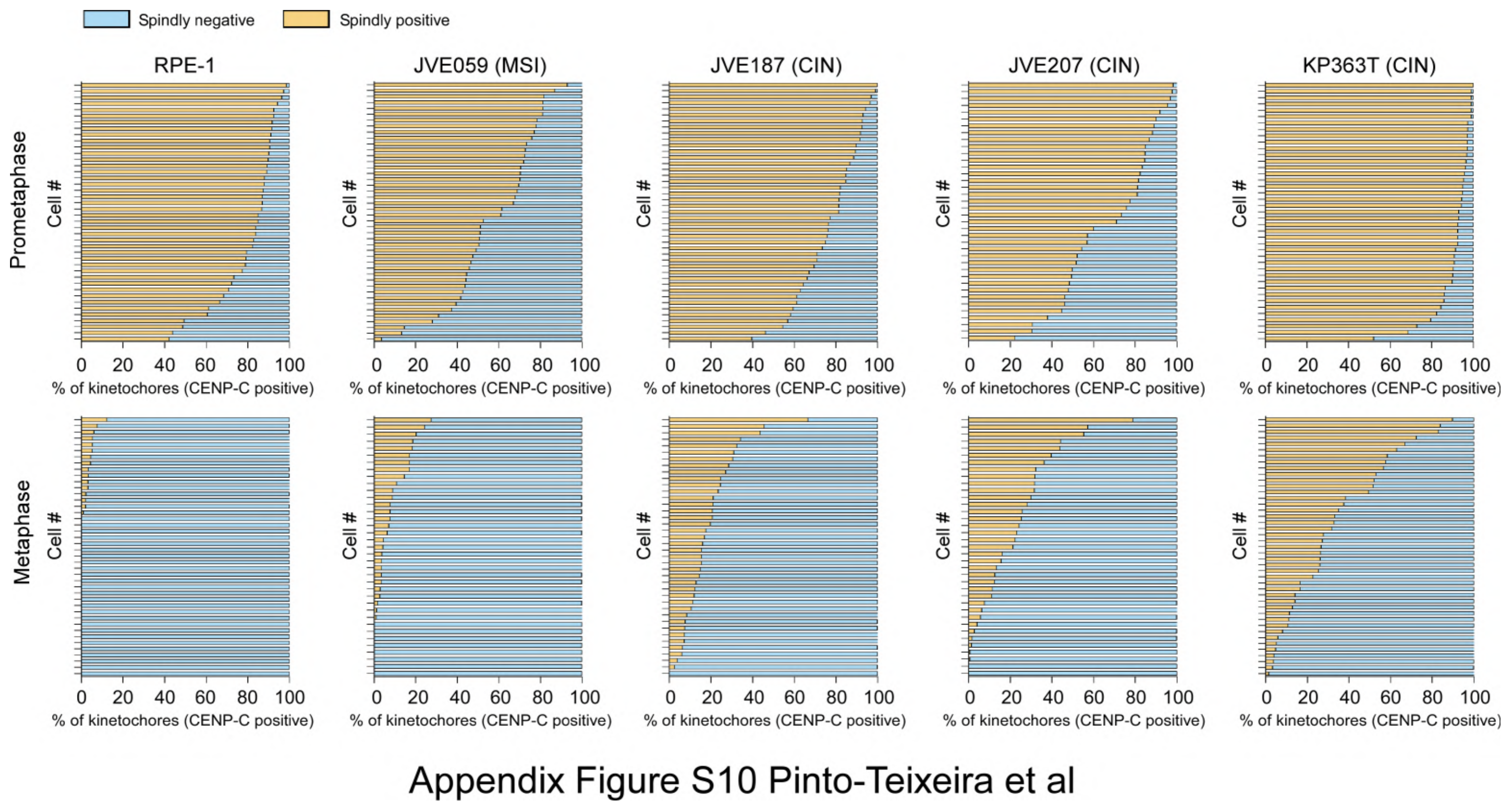


**Appendix Figure S10. Spindly is retained at metaphase kinetochores of CIN+ PCCCs. Related to Figure 9.** Histograms depicting the percentage of prometaphase and metaphase kinetochores per cell that are positive (orange bars) or negative (blue bars) for Spindly in RPE-1 cells and in the PCCCs JVE059 (MSI), JVE187 (CIN), JVE207 (CIN) and KP363T (CIN), as presented in Fig. 9A. CENP-C was used as kinetochore reference. Each bar represents one individual cell; n=2 independent experiments.

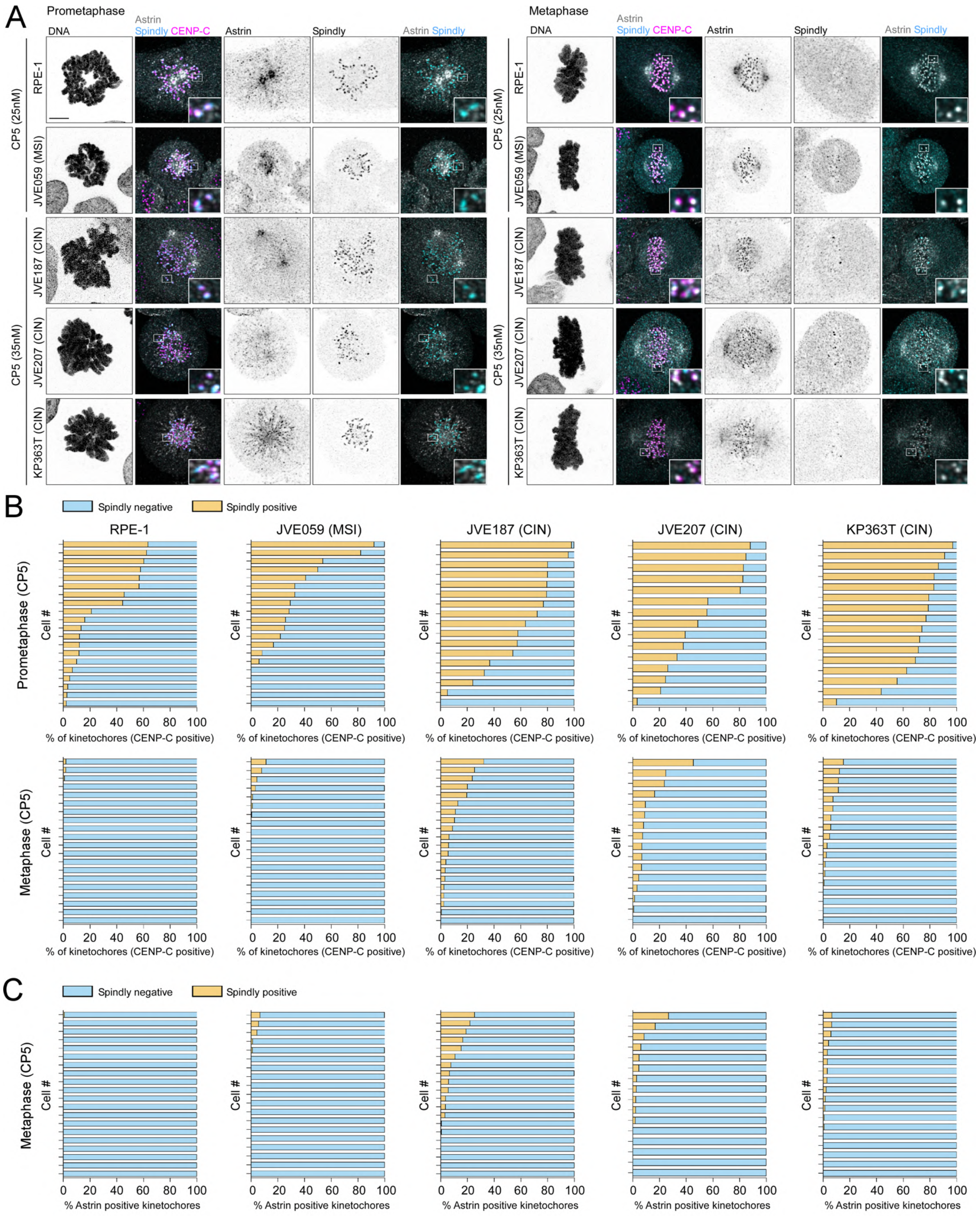


Appendix Figure S11 Pinto-Teixeira et al

**Appendix Figure S11. Spindly is retained at metaphase kinetochores of CIN$^{+}$ PCCCs in an MPS1-dependent manner. Related to Figure 9.**

**(A)** Representative immunofluorescence images of Spindly and Astrin localization at prometaphase and metaphase kinetochores in RPE-1 cells and PCCCs JVE059 (MSI), JVE187 (CIN), JVE207 (CIN) and KP363T (CIN) treated with suboptimal concentrations of the MPS1 inhibitor CP5. CENP-C was used as kinetochore marker. Insets depict magnifications of selected kinetochores.

**(B)** Histograms depicting the percentage of CENP-C-positive kinetochores per cell at prometaphase and metaphase that are also positive (orange bars) or negative (blue bars) for Spindly in the cell lines and conditions indicated in **(A)**. Each bar represents one individual cell; n=2 independent experiments.

**(C)** Histograms depicting the percentage of Astrin positive kinetochores per cell at metaphase that are also positive (orange bars) or negative (blue bars) for Spindly in the cell lines and conditions indicated in (A). Each bar represents one individual cell; n=2 independent experiments.

Data information: data in **(B)** and **(C)** are plotted as fractions of the total. Scale bar: 5µm.

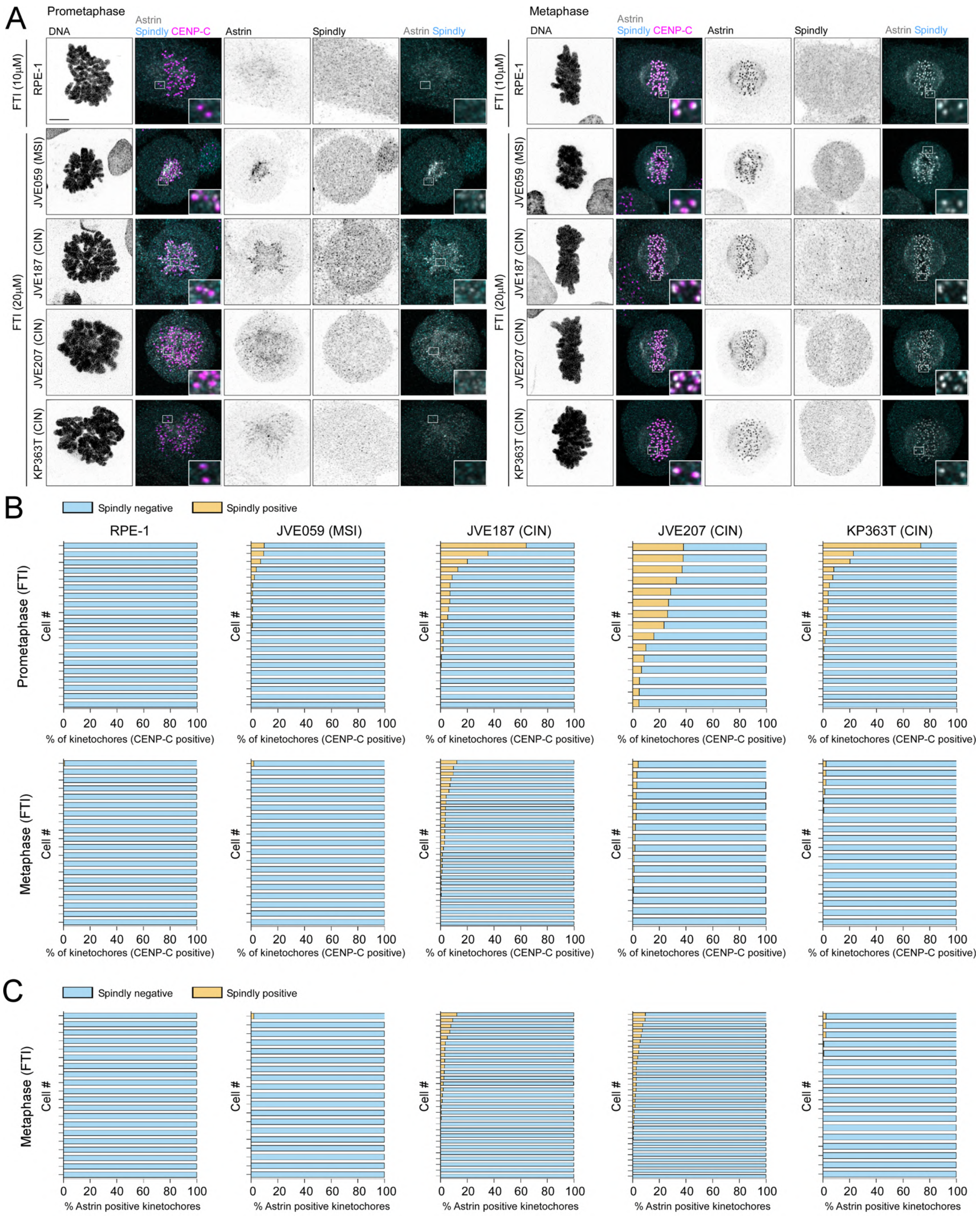


Appendix Figure S12 Pinto-Teixeira et al

**Appendix Figure S12. Blocking fibrous corona assembly with FTI-277 suppressed retention of Spindly at metaphase kinetochores of PCCCs. Related to Figure 9.**

(A) Representative immunofluorescence images of Spindly and Astrin localization at prometaphase and metaphase kinetochores in RPE-1 cells and PCCCs JVE059 (MSI), JVE187 (CIN), JVE207 (CIN) and KP363T (CIN) treated with the farnesyl transferase inhibitor FTI-277. CENP-C was used as kinetochore marker. Insets depict magnifications of selected kinetochores.

**(B)** Histograms depicting the percentage of CENP-C-positive kinetochores per cell at prometaphase and metaphase that are also positive (orange bars) or negative (blue bars) for Spindly in the cell lines and conditions indicated in **(A)**. Each bar represents one individual cell; n=2 independent experiments.

**(C)** Histograms depicting the percentage of Astrin positive kinetochores per cell at metaphase that are also positive (orange bars) or negative (blue bars) for Spindly in the cell lines and conditions indicated in **(A)**. Each bar represents one individual cell; n=2 independent experiments.

Data information: data in **(B)** and **(C)** are plotted as fractions of the total. Scale bar: 5μm.

### Movies Legends

**Movie EV1. Chromosome segregation in RPE-1 cells** (related to Fig. 2C,D). Time-lapse imaging of RPE-1 cells expressing EGFP–αTubulin and H2B–RFP. The merged EGFP–α-tubulin (green) and H2B–RFP (magenta) channels are shown on the left; the H2B–RFP channel is shown as inverted greyscale on the right. Images were acquired every 1 min. Time 0 min corresponds to anaphase onset.

**Movie EV2. Chromosome segregation in JVE059 (MSI) cells** (related to Fig. 2C,D). Time-lapse imaging of JVE059 cells expressing EGFP–αTubulin and H2B–RFP. The merged EGFP–αTubulin (green) and H2B–RFP (magenta) channels are shown on the left; the H2B–RFP channel is shown as inverted greyscale on the right. Images were acquired every 1 min. Time 0 min corresponds to anaphase onset.

**Movie EV3. Chromosome segregation in JVE187 (CIN) cells** (related to Fig. 2C,D). Time-lapse imaging of JVE187 cells expressing EGFP–αTubulin and H2B–RFP. The merged EGFP–αTubulin (green) and H2B–RFP (magenta) channels are shown on the left; the H2B–RFP channel is shown as inverted greyscale on the right. Images were acquired every 2 min. Time 0 min corresponds to anaphase onset.

**Movie EV4. Chromosome segregation in JVE187 (CIN) cells** (related to Fig. 2C,D). Time-lapse imaging of JVE187 cells expressing EGFP–αTubulin and H2B–RFP. The merged EGFP–αTubulin (green) and H2B–RFP (magenta) channels are shown on the left; the H2B–RFP channel is shown as inverted greyscale on the right. Images were acquired every 1 min. Time 0 min corresponds to anaphase onset.

**Movie EV5. Chromosome segregation in JVE207 (CIN) cells** (related to Fig. 2C,D). Time-lapse imaging of JVE207 cells expressing EGFP–αTubulin and H2B–RFP. The merged EGFP–αTubulin (green) and H2B–RFP (magenta) channels are shown on the left; the H2B–RFP channel is shown as inverted greyscale on the right. Images were acquired every 1 min. Time 0 min corresponds to anaphase onset.

**Movie EV6. Chromosome segregation in JVE207 (CIN) cells** (related to Fig. 2C,D). Time-lapse imaging of JVE207 cells expressing EGFP–αTubulin and H2B–RFP. The merged EGFP–αTubulin (green) and H2B–RFP (magenta) channels are shown on the left; the H2B–RFP channel is shown as inverted greyscale on the right. Images were acquired every 1 min. Time 0 min corresponds to anaphase onset.

**Movie EV7. Chromosome segregation in KP363T (CIN) cells** (related to Fig. 2C,D). Time-lapse imaging of KP363T cells expressing EGFP–αTubulin and H2B–RFP. The merged EGFP–αTubulin (green) and H2B–RFP (magenta) channels are shown on the left; the H2B–RFP channel is shown as inverted greyscale on the right. Images were acquired every 1 min. Time 0 min corresponds to anaphase onset.

**Movie EV8. Chromosome segregation in KP363T (CIN) cells** (related to Fig. 2C,D). Time-lapse imaging of KP363T cells expressing EGFP–αTubulin and H2B–RFP. The merged EGFP–αTubulin (green) and H2B–RFP (magenta) channels are shown on the left; the H2B–RFP channel is shown as inverted greyscale on the right. Images were acquired every 1 min. Time 0 min corresponds to anaphase onset.

**Movie EV9. Mitotic progression of RPE-1 cells** (related to Appendix Fig. S2B,C). Time-lapse imaging of RPE-1 cells expressing EGFP–αTubulin and H2B–RFP. The merged EGFP–αTubulin (green) and H2B–RFP (magenta) channels are shown on the left; the H2B–RFP channel is shown as inverted greyscale on the right. Images were acquired every 1 min. Time 0 min corresponds to nuclear envelope breakdown (NEB).

**Movie EV10. Mitotic progression of JVE059 (MSI) cells** (related to Appendix Fig. S2B,C). Time-lapse imaging of JVE059 cells expressing EGFP–αTubulin and H2B–RFP. The merged EGFP–αTubulin (green) and H2B–RFP (magenta) channels are shown on the left; the H2B–RFP channel is shown as inverted greyscale on the right. Images were acquired every 1 min. Time 0 min corresponds to nuclear envelope breakdown (NEB).

**Movie EV11. Mitotic progression of JVE187 (CIN) cells** (related to Appendix Fig. S2B,C). Time-lapse imaging of JVE187 cells expressing EGFP–αTubulin and H2B–RFP. The merged EGFP–αTubulin (green) and H2B–RFP (magenta) channels are shown on the left; the H2B–RFP channel is shown as inverted greyscale on the right. Images were acquired every 1 min. Time 0 min corresponds to nuclear envelope breakdown (NEB).

**Movie EV12. Mitotic progression of JVE207 (CIN) cells** (related to Appendix Fig. S2B,C). Time-lapse imaging of JVE207 cells expressing EGFP–αTubulin and H2B–RFP. The merged EGFP–αTubulin (green) and H2B–RFP (magenta) channels are shown on the left; the H2B–RFP channel is shown as inverted greyscale on the right. Images were acquired every 1 min. Time 0 min corresponds to nuclear envelope breakdown (NEB).

**Movie EV13. Mitotic progression of KP363T (CIN) cells** (related to Appendix Fig. S2B,C). Time-lapse imaging of KP363T cells expressing EGFP–αTubulin and H2B–RFP. The merged EGFP–αTubulin (green) and H2B–RFP (magenta) channels are shown on the left; the H2B–RFP channel is shown as inverted greyscale on the right. Images were acquired every 1 min. Time 0 min corresponds to nuclear envelope breakdown (NEB).

**Movie EV14. Chromosome segregation in parental RPE-1 cells** (related to Fig. 4C,D). Time-lapse imaging of parental RPE-1 cells expressing H2B–RFP. The H2B–RFP channel is shown in magenta on the left and as inverted greyscale on the right. Images were acquired every 1 min. Time 0 min corresponds to anaphase onset.

**Movie EV15. Chromosome segregation in RPE-1 cells constitutively overexpressing MPS1 under the CMV promoter ($P_{CMV}$)** (related to Fig. 4C,D). Time-lapse imaging of RPE-1 cells expressing EGFP–MPS1 and H2B–RFP.

The merged EGFP–MPS1 (green) and H2B–RFP (magenta) channels are shown on the left; the H2B–RFP channel is shown as inverted greyscale on the right. Images were acquired every 1 min. Time 0 min corresponds to anaphase onset.

**Movie EV16. Chromosome segregation in RPE-1 cells carrying a doxycycline-inducible MPS1 expression system in the absence of doxycycline** (related to Fig. 4C,D). Time-lapse imaging of RPE-1 cells carrying a doxycycline-inducible EGFP–MPS1 expression system ($P_{tet}$) and expressing H2B–RFP, cultured in the absence of doxycycline (−DOX). The EGFP–MPS1 and H2B–RFP channels are shown in green and magenta, respectively, on the left; the H2B–RFP channel is shown as inverted greyscale on the right. Images were acquired every 1 min. Time 0 min corresponds to anaphase onset.

**Movie EV17. Chromosome segregation in RPE-1 cells overexpressing doxycycline-inducible EGFP–MPS1** (related to Fig. 4C,D). Time-lapse imaging of RPE-1 cells carrying a doxycycline-inducible EGFP–MPS1 expression system ($P_{tet}$) and expressing H2B–RFP, cultured in the presence of doxycycline (+DOX). The EGFP–MPS1 and H2B–RFP channels are shown in green and magenta, respectively, on the left; the H2B–RFP channel is shown as inverted greyscale on the right. Images were acquired every 1 min. Time 0 min corresponds to anaphase onset.

**Movie EV18. Chromosome segregation in parental JVE059 (MSI) cells** (related to Appendix Fig. S4B,C). Time-lapse imaging of parental JVE059 cells expressing H2B–RFP. The H2B–RFP channel is shown in magenta on the left and as inverted greyscale on the right. Images were acquired every 1 min. Time 0 min corresponds to anaphase onset.

**Movie EV19. Chromosome segregation in JVE059 (MSI) cells constitutively overexpressing MPS1 under the CMV promoter ($P_{CMV}$)** (related to Appendix Fig. S4B,C). Time-lapse imaging of JVE059 cells expressing EGFP–MPS1 and H2B–RFP. The merged EGFP–MPS1 (green) and H2B–RFP (magenta) channels are shown on the left; the H2B–RFP channel is shown as inverted greyscale on the right. Images were acquired every 1 min. Time 0 min corresponds to anaphase onset.

**Movie EV20. Chromosome segregation in JVE059 (MSI) cells carrying a doxycycline-inducible MPS1 expression system in the absence of doxycycline** (related to Appendix Fig. S4B,C). Time-lapse imaging of JVE059 cells carrying a doxycycline-inducible EGFP–MPS1 expression system ($P_{tet}$) and expressing H2B–RFP, cultured in the absence of doxycycline (−DOX). The EGFP–MPS1 and H2B–RFP channels are shown in green and magenta, respectively, on the left; the H2B–RFP channel is shown as inverted greyscale on the right. Images were acquired every 1 min. Time 0 min corresponds to anaphase onset.

**Movie EV21. Chromosome segregation in JVE059 (MSI) cells overexpressing doxycycline-inducible EGFP–MPS1** (related to Appendix Fig. S4B,C). Time-lapse imaging of JVE059 cells carrying a doxycycline-inducible EGFP–MPS1 expression system ($P_{tet}$) and expressing H2B–RFP, cultured in the presence of doxycycline (+DOX). The EGFP–MPS1 and H2B–RFP channels are shown in green and magenta, respectively, on the left; the H2B–RFP channel is shown as inverted greyscale on the right. Images were acquired every 1 min. Time 0 min corresponds to anaphase onset.

**Movie EV22. Micronuclei formation in dividing RPE-1 cells constitutively overexpressing MPS1 under the CMV promoter ($P_{CMV}$)** (related to Fig. 4F,G). Time-lapse imaging of RPE-1 cells expressing EGFP–MPS1 and H2B–RFP. The merged EGFP–MPS1 (green) and H2B–RFP (magenta) channels are shown on the left; the H2B–RFP channel is shown as inverted greyscale on the right. Images were acquired every 5 min. Time 0 min corresponds to anaphase onset.

**Movie EV23. Three-dimensional STED reconstruction of an amphitelic kinetochore–microtubule attachment in a metaphase RPE-1 cell overexpressing MPS1** (related to Fig. EV4D). STED z-stacks were acquired through the kinetochore–microtubule attachment interface and rendered in three dimensions using Huygens Professional v25.10 (Scientific Volume Imaging, The Netherlands). Microtubules (tubulin; yellow), Spindly (blue) and centromeres/inner-kinetochores (ACA; red) are shown. The orientation cube in the bottom right rotates synchronously with the three-dimensional reconstruction to indicate the viewing angle.

**Movie EV24. Three-dimensional STED reconstruction of an amphitelic kinetochore–microtubule attachment in a metaphase RPE-1 cell overexpressing MPS1** (related to Fig. EV4D). STED z-stacks were acquired through the kinetochore–microtubule attachment interface and rendered in three dimensions using Huygens Professional v25.10 (Scientific Volume Imaging, The Netherlands). Microtubules (tubulin; yellow), Spindly (blue) and centromeres/inner-kinetochores (ACA; red) are shown. The orientation cube in the bottom right rotates synchronously with the three-dimensional reconstruction to indicate the viewing angle.

**Movie EV25. Three-dimensional STED reconstruction of a merotelic kinetochore–microtubule attachment in a metaphase RPE-1 cell overexpressing MPS1** (related to Fig. EV4E). STED z-stacks were acquired through the kinetochore–microtubule attachment interface and rendered in three dimensions using Huygens Professional v25.10 (Scientific Volume Imaging, The Netherlands). Microtubules (tubulin; yellow), Spindly (blue) and centromeres/inner-kinetochores (ACA; red) are shown. The orientation cube in the bottom right rotates synchronously with the three-dimensional reconstruction to indicate the viewing angle.

**Movie EV26. Three-dimensional STED reconstruction of a merotelic kinetochore–microtubule attachment in a metaphase RPE-1 cell overexpressing MPS1** (related to Fig. EV4E). STED z-stacks were acquired through the kinetochore–microtubule attachment interface and rendered in three dimensions using Huygens Professional v25.10 (Scientific Volume Imaging, The Netherlands). Microtubules (tubulin; yellow), Spindly (blue) and centromeres/inner-kinetochores (ACA; red) are shown. The orientation cube in the bottom right rotates synchronously with the three-dimensional reconstruction to indicate the viewing angle.

**Movie EV27. Three-dimensional STED reconstruction of a merotelic kinetochore–microtubule attachment on an anaphase lagging chromosome in an RPE-1 cell overexpressing MPS1** (related to Fig. EV4F). STED z-stacks were acquired through the kinetochore–microtubule attachment interface and rendered in three dimensions using Huygens Professional v25.10 (Scientific Volume Imaging, The Netherlands). Microtubules (tubulin; yellow), Spindly (blue) and centromere/inner-kinetochore (ACA; red) are shown. The orientation cube in the bottom right rotates synchronously with the three-dimensional reconstruction to indicate the viewing angle.

**Movie EV28. Three-dimensional STED reconstruction of a merotelic kinetochore–microtubule attachment on an anaphase lagging chromosome in an RPE-1 cell overexpressing MPS1** (related to Fig. EV4F). STED z-stacks were acquired through the kinetochore–microtubule attachment interface and rendered in three dimensions using Huygens Professional v25.10 (Scientific Volume Imaging, The Netherlands). Microtubules (tubulin; yellow), Spindly (blue) and centromere/inner-kinetochore (ACA; red) are shown. The orientation cube in the bottom right rotates synchronously with the three-dimensional reconstruction to indicate the viewing angle.

**Movie EV29. Chromosome segregation in RPE-1 cells carrying a doxycycline-inducible MPS1 expression system in the absence of doxycycline** (related to Fig. 8D,E). Time-lapse imaging of RPE-1 cells carrying a doxycycline-inducible EGFP–MPS1 expression system ($P_{tet}$) and expressing H2B–RFP, cultured in the absence of doxycycline (−DOX). The EGFP–MPS1 and H2B–RFP channels are shown in green and magenta, respectively, on the left; the H2B–RFP channel is shown as inverted greyscale on the right. Images were acquired every 1 min. Time 0 min corresponds to anaphase onset.

**Movie EV30. Chromosome segregation in RPE-1 cells overexpressing doxycycline-inducible EGFP–MPS1** (related to Fig. 8D,E). Time-lapse imaging of RPE-1 cells carrying a doxycycline-inducible EGFP–MPS1 expression system ($P_{tet}$) and expressing H2B–RFP, cultured in the presence of doxycycline (+DOX). The EGFP–MPS1 and H2B–RFP channels are shown in green and magenta, respectively, on the left; the H2B–RFP channel is shown as inverted greyscale on the right. Images were acquired every 1 min. Time 0 min corresponds to anaphase onset.

**Movie EV31. Chromosome segregation in RPE-1 cells overexpressing doxycycline-inducible EGFP–MPS1 and treated with CP5** (related to Fig. 8D,E). Time-lapse imaging of RPE-1 cells carrying a doxycycline-inducible EGFP–MPS1 expression system ($P_{tet}$) and expressing H2B–RFP, cultured in the presence of doxycycline (+DOX) and a suboptimal concentration of the MPS1 inhibitor CP5 (+CP5, 25 nM). The EGFP–MPS1 and H2B–RFP channels are shown in green and magenta, respectively, on the left; the H2B–RFP channel is shown as inverted greyscale on the right. Images were acquired every 1 min. Time 0 min corresponds to anaphase onset.

**Movie EV32. Chromosome segregation in RPE-1 cells overexpressing doxycycline-inducible EGFP–MPS1 and treated with FTI-277** (related to Fig. 8D,E). Time-lapse imaging of RPE-1 cells carrying a doxycycline-inducible EGFP–MPS1 expression system ($P_{tet}$) and expressing H2B–RFP, cultured in the presence of doxycycline (+DOX) and the farnesyl-transferase inhibitor FTI-277 (+FTI, 10 μM). The EGFP–MPS1 and H2B–RFP channels are shown in green and magenta, respectively, on the left; the H2B–RFP channel is shown as inverted greyscale on the right. Images were acquired every 1 min. Time 0 min corresponds to anaphase onset.

**Movie EV33. Mitotic progression and chromosome segregation in RPE-1 cells carrying a doxycycline-inducible MPS1 expression system in the absence of doxycycline and treated with non-targeting siRNA** (related to Appendix Fig. S9A-C). Time-lapse imaging of RPE-1 cells carrying a doxycycline-inducible EGFP–MPS1 expression system ($P_{tet}$) and expressing H2B–RFP, cultured in the absence of doxycycline (−DOX) and treated with non-targeting small interfering RNA (siNT). The EGFP–MPS1 and H2B–RFP channels are shown in green and magenta, respectively, on the left; the H2B–RFP channel is shown as inverted greyscale on the right. Images were acquired every 2 min. Time 0 min corresponds to nuclear envelope breakdown.

**Movie EV34. Mitotic progression and chromosome segregation in RPE-1 cells overexpressing doxycycline-inducible EGFP–MPS1 and treated with non-targeting siRNA** (related to Appendix Fig. S9A-C). Time-lapse imaging of RPE-1 cells carrying a doxycycline-inducible EGFP–MPS1 expression system ($P_{tet}$) and expressing H2B–RFP, cultured in the presence of doxycycline (+DOX) and treated with non-targeting small interfering RNA (siNT). The EGFP–MPS1 and H2B–RFP channels are shown in green and magenta, respectively, on the left; the H2B–RFP channel is shown as inverted greyscale on the right. Images were acquired every 2 min. Time 0 min corresponds to nuclear envelope breakdown.

**Movie EV35. Mitotic progression and chromosome segregation in RPE-1 cells overexpressing doxycycline-inducible EGFP–MPS1 and depleted of Spindly** (related to Appendix Fig. S9A-C). Time-lapse imaging of RPE-1 cells carrying a doxycycline-inducible EGFP–MPS1 expression system ($P_{tet}$) and expressing H2B–RFP, cultured in the presence of doxycycline (+DOX) and treated with small interfering RNA targeting *Spindly* (siSpindly). The EGFP–MPS1 and H2B–RFP channels are shown in green and magenta, respectively, on the left; the H2B–RFP channel is shown as inverted greyscale on the right. Images were acquired every 2 min. Time 0 min corresponds to nuclear envelope breakdown.
